# DNA damage at Lamina-Associated Domains triggers nuclear envelope reorganization and chromatin detachment to prevent nuclear envelope blebbing and genome instability

**DOI:** 10.64898/2026.10.25.761546

**Authors:** Diana Rubio-Contreras, Sylvain Audibert, Joanna Fernandez, Ruxandra A. Lambuta, Nikoleta Pateraki, Karen Meaburn, Jack Wootton, Raquel Carreira, Nishita Parnandi, Ramon Gonzalez-Mendez, Simon J. Boulton, Zuza Kozik, Stamatis Papathanasiou, Evi Soutoglou

**Author notes:** Equal contribution (Diana Rubio-Contreras, Sylvain Audibert). Equal contribution (Joanna Fernandez, Ruxandra A. Lambuta).

## Abstract

The nuclear periphery anchors large, transcriptionally silent chromatin domains to the nuclear envelope and plays a central role in maintaining genome stability, acting as a dynamic hub for DNA repair. Nevertheless, how cells process DNA damage within lamina-associated domains (LADs), which reside in this environment, remains poorly understood, despite their relevance to nuclear architecture, aging, and senescence. By employing systems for temporally controlled and spatially precise induction of DNA double-strand breaks (DSBs) within LADs, we show that DSBs within LADs trigger remodeling of the nuclear lamina and the dynamic detachment of damaged LADs from the nuclear envelope, resulting in their relocalization to the nuclear interior. This process is orchestrated by the ATM–γH2AX–MDC1–RNF168–53BP1 axis and requires the linker of nucleoskeleton and cytoskeleton (LINC) complex and cytoskeletal components while leaving DNA repair pathway choice unchanged. Persistent tethering of damaged LADs at the nuclear periphery delays the DNA damage response and repair, elevates genomic instability, and drives chromatin extrusion through nuclear blebs directly linking failed repair to dysregulation of nuclear integrity. These findings reveal a protective mechanism in which ATM-driven LAD dynamics and nuclear envelope remodeling might relieve mechanical stress, facilitate repair, and safeguard both genome stability and nuclear architecture.

---

The eukaryotic nucleus is a highly compartmentalized organelle that safeguards the genome while orchestrating essential processes such as transcription, replication, and DNA repair^1^. Chromatin is non-randomly distributed within the nuclear space, with transcriptionally silent, gene-poor regions often anchored at the nuclear periphery in lamina-associated domains (LADs), while gene-rich, active regions are more centrally located^2^. This three-dimensional organization is further defined by the formation of topologically associating domains (TADs) and chromatin loops, which facilitate long-range regulatory interactions^1^. Collectively, this hierarchical nuclear structure shapes cellular identity, function, and fate.

In addition to its role in gene regulation, nuclear architecture is becoming increasingly recognized as a critical determinant of genome stability^3^. Recent studies have uncovered that the spatial positioning of DNA double-strand breaks (DSBs) within the nucleus influences their recognition, mobility, and repair^3^. The nuclear periphery in particular has emerged as a specialized subnuclear compartment that facilitates the resolution of persistent DSBs^3^. In mammalian cells, two major pathways have been identified: one involves the linker of nucleoskeleton and cytoskeleton (LINC) complex and relies on actin- and myosin-dependent transport to translocate lesions from the interior to the periphery and particularly the nuclear pores^4–8^, and the other involves the formation of DSB-capturing nuclear envelope tubules (dsbNETs)^9^, which are transient invaginations of the nuclear envelope that bring the envelope itself closer to the break site. These tubules, composed of the inner and outer nuclear membranes and the nuclear lamina, are formed through coordinated action of cytoplasmic microtubules, LINC complexes, nuclear pore complexes (NPCs), and plus-end-directed kinesins. By reducing the spatial distance to repair hubs at the nuclear envelope, dsbNETs create a dynamic environment for damage sensing and resolution^9^.

While the nuclear periphery exhibits dynamic remodelling to engage with damaged chromatin within the nuclear interior, the response to lesions arising within LADs, which are constitutively tethered to the nuclear envelope, is unknown. This gap is particularly striking given that LADs encompass nearly 40% of the mammalian genome and serve as key architectural anchors that maintain nuclear integrity^10,11^. Importantly, disruptions in LAD–nuclear envelope interactions are a hallmark of senescence and aging^12^, suggesting that compromised DNA repair within these domains may have far-reaching consequences for genome stability and age-associated decline. Investigating how DNA lesions are sensed and resolved within LADs is therefore essential to understanding nuclear homeostasis under both physiological and pathological conditions.

By engineering systems that allow for precise temporal control of DSB formation within LADs, we uncovered that DSBs within LADs activate ATM signaling, which facilitates the release of damaged chromatin from the periphery and movement into the nuclear interior. LAD dynamics rely on the ATM–γH2AX–MDC1–RNF168–53BP1 axis and the LINC complex, which signals to the cytoskeleton and acts to buffer mechanical stress and maintain nuclear envelope integrity during repair. In contrast, artificially maintaining LAD anchoring impairs the DNA damage response and heightens genomic instability. Furthermore, failure to repair LAD-specific DSBs leads to mitotically driven lamina deformations in the form of nuclear blebs that persist in the following interphase. These observations reveal a protective mechanism in which ATM-dependent chromatin mobilization mitigates structural strain on the nuclear envelope while supporting genome maintenance, and identify a previously unrecognized pathway regulating LAD-nuclear lamina interconnection in response to genotoxic stress.

## Results

### Generation of an experimental system for site-specific induction of DSBs at LADs

Seeking to uncover how DNA damage within LADs affects nuclear organization and genomic stability, we established a novel experimental system enabling the precise and temporal induction of DSBs within LADs. This system leverages DamID technology^13^, a method previously employed for mapping chromatin domains proximal to the nuclear lamina. We utilized a fusion of the bacterial Adenine methylase Dam with lamin B1 (LaminB1-Dam), resulting in the targeted methylation of LAD-associated DNA (Fig. 1a). For this, we have used HT1080 fibrosarcoma cells stably expressing LaminB1-Dam, which is tightly controlled by its fusion to a destabilization domain (DD) (Fig. 1b). Addition of a “protein shield” (Shield1) transiently stabilizes the LaminB1-Dam and leads to DNA methylation of LADs (Fig. 1b). The fidelity of this targeting was confirmed by the peripheral localization of a GFP-tagged m⁶A-Tracer protein^14^, which specifically binds Adenine-methylated DNA (m⁶A) (Fig. 1a, b). To induce DSBs at these defined genomic regions, we expressed the DpnI restriction endonuclease, which specifically cleaves m^6^A (Fig. 1c). Temporal control over DSB induction and subsequent repair was achieved by engineering DpnI to include a modified estrogen receptor (ER*), which triggers nuclear translocation of DpnI upon addition of tamoxifen (4OHT), and an Auxin-Inducible Degron (AID) system^15^, which induces DpnI degradation in the presence of Auxin (Fig. 1c). Efficient DpnI-dependent DSB induction at methylated LADs was observed, with 20 hours treatment with Shield1 and 2 hours treatment with 4OHT added in the last 2 hours of Shield1 (these experimental conditions were used in all experiments), leading to a significant increase in cells displaying peripheral γH2AX and 53BP1 foci (Fig. 1c and Extended Data Fig. 1a). This was accompanied by ATM autophosphorylation (pATM), a hallmark of DNA damage signaling (Extended Data Fig. 1a).

**Figure 1:**
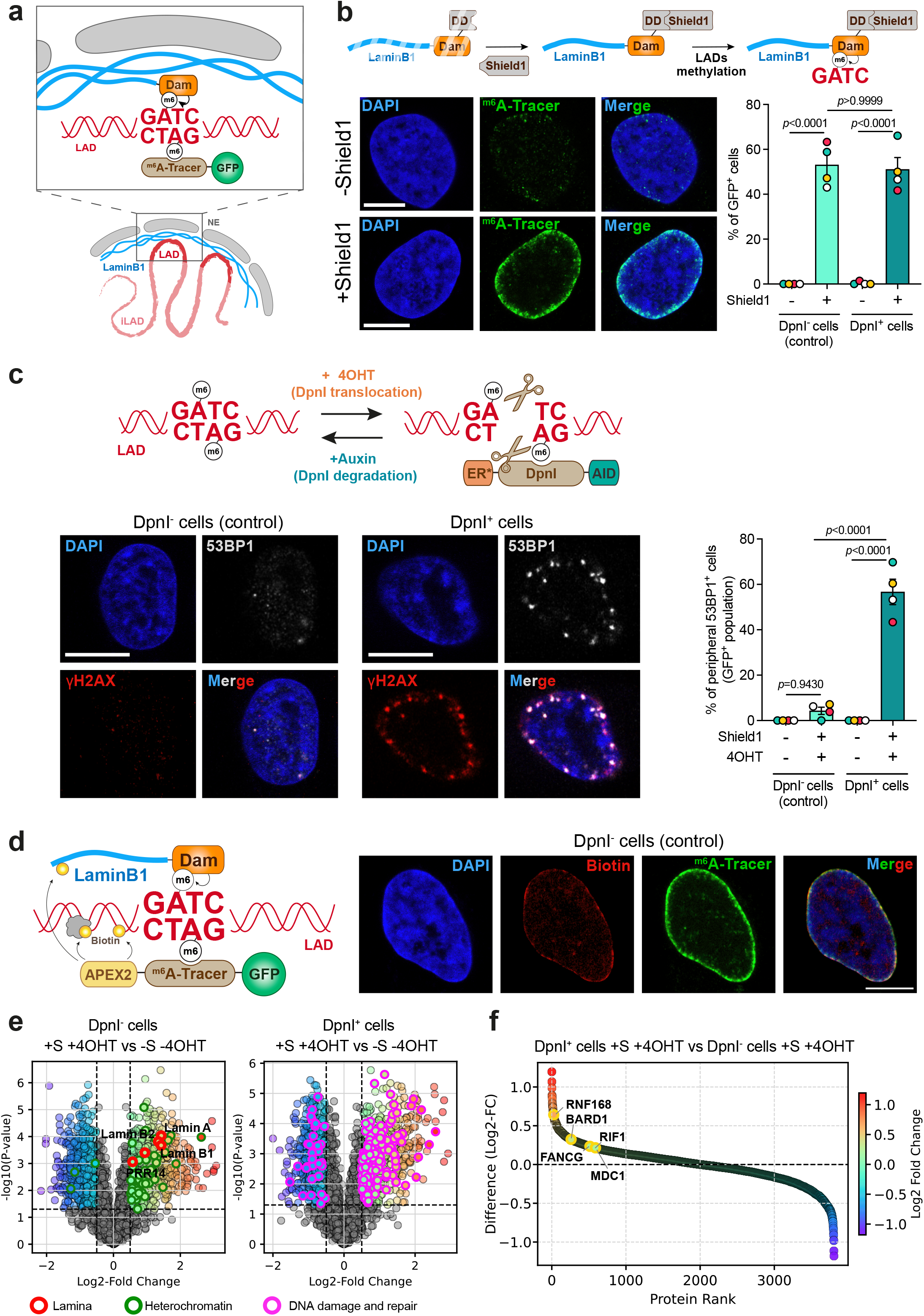
A novel experimental system to induce DSBs at LADs. a) Schematic representation of the experimental system to visualize LADs. Lamin B1 is fused to Dam methylase which induces methylation at Adenines in GATC (m^6^A). m^6^A can be visualised by m^6^A-Tracer fused to GFP. iLADs are regions of the genome which are not associated with lamina. b) Schematic representation of the experimental system used to stabilize lamin B1 fused to Dam and induce m^6^A methylation. *Below left*, representative image of the localization of the m^6^A-Tracer (green) in the absence or presence of Shield1 for 20 h. The DNA is stained with DAPI (blue). Scale bar 10µm. *Below right,* quantification of GFP-positive cells in the presence (DpnI⁺) or absence (DpnI⁻, control) of DpnI expression, and with or without Shield1 treatment. Data are presented as mean ± SEM from 4 independent biological replicates. Statistical comparisons were performed using one-way ANOVA with Bonferroni correction for multiple comparisons. c) Schematic representation of the experimental system to induce DSBs using DpnI. 4OHT, added in the last 2 h of Shield1, leads to the nuclear translocation of DpnI protein and IAA treatment leads to its degradation. *Below left*, representative confocal images of 53BP1 (white) and γH2AX (red) in cells expressing or not DpnI in the absence and presence of Shield1 and 4OHT. The DNA is stained with DAPI (blue). Scale bar 10µm. *Right*, quantification of 53BP1-positive cells in the GFP^+^ population in cells expressing or not DpnI in the absence and presence of Shield1 and 4OHT. Data are presented as mean ± SEM from 4 independent biological replicates. Statistical comparisons were performed using one-way ANOVA with Bonferroni correction for multiple comparisons. d) *Left*, schematic representation of the experimental system to target APEX2 at LADs. APEX2 fused to m^6^A-Tracer binds to m^6^A and induces biotinylation of proteins at the vicinity. *Right*, representative confocal image of APEX2-induced biotinylation (red) and m^6^A-Tracer (green) at the nuclear periphery. The DNA is stained with DAPI (blue). Scale bar 10µm. e) Volcano plots showing differential protein abundance under DpnI expression and treatment conditions. *Left*, comparison of cells not expressing DpnI, with or without Shield1 (S) and 4OHT treatment, revealing significant enrichment of nuclear lamina proteins (red) and heterochromatin-associated proteins (green). *Right*, comparison of DpnI expressing cells, with or without S and 4OHT treatment, highlighting proteins upregulated in response to DpnI-induced DNA damage (magenta). Significantly regulated proteins were defined by two-sample *t*-test (p < 0.05) and absolute log₂ fold change > 0.5. f) DpnI induces selective upregulation of DNA repair factors. Rank-ordered plot of protein abundance differences between cells expressing or not DpnI and treated with Shield1 and 4OHT, with proteins color-coded by log₂ fold change. Some DNA damage response factors are highlighted and annotated, revealing selective upregulation consistent with DpnI-induced DNA repair activation.

To further validate the precision of our targeting, we sought to determine the proteomic landscape of LAD chromatin and its dynamic modulation following DNA damage. To this end, we integrated DamID with proximity-dependent biotinylation (BioID)^16^. This involved generating an APEX2-m⁶A-Tracer fusion to direct the promiscuous biotin ligase APEX2 to LADs (Fig. 1d), enabling the spatially restricted biotinylation of neighboring proteins (Fig. 1d), which were subsequently analysed using Tandem Mass Tag (TMT) mass spectrometry.

Our analysis involved comparing protein enrichment under two conditions: one where APEX2 was targeted to LADs using LaminB1-Dam stabilization (+Shield1; +S), and a control where APEX2 was diffusely distributed throughout the cell (-Shield1; -S) conducted in both the presence (DpnI⁺ cells; Supplementary Table 1) and absence (DpnI⁻ cells, control; Supplementary Table 1) of the DpnI nuclease (Fig. 1e). We performed the experiment in biological triplicates and quantified a total of 3,807 proteins across all conditions. Principal Component Analysis (PCA) of the quantitative proteomic data revealed tight clustering of biological replicates, indicative of strong experimental reproducibility (Extended Data Fig. 1b). Furthermore, the PCA analysis demonstrated close clustering of the control cell lines (-S condition, blue and red circles), reflecting shared basal proteome (Extended Data Fig. 1b). Notably, the first principal component (PC1) clearly segregated proteins enriched at the nuclear lamina from those broadly distributed throughout the cell, underscoring a primary source of variance in the dataset that aligns with our experimental design (compare orange and purple, respectively; Extended Data Fig. 1b).

To validate the specificity of our approach, we performed a two-sample *t*-test comparing protein abundance between -S -4OHT and +S +4OHT conditions in cells which do not express DpnI (DpnI⁻). Among the 694 proteins significantly upregulated, several core components of the nuclear lamina—including lamin A/C, lamin B1, lamin B2, and PRR14—were markedly enriched upon APEX2 targeting (Fig. 1e, DpnI⁻ cells). Consistent with the expected compartmental specificity, multiple heterochromatin-associated proteins were also significantly enriched in the lamina fraction (Fig. 1e, DpnI⁻ cells). This pronounced and selective enrichment of canonical nuclear periphery markers confirms that our strategy effectively isolates lamina-associated chromatin from the broader nuclear and cellular compartment. As expected, the same treatments in DpnI⁺ cells led to significant upregulation of 724 proteins, including key components of the DNA damage response (Fig. 1e, DpnI⁺ cells). These findings support the robustness of our system in capturing proteins localized near the nuclear periphery in the absence and presence of DNA damage, thereby enabling high-confidence downstream analyses of chromatin compartmentalization.

Gene Ontology (GO) analysis was performed using Fisher’s exact test on proteins significantly upregulated in each condition. In DpnI⁻ cells (no damage), this revealed strong enrichment for pathways associated with the nuclear periphery, including nuclear envelope and pore organization, heterochromatin formation, and nucleocytoplasmic transport—consistent with the biochemical specificity of our fractionation strategy (Extended Data Fig. 1c). In contrast, DpnI⁺ cells (damaged) showed significant enrichment for GO terms related to DNA damage response (DDR), checkpoint activation, and DSB repair (Extended Data Fig. 1c).

Collectively, this functional enrichment analysis provides independent validation of the specificity and biological relevance of our lamina pull-down strategy. Capture of nuclear envelope structures, chromatin complexes, DDR and DNA repair factors showcases the utility of our approach for dissecting subnuclear architecture and associated molecular activities.

To investigate how lamina-associated chromatin is remodelled following DNA damage, we first assessed whether known DNA repair factors are differentially recruited to this nuclear compartment upon DSB induction. Multiple well-characterized DNA repair proteins—including RNF168, RIF1, MDC1 and BARD1—were significantly enriched in the lamina-associated fraction after damage (Fig. 1f). In line with this, inhibition of ATM resulted in a marked reduction of enrichment of several key DDR factors, including the ones mentioned above (Extended Data Fig. 1d). These results suggest that ATM activity is required for the efficient recruitment or stabilization of repair proteins at LADs. These results are unlikely to reflect changes in the proteome resulting from cell cycle alterations, as no changes in cell cycle distribution were observed during the treatment period (Extended Data Fig. 1e, left).

All together, these data indicate that our system is both specific and robust in targeting LADs and in inducing DSBs within these regions, and in profiling the LAD-recruited machinery responsible for DNA damage sensing and repair.

### DSBs at LADs trigger reorganization of the nuclear envelope and lead to LAD detachment

To explore how DNA damage at LADs affects nuclear lamina organization, we examined the distribution of major lamina components following DSB induction. Interestingly, the relative enrichment of lamin B1 and lamin B2 at the nuclear periphery was decreased in cells where DNA damage was induced (Fig. 2a), suggesting that lamin B1 and lamin B2 localization may be dynamically regulated in response to DNA damage at LADs, or that LADs may detach from the nuclear envelope under these conditions. In contrast, lamin A, which is distributed between the nuclear envelope and the nucleoplasm, remained unchanged (Fig. 2a).

**Figure 2:**
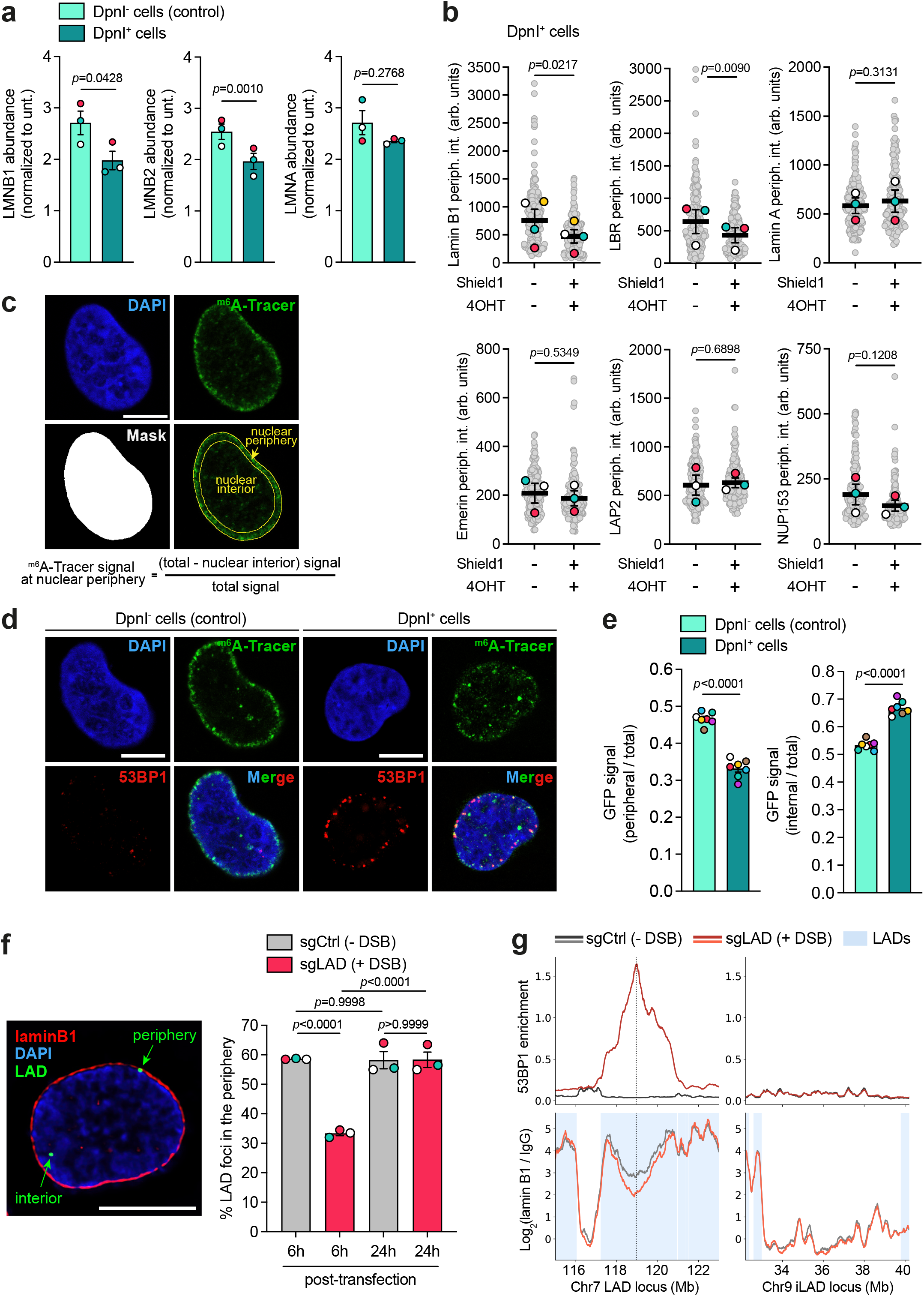
DSBs at LADs lead to nuclear envelope reorganization and LAD detachment. a) Relative enrichment of lamin B1, lamin B2 and lamin A at the nuclear periphery in cells expressing or not DpnI and treated with Shield1 and 4OHT. Quantification was performed using log₂ fold-change values derived from our TMT proteomics dataset. Data are presented as mean ± SEM from 3 independent biological replicates. Statistical comparisons were performed using paired *t-*test. b) Quantification in arbitrary units (linear scale) of lamin B1, LBR, lamin A, emerin, LAP2 and NUP153 at the nuclear periphery in cells expressing DpnI in the absence and presence of Shield1 and 4OHT. Data are presented as mean ± SEM from at least 3 independent biological replicates. Ratio paired *t*-test was used to assess fold-change differences. c) A representative confocal image shows the nuclear localization of the m^6^A-Tracer. Illustration of the method used for quantifying relative enrichment of m^6^A-Tracer. Scale bar 10µm. The relative enrichment of m⁶A-Tracer at the nuclear rim was calculated as the ratio of (total signal − internal signal) / total signal. d) Representative confocal images of m^6^A-Tracer (green) and 53BP1 (red) localization in cells expressing or not DpnI, treated with Shield1 and 4OHT. The DNA is stained with DAPI (blue). Scale bar 10µm. e) Quantification of the relative abundance of m^6^A-Tracer at the nuclear rim (*left*) or nuclear interior (*right*) in cells expressing or not DpnI, treated with Shield1 and 4OHT. Data are presented as mean ± SEM from 8 biological replicates. Statistical comparisons were performed using unpaired *t-*test. f) *Left*, representative image of RPE-1 cells expressing Cas9 6h post-transfection with the sgLAD. The locus at the LAD where the DNA damage is induced is visualised by FISH (green) and the nuclear periphery is stained with lamin B1 (red). DNA is stained by DAPI (blue). Scale bar 10µm. *Right*, percentage of LAD foci which colocalise with lamin B1 in cells transfected with either non-targeting control guide (sgCtrl) or sgLAD for 6 h and 24 h. Data are presented as mean ± SEM from 3 biological replicates. Statistical comparisons were performed using two-way ANOVA with Šídák correction for multiple comparisons. g) CUT&RUN followed by NGS showing enrichment of 53BP1 (top panels), and the log₂ ratio of lamin B1 signal relative to IgG (bottom panels) at a LAD upon a single DSB in RPE-1 cells expressing Cas9 and transfected with sgCtrl (grey) or sgLAD (red), 6 h post-transfection (*left*). iLAD region lacking a DSB as negative control (*right*). Blue shaded regions denote LAD annotations. Data represent the mean of 3 independent biological replicates.

To discriminate between these two scenarios and to examine how the nuclear envelope responds to DSBs at LADs, we compared the levels of nuclear periphery-associated proteins in cells with and without induced DNA damage. We found that DSB induction at LADs led to a reduction in lamin B1 and lamin B receptor (LBR) levels (Fig. 2b and Extended Data Fig. 2a, b). In contrast, other lamina-associated proteins—including lamin A, emerin, LAP2, and the nucleoporin NUP153—remained unchanged (Fig. 2b and Extended Data Fig. 2a, b). These results suggest that DSBs at LADs trigger reorganization of the nuclear lamina, potentially reflecting targeted remodelling of chromatin–nuclear envelope tethering.

To examine whether DSBs at LADs triggered their mobilization away from the nuclear envelope, we quantified LAD enrichment using the m⁶A-Tracer (Fig. 2c) in DpnI^-^ cells (no damage) and DpnI⁺ cells treated with +S +4OHT (damaged) (Fig. 2d, e). Upon damage, the peripheral LAD signal was significantly decreased (Fig. 2d, e), accompanied by a corresponding increase in nucleoplasmic m⁶A-Tracer intensity (Fig. 2d, e), suggesting that damaged LADs detach from the nuclear envelope and relocate to the nuclear interior. Rather than showing increased enrichment near the nucleoporin NUP153 signal, m⁶A-Tracer enrichment was in fact reduced, indicating that this behavior diverges from previously reported findings in *Drosophila*, where DNA damage triggers relocalization of heterochromatin regions to the nuclear pore complex (Extended Data Fig. 2c).

We next sought to develop an orthogonal system to study LAD dynamics in response to DSBs. To this end, we employed a Cas9 and single guide RNA (sgRNA) approach in RPE-1 cells (hTERT RPE-1 Cas9; Supplementary Table 1) to induce a single DSB within a previously characterized LAD^17^ (Extended Data Fig. 2d; sgLAD; gRNA #1; Supplementary Table 2). To compare the effect of DSB induction against the baseline of the same locus under non-damaging conditions, a non-targeting control guide RNA that does not induce DNA breaks (sgCtrl; crRNA #1; Supplementary Table 2) was also employed. As a further control, we targeted a genomic region that does not overlap with LADs (inter LAD: iLAD; Extended Data Fig. 2d; sgiLAD; gRNA #2; Supplementary Table 2). Expression of the targeting sgRNA successfully induced DSBs at the selected loci (Extended Data Fig. 2e), with cells containing two 53BP1 foci peaking 6 hours post-transfection (Extended Data Fig. 2f), confirming both the induction of DNA damage and the spatial positioning of these loci within the nucleus (Extended Data Fig. 2e). Parallel DNA insertion/deletion (indel) analysis revealed that while the peak of DNA damage occurred 6 hours after guide transfection, indels continued to accumulate over time, with the majority of alleles being damaged and repaired between 24–48 hours post-transfection (Extended Data Fig. 2g). The most prevalent indel was a +1 insertion, indicative of the usage of non-homologous end joining (NHEJ) at this locus (Extended Data Fig. 2g). Using fluorescent *in situ* hybridization (FISH), we observed that, under basal conditions, the LAD locus is closely associated with lamin B1 in 60% of cases (Fig. 2f). In marked contrast, only 25% of alleles of the iLAD region were associated with the nuclear lamina (Extended Data Fig. 2h). Strikingly, upon DNA damage, the association of the LAD with the nuclear envelope was reduced by half (Fig. 2f), reinforcing the notion that DSBs at LADs drive their dissociation from the nuclear envelope. Tracking the fate of the LAD 24 hours after DSB induction revealed its re-association with the nuclear envelope (Fig. 2f), consistent with the hypothesis that LAD detachment is a transient and reversible response to DNA damage. In contrast to the LAD locus, and consistent with the literature, upon DSB induction, the iLAD locus was associated with the lamina at an increased frequency (Extended Data Fig. 2h).

To complement the FISH analysis, we performed CUT&RUN using lamin B1 and 53BP1 antibodies to map lamina-associated and DNA damage–associated chromatin interactions respectively with next-generation sequencing (NGS) (Fig. 2g). Under basal conditions, CUT&RUN revealed strong enrichment of lamin B1 at the LAD locus (Fig. 2g, left), consistent with their close association with the nuclear lamina observed by FISH (Fig. 2f). In contrast, the iLAD region in chromosome 9 showed minimal lamin B1 occupancy (Fig. 2g, right), consistent with the FISH analysis (Extended Data Fig. 2h).

Upon induction of DNA damage in the LAD locus, lamin B1 occupancy was reduced centring on the Cas9 cut site (Fig. 2g, left), mirroring the decreased lamina association detected by FISH (Fig. 2f). Concurrently, 53BP1 occupancy increased specifically across this LAD domain spreading around the DSB (Fig. 2g, left), linking DNA damage to chromatin-lamina dissociation. The overlap of these two domains suggests that the reduction of lamin B1 occupancy is not confined to the immediate vicinity of the DSB but extends across the chromatin domain that has responded to the DDR. Importantly, LAD detachment cannot be explained by cell cycle redistribution, as damaged and undamaged cells displayed comparable cell cycle profiles at the peak of DNA damage (6 hours), with only a modest increase in the G2/M population observed at 24 hours (Extended Data Fig. 2i) where LADs have reattached at the nuclear envelope.

Together, these findings demonstrate that DNA double-strand breaks within LADs trigger a transient reorganization of chromatin–nuclear lamina interactions, resulting in the reversible release of damaged chromatin from the nuclear periphery into the nuclear interior.

### DNA damage-induced LAD detachment is not driven by loss of LBR or lamin B1

Since both lamin B1 and LBR become destabilized from the nuclear periphery following DNA damage, we investigated whether these nuclear lamina components contribute to DNA damage-induced LAD detachment. To assess the role of LBR, we first depleted LBR in DpnI cells using siRNA (Extended Data Fig. 3a). Consistent with its established function in tethering heterochromatin to the nuclear periphery, LBR depletion reduced LAD association with the nuclear envelope even in the absence of DNA damage (Extended Data Fig. 3b). Importantly, induction of DNA damage in LBR-depleted cells caused a further decrease in LAD association with the nuclear periphery (Extended Data Fig. 3b). These findings indicate that loss of LBR alone is not sufficient to account for DNA damage-induced LAD detachment, and suggest that additional mechanisms contribute to this process. In agreement with this conclusion, LBR depletion in RPE-1 cells using a single guide RNA targeting the *LBR* coding sequence, which generated deletions within the gene (Extended Data Fig. 3c, d), did not impair LAD detachment following induction of a single DSB (Extended Data Fig. 3e).

We next examined the contribution of lamin B1 by generating lamin B1-deficient RPE-1 cells through Cas9-mediated targeting of the *LMNB1* coding sequence (Extended Data Fig. 3f, g). DNA damage induced within LADs resulted in a similar level of LAD detachment in lamin B1-deficient and control cells (Extended Data Fig. 3h), indicating that although lamin B1 levels decrease at the nuclear periphery following DNA damage, this reduction is not responsible for LAD detachment. These experiments were performed exclusively in RPE-1 cells because DNA damage induction in DpnI cells depends on expression of the Dam–lamin B1 fusion protein. Consequently, depletion of lamin B1 in this system would disrupt the DNA damage induction strategy itself, preventing meaningful interpretation of the results.

Collectively, these results show that changes in lamin B1 and LBR accompany DNA damage but are not sufficient to drive LAD detachment.

### ATM kinase and its downstream effectors are required for LAD detachment from the nuclear envelope

We have previously demonstrated that a DSB at a locus tethered to the nuclear lamina is predominantly repaired via NHEJ and microhomology-mediated end-joining (MMEJ), while recruitment of homologous recombination (HR) factors is suppressed^18^. Consistent with this, Cas9-induced DSBs exhibit increased usage of MMEJ over NHEJ when targeted to genomic regions associated with LAD heterochromatin (defined as chromatin enriched in H3K9me2, H3K27me3, late replicating DNA, and interacts with lamin B1) and lamin B1^17^. To investigate whether DNA repair pathway choice influences LAD detachment, we inhibited key components of NHEJ (DNAPKcs), MMEJ (Polθ), and HR (RAD51). None of these perturbations affected LAD detachment (Fig. 3a and Extended Data Fig. 4a). Interestingly, prior inhibition of ATM prevented LAD detachment from the nuclear periphery in DpnI cells (Fig. 3a and Extended Data Fig. 4b) independently from changes in the cell cycle (Extended Data Fig. 1e, left), indicating a specific role for ATM signaling in facilitating LAD mobilization in response to DNA damage.

**Figure 3:**
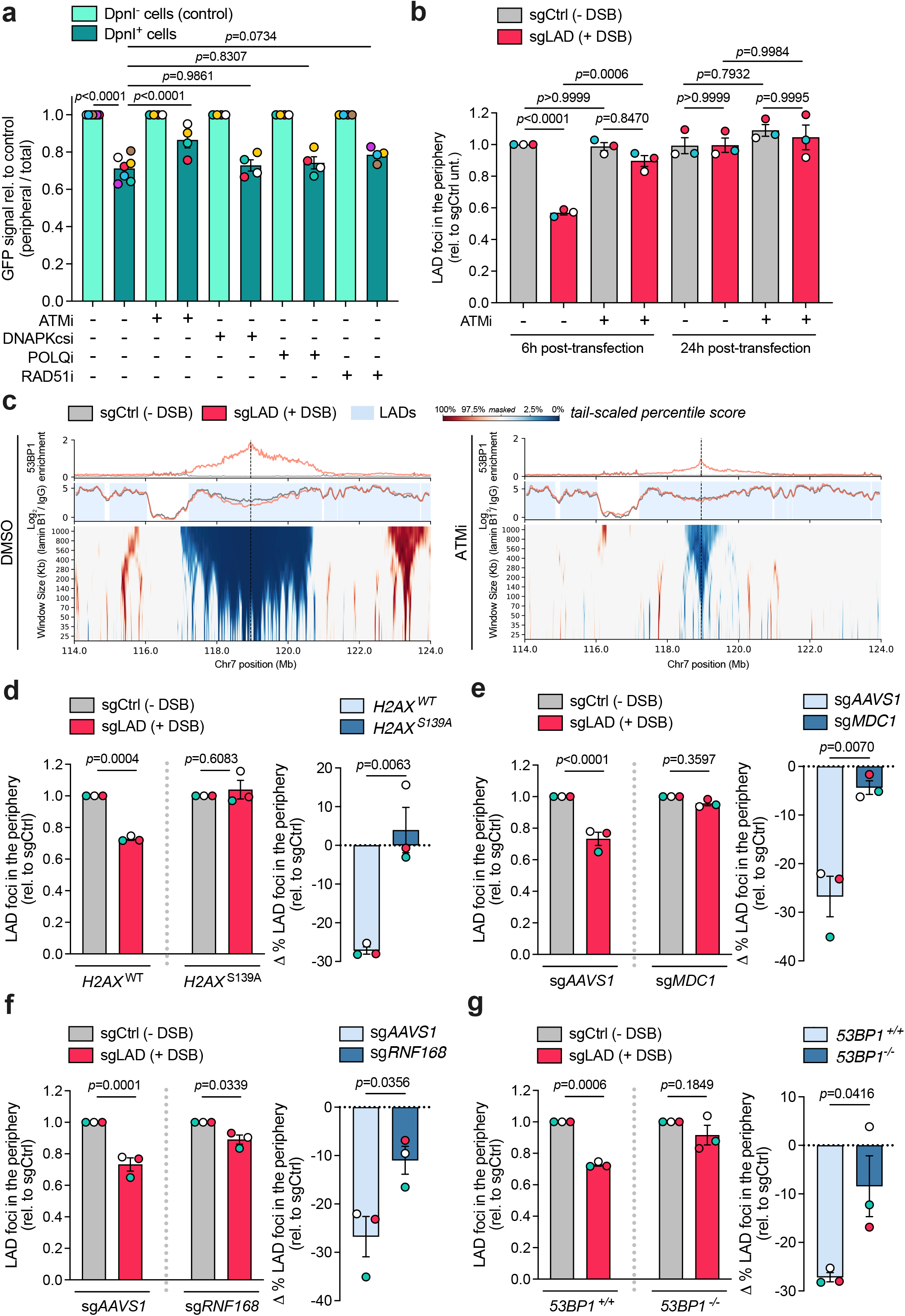
LAD detachment induced by DNA damage involves the ATM signaling axis. a) Quantification of the relative abundance of m^6^A-Tracer at the nuclear rim in cells expressing or not DpnI, treated with Shield1 and 4OHT in control conditions (DMSO) or treated with inhibitors for ATM (ATMi, 10µM), DNAPKcs (DNAPKi, 10µM), Pol theta (PolQi, 10µM) and RAD51i (20µM). Data are presented as mean ± SEM from at least 4 biological replicates. Statistical comparisons were performed using two-way ANOVA with Šídák correction for multiple comparisons. b) Quantification of LAD foci which colocalise with lamin B1 in cells transfected with either non-targeting control guide (sgCtrl) or sgLAD for 6 h and 24 h treated with ATMi (10µM) or DMSO (-). Data are presented as mean ± SEM from 3 biological replicates. Statistical comparisons were performed using three-way ANOVA with DSB, ATMi, and time as factors, followed by Šídák’s multiple comparisons test. c) CUT&RUN sequencing maps of a 10 Mb region on chromosome 7 in RPE-1 cells expressing Cas9 in DMSO treated conditions (*left*) and ATMi (10µM) treated conditions (*right*), centered on the site of DSB induction denoted by the dashed vertical line. Samples transfected for 6 h with sgCtrl are displayed in grey, and sgLAD are displayed in red. *Top,* enrichment of 53BP1 signal. *Middle,* the log₂ ratio of lamin B1 signal relative to IgG. Blue shaded regions denote LAD annotations. *Bottom,* domainograms represent the ranked log₂(lamin B1 / IgG) ratio deltas (sgLAD - sgCtrl) across a range of increasing window sizes (y-axis). Each window is ranked independently across all genome-wide windows of log₂(lamin B1 / IgG) ratio deltas. The upper and lower 5% of relative lamin B1 signal changes genome-wide following DSB induction are represented in red and blue respectively. The central 90% of log₂(lamin B1 / IgG) ratio delta are masked. Three biological replicates are presented for DMSO conditions, and two biological replicates for ATMi (10µM) treated conditions. d) *Left*, quantification of LAD foci which colocalise with lamin B1 in WT (*H2AX*^WT^) and *H2AX*^S139A^ mutant RPE-1 cells transfected with Cas9 and either non-targeting control guide (sgCtrl) or sgLAD for 7 h. Data are presented as mean ± SEM from 3 biological replicates. Statistical comparisons were performed using two-way ANOVA followed by Šídák’s multiple comparisons test. *Right*, percentage change in LAD foci in the nuclear periphery (sgLAD - sgCtrl) relative to the corresponding sgCtrl condition. Data are presented as mean ± SEM from 3 biological replicates. Statistical comparisons were performed using unpaired *t*-test. e) *Left*, quantification of LAD foci which colocalise with lamin B1 in mock-depleted (sg*AAVS1*) or MDC1-depleted (sg*MDC1*) RPE-1 cells transfected with either non-targeting control guide (sgCtrl) or sgLAD for 6 h. Data are presented as mean ± SEM from 3 biological replicates. Statistical comparisons were performed using two-way ANOVA followed by Šídák’s multiple comparisons test. *Right*, percentage change in LAD foci in the nuclear periphery (sgLAD - sgCtrl) relative to the corresponding sgCtrl condition. Data are presented as mean ± SEM from 3 biological replicates. Statistical comparisons were performed using unpaired *t*-test. f) *Left*, quantification of LAD foci which colocalise with lamin B1 in mock-depleted (sg*AAVS1*) or RNF168-depleted (sg*RNF168*) RPE-1 cells transfected with either non-targeting control guide (sgCtrl) or sgLAD for 6 h. Data are presented as mean ± SEM from 3 biological replicates. Statistical comparisons were performed using two-way ANOVA followed by Šídák’s multiple comparisons test. *Right*, percentage change in LAD foci in the nuclear periphery (sgLAD - sgCtrl) relative to the corresponding sgCtrl condition. Data are presented as mean ± SEM from 3 biological replicates. Statistical comparisons were performed using unpaired *t*-test. g) *Left*, quantification of LAD foci which colocalise with lamin B1 in WT (*53BP1*^+/+^) and 53BP1-deficient (*53BP1*^-/-^) cells transfected with Cas9 and either non-targeting control guide (sgCtrl) or sgLAD for 7 h. Data are presented as mean ± SEM from 3 biological replicates. Statistical comparisons were performed using two-way ANOVA followed by Šídák’s multiple comparisons test. *Right*, percentage change in LAD foci in the nuclear periphery (sgLAD - sgCtrl) relative to the corresponding sgCtrl condition. Data are presented as mean ± SEM from 3 biological replicates. Statistical comparisons were performed using unpaired *t*-test. Note: *H2AX*^WT^ and *53BP1^+/+^*are the same WT cell line. All six conditions (WT, *H2AX*^S139A^, and *53BP1^-/-^*, ±DSB) were processed in the same experiment. The WT control lanes shown in panels (d) and (g) derive from the same samples and are presented separately for clarity of comparison with each gene knock-out. Note: All six conditions (sg*AAVS1*, sg*MDC1*, and sg*RNF168*, ±DSB) were processed in the same experiment. The sg*AAVS1* control lanes shown in panels (e) and (f) derive from the same samples and are presented separately for clarity of comparison with each gene depletion.

Likewise, inhibition of ATM effectively blocked Cas9-induced DSB-driven LAD detachment (Fig. 3b), without affecting the cell cycle (Extended Data Fig. 4c), and suppressed the relocation of the damaged iLAD to the nuclear periphery (Extended Data Fig. 4d). Genome-wide tail-scaled percentile scoring of the mean delta signal between damaged and undamaged conditions (sgLAD – sgCtrl) at the Cas9 induced break revealed a several mega-base domain of LAD detachment from the nuclear envelope in an ATM-dependent manner (Fig. 3c). Intriguingly, LAD detachment was followed by increased interactions with neighboring chromatin regions with the nuclear lamina in the DMSO condition only (Fig. 3c). Together, these findings demonstrate that DNA damage at LADs triggers a reorganization of the nuclear envelope and mobilization of associated chromatin, establishing the ATM signaling pathway as a central orchestrator of this dynamic response.

As ATM is responsible for the phosphorylation and recruitment of key DDR mediators and effectors to sites of DSBs, we next investigated whether its role in LAD detachment is mediated through the canonical ATM signaling cascade. To this end, we assessed the contribution of H2AX, MDC1, RNF168, and 53BP1 to LAD detachment. We first investigated whether H2AX phosphorylation contributes to LAD detachment following DNA damage. To this end, we used either H2AX-deficient cells (Extended Data Fig. 4e and Supplementary Table 1) or *H2AX*^S139A^ knock-in cells, in which the serine residue at position 139 is replaced by alanine, thereby preventing H2AX phosphorylation (Extended Data Fig. 4f and Supplementary Table 1). As expected, induction of DSBs at LADs using sgLAD led to a reduction in the interaction between the LAD loci and lamin B1, as assessed by immunoFISH (Fig. 3d and Extended Data Fig. 4g). Notably, in cells lacking H2AX or expressing the non-phosphorylatable *H2AX*^S139A^ mutant, LADs remained associated with the nuclear periphery, phenocopying the effect observed following ATM inhibition (Fig. 3d and Extended Data Fig. 4g).

Using the same approach, we next investigated the roles of MDC1, RNF168, and 53BP1 in DNA damage-induced LAD detachment. To this end, we generated pooled *MDC1* and *RNF168* knockout cell populations using guide RNAs targeting their coding sequences (Extended Data Fig. 5a-d) and utilised a 53BP1-deficient RPE-1 cell line (Extended Data Fig. 4f and Supplementary Table 1). Similar to the phenotypes observed in H2AX-deficient cells, loss of MDC1, RNF168, or 53BP1 impaired LAD detachment following DSB induction, resulting in the persistent association of damaged LADs with the nuclear lamina (Fig. 3e-g). Notably, although RNF168 depletion had a milder effect, it phenocopied the effects of pharmacological inhibition of ubiquitination (Extended Data Fig. 5e), indicating that ubiquitin-dependent signaling downstream of ATM is required for the relocalization of damaged LADs away from the nuclear periphery.

53BP1 is a key mediator of the DNA damage response that promotes NHEJ and suppresses DNA end resection through the recruitment of RIF1 and the shieldin/CST complex^19^. In addition, 53BP1 facilitates the propagation of ATM signaling, particularly within heterochromatin domains, likely by organizing the three-dimensional architecture of DNA repair within chromatin microdomains^20^. Consistent with this role, pATM induction was markedly reduced in 53BP1-deficient cells compared with wild-type cells, indicating that 53BP1 reinforces ATM activation through a positive feedback mechanism (Extended Data Fig. 5f).

Together, these findings demonstrate that the ATM–H2AX–MDC1–RNF168–53BP1 axis promotes LAD detachment in response to DNA damage. They further identify phosphorylation- and ubiquitin-dependent chromatin signaling as the key mechanistic driver of this process by sustaining ATM activity and amplifying damage signaling within chromatin.

### LAD detachment extends beyond the local chromatin accessibility induced by DNA damage

As ATM is known to promote chromatin decondensation surrounding DSBs, particularly within heterochromatin, we investigated whether its role in LAD detachment is mediated through chromatin opening. To determine whether DSB induction within LADs increases local chromatin accessibility in an ATM-dependent manner, we performed ATAC-seq in the presence or absence of Cas9-generated DNA damage and following ATM inhibition. As expected, the targeted LAD region exhibited low chromatin accessibility under basal conditions (sgCtrl, DMSO) (Fig. 4a), consistent with the compact chromatin organization characteristic of LADs. Induction of a DSB led to a marked increase in chromatin accessibility surrounding the break site (sgLAD, DMSO), with the strongest effect observed within approximately 15 kb of the lesion and the highest signal detected within the first 5 kb (Fig. 4a). Therefore, the region displaying increased chromatin accessibility is substantially smaller than the domain marked by 53BP1 accumulation, which serves as a proxy for DNA damage response spreading, and also smaller than the chromatin domain that appears to detach from the nuclear lamina following DNA damage (Fig. 3c). Inhibition of ATM attenuated this increase in accessibility both in terms of maximum ATAC-seq signal around the sgLAD cut site, and signal width extending from the DSB, demonstrating that chromatin opening close to DSBs within LADs is ATM dependent (Fig. 4a).

**Figure 4:**
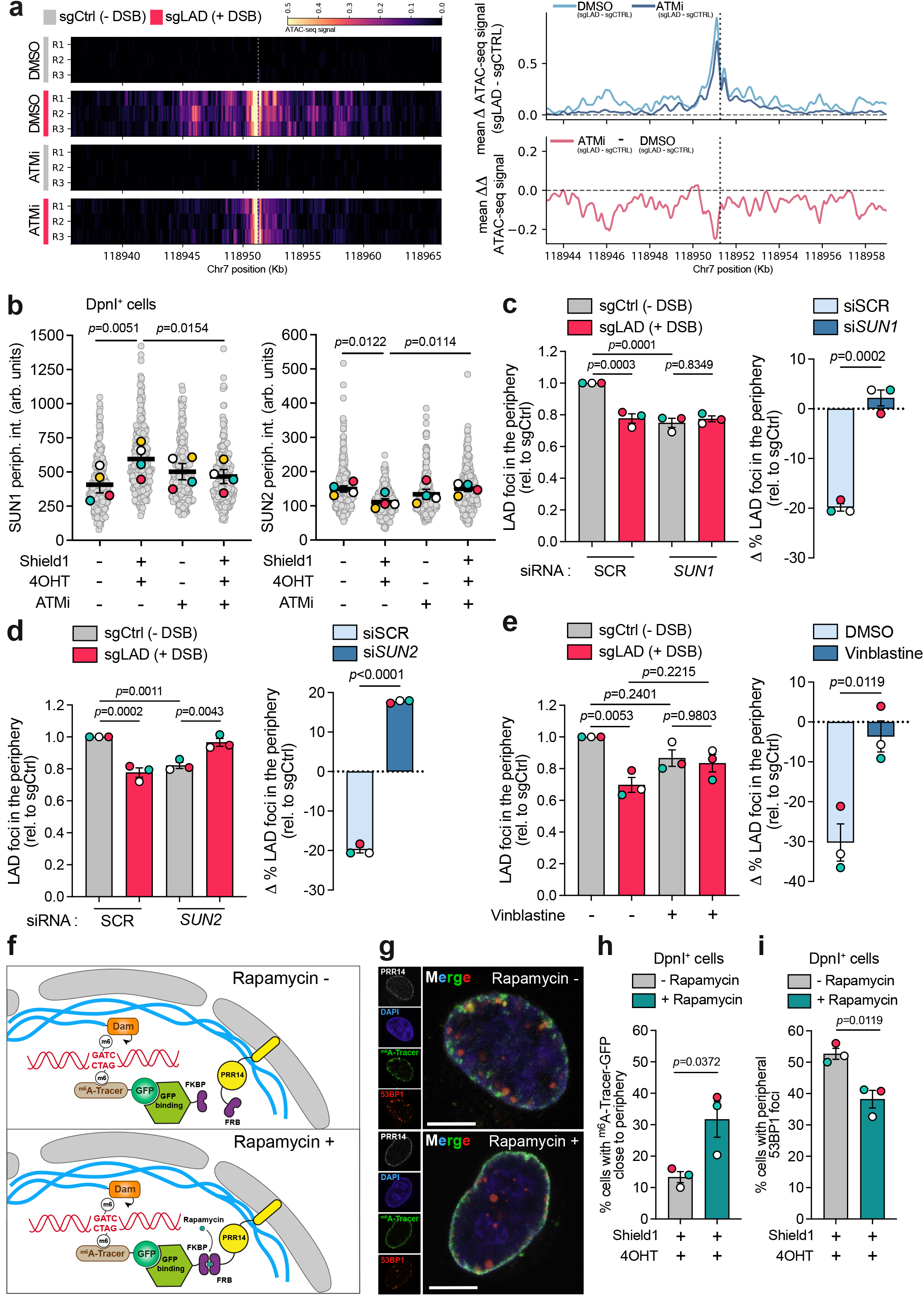
LAD detachment is crucial for prompt DNA damage response and DNA repair. a) *Left,* heatmaps of ATAC-seq signal across a 30 Kb region on chromosome 7 in RPE-1 cells expressing Cas9 transfected with either non-targeting control guide (sgCtrl) or sgLAD for 6 h in DMSO treated conditions and ATMi (10µM) treated conditions. Site of DSB induction is denoted by the dotted vertical line. Three independent biological replicates for each condition are represented. *Top right,* the mean difference in ATAC-seq signal between sgLAD and sgCtrl transfected sample tracks for DMSO and ATMi treated samples (mean Δ ATAC-seq signal) across a 15 Kb region on chromosome 7 with 200bp smoothing. *Bottom right,* the difference between the mean Δ ATAC-seq signal of ATMi and DMSO treated conditions is represented below in pink, termed as the mean ΔΔ ATAC-seq signal. Vertical dotted lines denote the site of DSB induction. b) Quantification in arbitrary units (linear scale) of SUN1 and SUN2 at the nuclear periphery in cells expressing DpnI in the absence (-Shield1 -4OHT) and presence (+Shield1 +4OHT) of DSBs at LADs in control conditions (DMSO) or treated with ATMi (10µM). Data are presented as mean ± SEM from 3 biological replicates. Statistical comparisons were performed using repeated measures two-way ANOVA followed by Šídák’s multiple comparisons test. c) *Left*, quantification of LAD foci which colocalise with lamin B1 in cells depleted using siRNA for *SUN1* or control (SCR) and then transfected with either non-targeting sgCtrl or sgLAD for 6 h. Data are presented as mean ± SEM from 3 biological replicates. Statistical comparisons were performed using two-way ANOVA followed by Šídák’s multiple comparisons test. *Right*, percentage change in LAD foci in the nuclear periphery (sgLAD - sgCtrl) relative to the corresponding sgCtrl condition. Data are presented as mean ± SEM from 3 biological replicates. Statistical comparisons were performed using unpaired *t*-test. d) *Left*, quantification of LAD foci which colocalise with lamin B1 in cells depleted using siRNA for *SUN2* or control (SCR) and then transfected with either non-targeting sgCtrl or sgLAD for 6 h. Data are presented as mean ± SEM from 3 biological replicates. Statistical comparisons were performed using two-way ANOVA followed by Šídák’s multiple comparisons test. *Right*, percentage change in LAD foci in the nuclear periphery (sgLAD - sgCtrl) relative to the corresponding sgCtrl condition. Data are presented as mean ± SEM from 3 biological replicates. Statistical comparisons were performed using unpaired *t*-test. e) *Left*, quantification of LAD foci which colocalise with lamin B1 in cells transfected with either non-targeting control guide (sgCtrl) or sgLAD for 6 h treated with the microtubule polymerization inhibitor Vinblastine (100nM) or DMSO (-). Data are presented as mean ± SEM from 3 biological replicates. Statistical comparisons were performed using two-way ANOVA followed by Šídák’s multiple comparisons test. *Right*, percentage change in LAD foci in the nuclear periphery (sgLAD - sgCtrl) relative to the corresponding sgCtrl condition. Data are presented as mean ± SEM from 3 biological replicates. Statistical comparisons were performed using unpaired *t*-test. f) Schematic representation of the experimental system for controlled tethering of LADs to the nuclear periphery. GFP nanobody fused to FKBP binds the GFP tag on m^6^A-Tracer, while the nuclear envelope protein PRR14 fused to FRB localizes to the nuclear envelope. Upon Rapamycin treatment, FKBP and FRB dimerize, bridging LADs to the nuclear periphery. g) Representative images of DpnI^+^ cells treated with Shield1 and 4OHT, expressing FRB fused to PRR14 and GFP-nanobody fused to FKBP and treated or not with Rapamycin. In the absence of Rapamycin, broad m^6^A-Tracer-GFP (green) recruitment to methylated LADs is observed. 53BP1 is labelled in red, PRR14 in white and DAPI in blue. Upon treatment with Rapamycin, LADs are tethered to the nuclear envelope through FKB-FKBP binding, leading to the defined, continuous GFP ring visible at the nuclear edge. h) Quantification of DpnI^+^ cells treated with Shield1 and 4OHT with m^6^A-Tracer-GFP localised close to the nuclear periphery in the absence or presence of Rapamycin, as indicated. Data are presented as mean ± SEM from 3 biological replicates. Statistical comparisons were performed using unpaired *t*-test. i) Percentage of cells showing 53BP1 foci at the nuclear periphery in DpnI^+^ cells expressing FRB fused to PRR14 and GFP-nanobody fused to FKBP, treated with Shield1 and 4OHT and in the absence or presence of Rapamycin, as indicated. Data are presented as the percentage of total PRR14 positive nuclei and is the mean ± SEM from 3 biological replicates. Statistical comparisons were performed using unpaired *t*-test. Note: All six conditions (siSCR, si*SUN1*, and si*SUN2*, ±DSB) were processed in the same experiment. The siSCR control lanes shown in panels (c) and (d) derive from the same samples and are presented separately for clarity of comparison with each depletion.

To further investigate the relationship between chromatin accessibility and LAD detachment, we performed ATAC-seq in *H2AX*^S139A^ knock-in and 53BP1-deficient RPE-1 cells, in which LAD detachment is impaired (Extended Data Fig. 6a-d). We found that although chromatin accessibility was reduced in the *H2AX*^S139A^ knock-in cells and phenocopied ATM inhibition (Extended Data Fig. 6a, c), 53BP1-deficient cells did not show any difference compared to wild-type cells (Extended Data Fig. 6b, d), breaking the correlation between LAD detachment and chromatin accessibility. These findings suggest that changes in local chromatin accessibility do not necessarily accompany or cause LAD detachment and may therefore represent a distinct response to DNA damage.

### The LINC complex modulates microtubule-dependent LAD detachment in response to DSBs

Chromatin mobility at DSBs and telomeres is known to be driven by cytoskeletal forces transmitted through the LINC complex, with SUN1 playing a role in tethering DSBs within active genes near the nuclear periphery, and both SUN1 and SUN2 in the mobility of telomeres and in the formation of dsbNETs^8,9,21^. Therefore, we sought to explore how SUN1 and SUN2 respond to DSBs within LADs and whether LINC components contribute to LAD dynamics following DNA damage. To this end, we measured the changes in the intensity of SUN1 and SUN2 in the periphery in response to DNA damage. DSB induction at LADs triggered a dynamic response from the LINC complex, with SUN1 robustly accumulating at the nuclear envelope and SUN2 intensity moderately reduced (Fig. 4b and Extended Data Fig. 6e-g), indicating opposite behaviors. This response was blocked by treatment with the ATM inhibitor (Fig. 4b), suggesting that ATM drives LINC remodeling at the nuclear envelope in response to LAD-associated DSBs—or that the LINC complex senses LAD detachment and actively responds to it.

Depletion of both SUN1 and SUN2 reduced LAD tethering to the nuclear envelope under basal conditions. Notably and in line with the response above, depleting SUN1 did not affect LAD relocalization following DSB induction (Fig. 4c and Extended Data Fig. 6h, i). By contrast, SUN2 depletion actively suppressed LAD relocalization (Fig. 4c, d and Extended Data Fig. 6j, k, l). Together, these findings indicate distinct roles for SUN1 and SUN2 in LAD organization upon DSBs. Whereas both primarily contribute to the constitutive tethering of LADs to the nuclear envelope under steady-state conditions, SUN2 is specifically required for the DNA damage-induced detachment and relocalization of LADs. These results further suggest that SUN1 may instead function in re-establishing LAD tethering to the nuclear envelope following completion of DNA repair.

Through interactions with the nuclear lamina and chromatin, the LINC complex transmits forces between the nucleus and the cytoplasm. On the cytoplasmic side, it links to actin filaments via nesprins, allowing myosin-generated contractile forces to be conveyed to the nucleus, and it also associates with microtubules through motor proteins such as dynein and kinesin^22^. To determine the respective contributions of these cytoskeletal components to LINC-mediated force transmission required for LINC-dependent LAD detachment, we inhibited actin polymerization, myosin contractility, and microtubule dynamics upon DNA damage. We first observed that myosin perturbation by blebbistatin had no detectable impact on LAD detachment (Extended Data Fig. 7a). Inhibition of actin polymerization by latrunculin B and Arp2/3 inhibition produced a modest effect in the DpnI system and no effect in the Cas9 system (Extended Data Fig. 7b-d). By contrast, inhibition of microtubule dynamics with vinblastine substantially impaired LAD detachment in both the DpnI and Cas9 systems (Fig. 4e, Extended Data Fig. 7e). These findings point to microtubules as the primary cytoskeletal driver of LAD detachment, with little to no wide contribution from actin polymerization.

### Tethering of LADs at the nuclear envelope impairs the DNA damage response and delays DNA repair

To investigate the functional significance of nuclear periphery rewiring during DNA damage response and repair, we examined how inhibition of LAD detachment affects these processes. Specifically, we measured the kinetics of DNA repair by assessing the rate at which indels were generated over a 48 hours period, under conditions that altered LAD relocation. As shown previously, repair events accumulate progressively over time, with the majority of indels forming between 24- and 48-hours post-damage (Extended Data Fig. 2g). When cells are treated with the ATM inhibitor, this process is significantly delayed (Extended Data Fig. 8a), consistent with a specific role for ATM in the repair of Cas9-induced DSBs within heterochromatin^23^. In contrast, ATM inhibition following induction of a DSB at an iLAD locus resulted in only a transient delay at the onset of repair and did not impair the prompt emergence of the full spectrum of indels, suggesting an effect specific to DSBs induced at the LAD locus (Extended Data Fig. 8b). *H2AX*^S139A^ mutant and 53BP1-deficient cells exerted a similar delay in repair kinetics, with 71% and 77% of sequences exhibiting +1 insertions respectively at 24h compared to 87% in WT cells. This persists at 48h, with 86% of sequences containing the +1 insertion in both mutants compared to 97% and 96% in WT cells (Extended Data Fig. 8c, d).

We next asked whether LAD detachment alters DNA repair pathway choice. Inhibition of Polθ caused a markedly greater increase in chromosomal instability at a LAD-associated DSB than inhibition of RAD51 (Extended Data Fig. 8e), indicating that MMEJ remains the predominant repair pathway despite detachment from the nuclear periphery. In contrast, a DSB induced in the iLAD locus showed a greater dependence on HR (Extended Data Fig. 8e). Thus, although DNA damage triggers LAD relocalization, repair pathway choice remains unchanged. This is consistent with our previous finding that DSBs generated while tethered at the nuclear periphery are repaired predominantly by MMEJ, with only a limited contribution from HR^18^. These results suggest that LAD detachment is important for prompt DNA repair but does not alter DNA repair choice.

To directly prove the involvement of LAD detachment in DNA repair and to exclude the possibility that the effects seen above are due to generic roles of ATM and H2AX in DDR and repair, we developed a system to artificially maintain LAD–nuclear envelope association using a rapamycin-inducible dimerization system^24^. This system relies on two components: FKBP12 (FKBP), which binds rapamycin, and the FKBP–rapamycin binding domain (FRB) of mTOR (Fig. 4f). In our design, the chromatin-tethering domain of PRR14, a protein that tethers heterochromatin to the nuclear lamina^25^, was fused to FRB, while 3XFKBP was fused to a GFP nanobody (Fig. 4f). This system was expressed in HT1080 cells stably expressing the GFP-tagged m^6^A-Tracer. Upon rapamycin addition, FKBP and FRB heterodimerize, resulting in the tethering of GFP-labeled chromatin to the nuclear periphery via PRR14 (Fig. 4g, h and Extended Data Fig. 8f). Interestingly, forced tethering of LADs to the nuclear envelope diminished DDR, as indicated by reduced presence of peripheral 53BP1 foci (Fig. 4i), and had no effect in cells where DNA damage was not induced (DpnI^-^ cells, Extended Data Fig. 8g), suggesting that LADs are released from the nuclear envelope to be readily recognised by the DDR machinery.

### LAD detachment preserves genome stability and maintains nuclear envelope integrity

To determine whether inhibiting the release of the DSB from the lamina results in increased genomic instability, we first examined chromosomal breaks on chromosome 7, where the LAD-associated DSB was introduced via Cas9, using metaphase spread analysis with whole chromosome painting. This revealed a higher incidence of chromosomal breaks within chromosome 7 following DSB induction at LADs (Fig. 5a, c). Likewise, DSB induction using DpnI caused increased breaks on both chromosomes 7 and 9 (Fig. 5b, d). In both cellular models, the frequency of chromosomal breaks rose further upon inhibition of ATM (Fig. 5c, d). In line with their role in LAD detachment and DNA repair delay, induction of a DSB in the LAD locus in *H2AX*^S139A^ mutant or H2AX-, RNF168- and 53BP1-deficient RPE-1 cells increased the frequency of chromosomal breaks (Extended Data Fig. 9a-d), indicating that the ATM-H2AX-MDC1-RNF168-53BP1 axis is critical for maintaining genome integrity at peripheral DSBs. These results align with the role of ATM in promoting prompt DNA repair at these DSBs (Extended Data Fig. 8a). Notably, co-depletion of SUN1 and SUN2 resulted in a significant increase in chromosomal breaks (Fig. 5e and Extended Data Fig. 6l), revealing that, despite their distinct roles in regulating LAD dynamics following DSB induction, both proteins are important for maintaining LAD integrity. Consistent with a genetic interaction between SUN1/SUN2 and ATM, inhibiting ATM in cells lacking both SUN1 and SUN2 did not further increase the frequency of chromosomal breaks (Fig. 5f), suggesting that they function within the same pathway to suppress genome instability.

**Figure 5:**
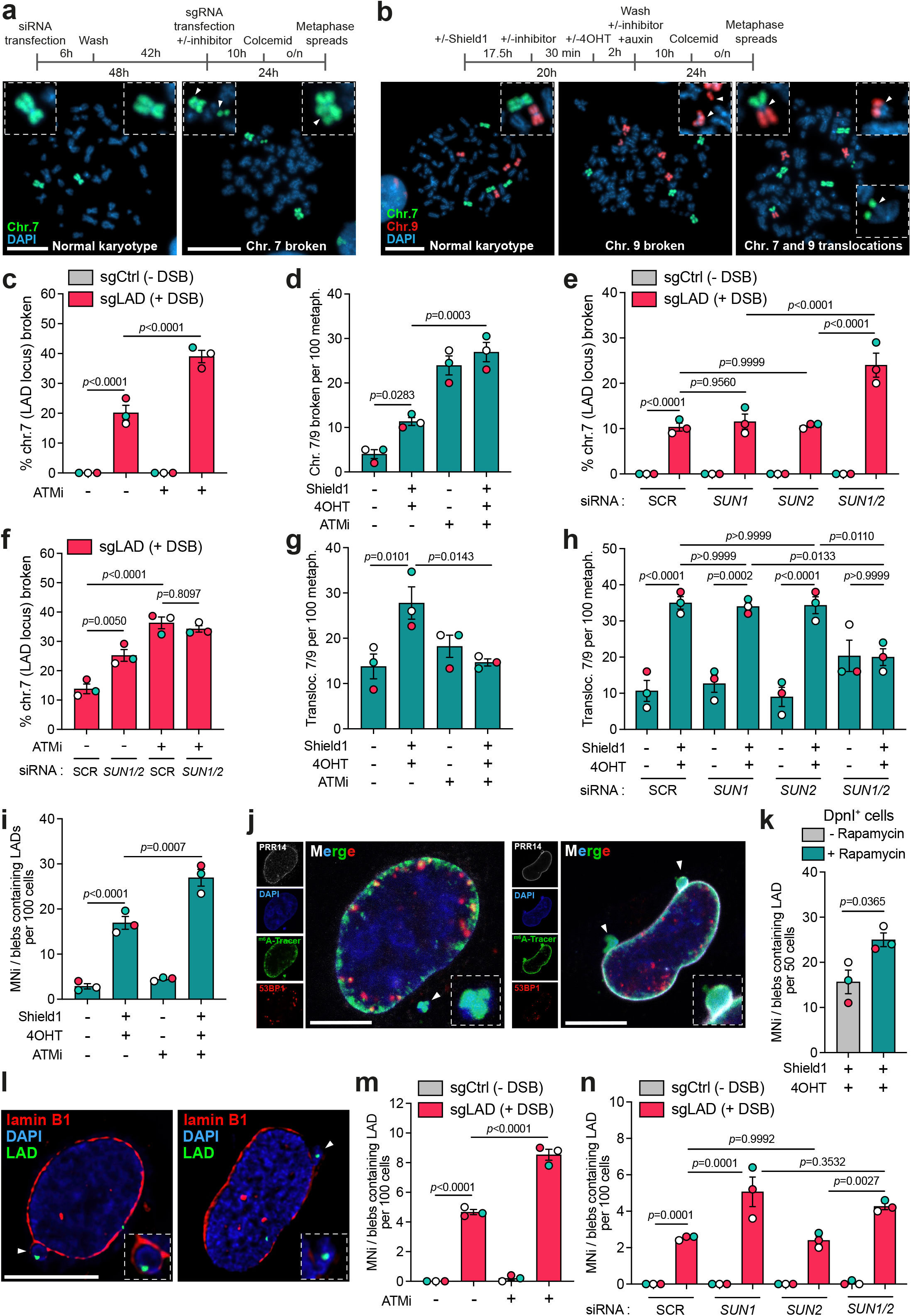
Failure to detach leads to LAD genomic instability and compromises nuclear envelope integrity. a) Experimental outline and representative microscopy images of metaphase spreads in RPE-1 cells in which chromosome 7 (chr.7) is painted in green in the absence (*left*) and presence of DNA damage (*right*) induced by sgLAD. Scale bar 20µm. b) Experimental outline and representative microscopy images of metaphase spreads from DpnI expressing cells, with chromosome 7 (chr.7) labeled in green and chromosome 9 (chr.9) in red. The images depict three conditions: untreated cells (*left*), DNA damage-induced cells showing visible chromosomal breaks (*middle*), and DNA damage-induced cells exhibiting translocations (*right*). Scale bar: 20 µm. c) Percentage of broken chr.7 in RPE-1 cells expressing Cas9 and transfected with a non-targeting control guide (sgCtrl) or sgLAD treated with ATM inhibitor (ATMi; 10µM) or with DMSO (-). Data are presented as mean ± SEM from 3 biological replicates. Statistical comparisons were performed using two-way ANOVA followed by Šídák’s multiple comparisons test. d) Quantification of broken chr.7 and chr.9 per 100 metaphases in DpnI expressing cells in the absence (-Shield1 -4OHT) and presence (+Shield1 +4OHT) of DNA damage treated treated with ATMi (10µM) or DMSO (-). Data are presented as mean ± SEM from 3 biological replicates. Statistical comparisons were performed using two-way ANOVA followed by Šídák’s multiple comparisons test. e) Percentage of broken chr.7 in RPE-1 cells expressing Cas9 in control (siSCR) or depleted for *SUN1* (si*SUN1*), *SUN2* (si*SUN2*) or both (si*SUN1*/*SUN2*) and then transfected with sgCtrl or sgLAD. Data are presented as mean ± SEM from 3 biological replicates. Statistical comparisons were performed using two-way ANOVA followed by Šídák’s multiple comparisons test. f) Percentage of broken chr.7 per 100 metaphases in RPE-1 cells expressing Cas9 in control (siSCR) or depleted for *SUN1* (si*SUN1*), *SUN2* (si*SUN2*) or both (si*SUN1*/*SUN2*) and then transfected with sgCtrl or sgLAD treated with ATM inhibitor (ATMi; 10µM) or with DMSO (-). Data are presented as mean ± SEM from 3 biological replicates. Statistical comparisons were performed using two-way ANOVA followed by Šídák’s multiple comparisons test. g) Quantification of chromosomal translocations involving chr.7 and chr.9 per 100 metaphases in DpnI expressing cells in the absence (-Shield1 -4OHT) and presence (+Shield1 +4OHT) of DNA damage treated with ATMi (10µM) or DMSO (-). Data are presented as mean ± SEM from 3 biological replicates. Statistical comparisons were performed using two-way ANOVA followed by Šídák’s multiple comparisons test. h) Quantification of chromosomal translocations involving chr.7 and chr.9 per 100 metaphases in DpnI expressing cells in the absence (-Shield1 -4OHT) and presence (+Shield1 +4OHT) of DNA damage in control (siSCR) or depleted for *SUN1* (si*SUN1*), *SUN2* (si*SUN2*) or both (si*SUN1*/*SUN2*). Data are presented as mean ± SEM from 3 biological replicates. Statistical comparisons were performed using two-way ANOVA followed by Šídák’s multiple comparisons test. i) Quantification of MNi/blebs containing LADs (m^6^A-Tracer) in DpnI expressing cells in the absence (-Shield1 -4OHT) or presence (+Shield1 +4OHT) of DNA damage, treated with either DMSO or ATM inhibitor (ATMi, 10µM). Data are presented as mean ± SEM from 3 biological replicates. Statistical comparisons were performed using two-way ANOVA followed by Šídák’s multiple comparisons test. j) Representative images of DpnI expressing cells bearing MNi/blebs transfected with PRR14-FRB/FKBP-GFP nanobodies following DSB induction with Shield1 and 4OHT and treated with Rapamycin. m^6^A-Tracer is labelled in green, 53BP1 is labelled in red, PRR14 in white and DAPI in blue. White arrows indicate MNi/blebs containing LADs (m^6^A-Tracer-GFP). Scale bar 10µm. k) Quantification of cells with MNi/blebs formation in DpnI expressing cells transfected with PRR14-FRB/FKBP-GFP nanobodies following DSB induction with Shield1 and 4OHT and in the absence or presence of Rapamycin, as indicated. Data are presented as mean ± SEM from 3 biological replicates. Statistical comparisons were performed using unpaired *t*-test. l) Representative images of RPE-1 cells expressing Cas9 and transfected with sgLAD bearing MNi/blebs containing the LAD locus. The locus at the LAD where the DNA damage is induced is visualised by FISH (green) and the nuclear periphery is stained with lamin B1 (red). DNA is stained by DAPI (blue). Scale bar 10µm. m) Quantification of MNi/blebs containing the LAD locus in RPE-1 cells expressing Cas9 and transfected with non-targeting control (sgCtrl) and sgLAD in control conditions (DMSO) or treated with ATMi (10µM). Data are presented as mean ± SEM from 3 biological replicates. Statistical comparisons were performed using two-way ANOVA followed by Šídák’s multiple comparisons test. n) Quantification of MNi/blebs containing the LAD locus in RPE-1 cells expressing Cas9 in control conditions (SCR) or depleted for *SUN1* (si*SUN1*), *SUN2* (si*SUN2*) or both (si*SUN1*/*SUN2*) and then transfected with non-targeting control (sgCtrl) or sgLAD. Data are presented as mean ± SEM from 3 biological replicates. Statistical comparisons were performed using two-way ANOVA followed by Šídák’s multiple comparisons test.

These findings suggest that impaired LAD detachment may compromise genome stability by hindering the efficient repair of DSBs at the nuclear periphery.

To further assess genomic instability, we measured translocation frequency involving chromosomes 7 and 9 under DpnI-expressing conditions. As expected, DSB induction led to an increase in translocation frequency (Fig. 5b, g, h). However, despite the substantial rise in chromosomal breaks observed upon ATM inhibition and co-depletion of SUN1 and SUN2, we detected a marked decrease in translocation events (Fig. 5g, h). These findings suggest that while peripheral DSBs accumulate due to defective repair, they may be spatially or mechanistically restricted from engaging in aberrant chromosomal rearrangements.

While the persistent sequestration of unrepaired DSBs within LADs at the nuclear periphery appears to limit chromosomal translocations by spatially isolating damaged DNA ends, we investigated whether this mechanism might introduce other genomic and cellular vulnerabilities. Strikingly, we found that inducing DSBs within LADs under DpnI-expressing conditions led to a significant increase in the formation of micronuclei (MNi), as well as micronuclei-like or nuclear bleb structures, containing LAD-derived sequences (Fig. 5i and Extended Data Fig. 9e, f). This phenomenon was further amplified by ATM inhibition (Fig. 5i and Extended Data Fig. 9f) or enforced tethering that prevented LAD detachment (Fig. 5j, k and Extended Data Fig. 9g) and was specific to DNA damage induction (Extended Data Fig. 9h, i). Remarkably, the induction of a single DSB within a LAD using Cas9 was sufficient to reproduce these effects, reinforcing the conclusion that DNA breaks within LADs promote the generation of aberrant nuclear structures (Fig. 5l, m and Extended Data Fig. 9j). In this experimental system, the resulting MNi/blebs frequently contained the genomic region surrounding the targeted LAD (Fig. 5l, m), as analysed by FISH, and their occurrence also increased substantially upon ATM inhibition (Fig. 5m and Extended Data Fig. 9j), without a visible effect on the cell cycle distribution (Extended Data Fig. 9l). Intriguingly, in SUN1 and SUN2-depleted cells, undetached LADs are frequently localized within nuclear envelope blebs or MNi (Fig. 5n and Extended Data Fig. 9k), suggesting that LAD detachment serves a protective role in preventing nuclear envelope deformation in response to DSBs. These findings indicate that while defective LAD dynamics upon damage protects against translocations, it may alternatively lead to the generation of abnormal nuclear structures with potential consequences on genome integrity.

ATR (ataxia telangiectasia and Rad3-related), another DNA damage response kinase, has previously been implicated in sensing mechanical stress and promoting nuclear envelope rupture. To investigate whether the increased blebbing observed following ATM inhibition was specific to ATM inhibition, or whether ATR inhibition similarly phenocopied the effects observed after induction of a DSB at the LAD, we first quantified the number of MNi/blebs containing the LAD by immunoFISH. Interestingly, we observed a similar increase in LAD-containing MNi/blebs upon ATR inhibition (Extended data Fig. 9m-o). The detection of LAD signals outside the primary nucleus, within structures resembling nuclear blebs or MNi, prompted us to differentiate between these two possibilities. We performed live-cell microscopy to track the dynamic formation of these structures and assess their respective contributions to the observed phenotype. For this, we used RPE-1 cells expressing GFP-H2B and RFP-NLS to visualize nuclear shape and integrity and SNAP-MDC1 to visualise sensing of DSBs^26^ (Supplementary Table 1).

While MNi formation was observed in sgLAD-transfected cells and exacerbated by ATM inhibition (Fig. 6a, b, Supplementary Videos 1,2), the micronucleation levels were comparable with their sgCtrl-transfected counterparts (Fig. 6b), suggesting that LAD-specific DSB generation is not contributing to MNi formation. In contrast, nuclear shape evaluation showed a significant increase of nuclear blebs and other deformations such as nuclear lobulations following LAD targeting compared to control (Fig. 6a, c, d, Supplementary Videos 3-7). Intriguingly, in almost all cases formation of these nuclear abnormalities required cell transit through mitosis, since they were not present in the parent cell. Notably, ATM inhibition enhanced post-mitotic bleb formation, but not deformations (Fig. 6c, d) after LAD-specific DSB generation compared to matched controls.

**Figure 6:**
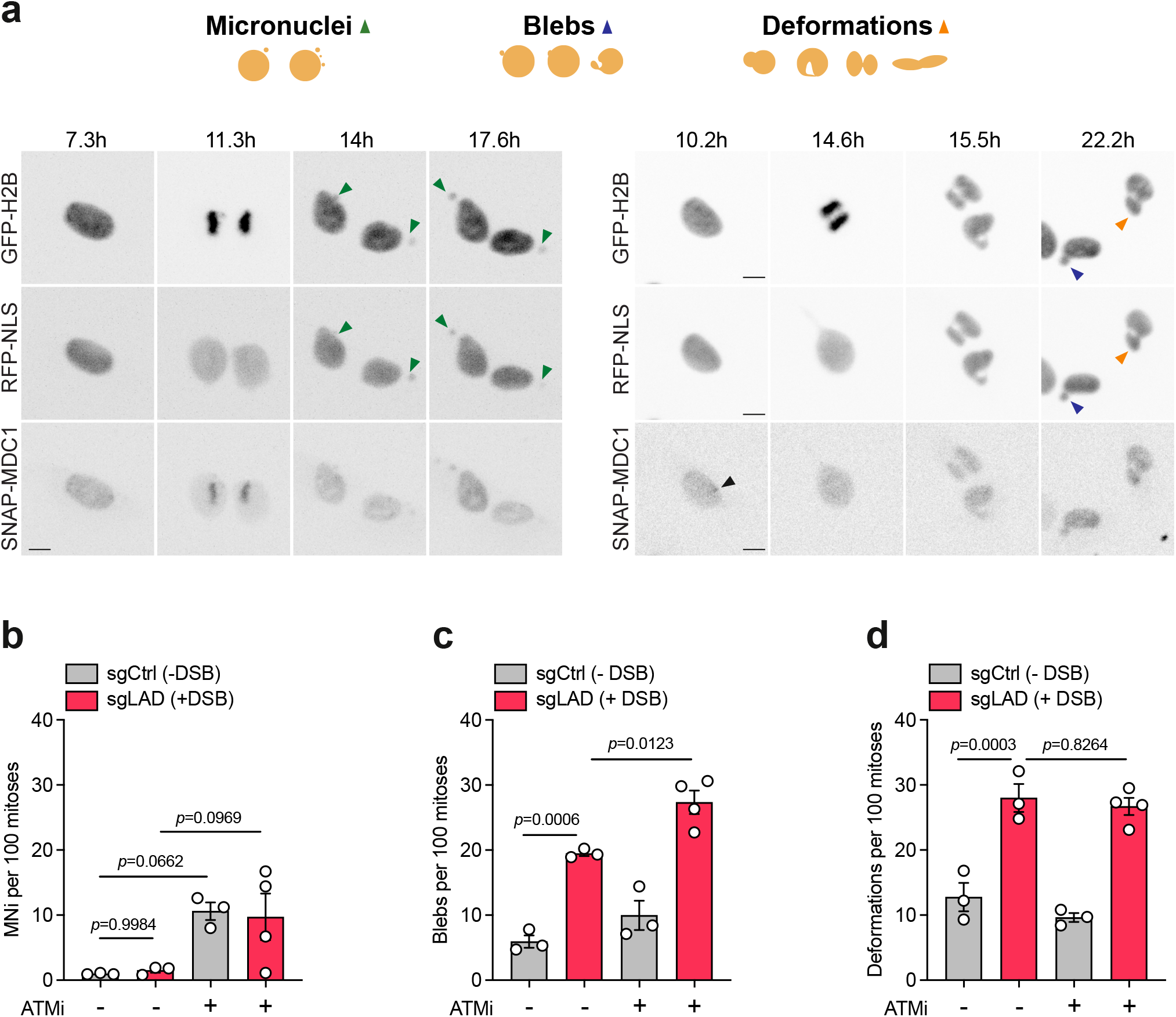
DNA damage at LADs results in nuclear deformations and the emergence of blebs, primarily after mitosis. a) Representative frames from live-cell imaging of RPE-1 cells expressing GFP-H2B, NLS-RFP, and SNAP-MDC1 transfected with Cas9 and sgLAD after ATMi treatment (10µM), highlighting post-mitotic micronucleus (MN) formation (left), and a bleb and a nuclear deformation (right). Black arrowheads indicate MDC1 foci, green arrowheads indicate MN, purple arrowheads indicate blebs, orange arrowheads indicate nuclear deformation. Scale bars 10 µm. b-d) Quantification of micronuclei (MNi) (b), blebs (c) and deformations (d) in one or both daughter cells per 100 mitoses in RPE-1 cells expressing GFP-H2B, NLS-RFP, and SNAP-MDC1 transfected with Cas9 and non-targeting control (sgCtrl) or sgLAD in control conditions (DMSO) or treated with ATMi (10µM). Data are presented as mean ± SEM from 3 or 4 biological replicates. Statistical comparisons were performed using two-way ANOVA followed by Šídák’s multiple comparisons test.

In contrast to ATM inhibition, we found that the increase in MNi/blebs observed after ATR inhibition was primarily due to a basal increase in MNi regardless of LAD targeting (Extended Data Figure 10a). Compared to DMSO-treated control, ATR inhibition led to no obvious increase in bleb formation nor deformations (Extended Data Figure 10b, c).

These findings suggest that ATM specifically modulates distinct nuclear abnormalities after mitosis, potentially shaping the outcome of chromatin repair and nuclear architecture during nuclear envelope reassembly.

Altogether, these results demonstrate that targeted lesions at LADs primarily lead to the formation of nuclear blebs, emerging mainly after mitosis. This implies a defect in nuclear envelope reassembly at damaged LAD sites, leading to the sequestration of unresolved DNA DSBs into bleb-like structures.

## Discussion

Our findings reveal that the timely release of LADs from the nuclear envelope following DNA damage acts as a critical protective response. This spatial reorganization enhances the DDR and facilitates efficient repair of DSBs originating from the nuclear periphery, thereby safeguarding genome stability. Interference with this process via ATM inhibition, depletion of LINC complex components (such as SUN1 or SUN2), or artificial chromatin tethering disrupts DSB repair kinetics, exacerbates nuclear envelope deformations and blebbing, and elevates chromosomal instability.

These observations support a model wherein ATM -signaling at damaged chromatin around LADs is transmitted to the cytoskeleton through the LINC complex. This relay promotes transient detachment and coordinated re-attachment of LADs, preserving both nuclear architecture and the integrity of compromised chromatin (see Extended Data Fig. 10d). When this signaling or relay is impaired, damaged LADs remain abnormally tethered, leading to their extrusion through nuclear blebs and contributing to genomic instability.

Our findings indicate that damage-induced detachment of LADs from the nuclear periphery is not accompanied by their tethering to nuclear pores. This represents a distinct response that does not recapitulate the nuclear pore relocalization of heterochromatin previously reported in *Drosophila*^27^. Our data emphasize a dynamic reconfiguration of genome architecture in response to genotoxic stress, highlighting the role of nuclear mechanics in maintaining genomic integrity. Echoing previous studies showing that DSBs in transcriptionally active regions often migrate to the nuclear periphery via directed movement or nuclear envelope invaginations^7–9^, we find that cytoskeleton-dependent mobility similarly aids in the repair of LAD-associated DSBs. DSBs at open chromatin often associate with the NPC, where HR is preferentially employed, reducing erroneous repair outcomes. In contrast, our results suggest in accordance with the literature on mechanical tension of the nuclear envelope, that LAD detachment could contribute to reducing mechanical stress, thereby limiting mislocalization of damaged DNA into blebs. LADs undergoing chromatin remodeling or transcriptional activation^28,29^ also show reduced lamina association, indicating a broader mechanism underlying detachment.

Cancer cells often exhibit nuclear envelope defects—such as blebbing and rupture—concurrent with genomic instability, both key features of malignant transformation^30^. Yet, the molecular connection between persistent DNA damage and nuclear architecture disruption has remained elusive. We show that the buildup of unrepaired lesions at LADs compromises chromatin–lamina tethering, triggering nuclear envelope blebbing. This phenotype mainly manifests after mitosis, indicating that damage at LADs interferes with nuclear envelope reassembly.

Interestingly, we observe that although ATM inhibition increases the number of unrepaired DNA ends, these breaks do not lead to a corresponding rise in translocations. One possible explanation is that impaired mobility or failure to resolve LAD-associated breaks prevents their illegitimate fusion with distant genomic loci, thereby limiting translocation formation despite elevated DNA damage. This indicates that the relocation of damaged chromatin away from the nuclear periphery must be both transient and precisely regulated. When this process is disrupted, while the persistent sequestration of breaks at the periphery may paradoxically prevent harmful rearrangements, it imposes significant mechanical stress on the nucleus. Collectively, our findings uncover a direct mechanistic link between disrupted chromatin–nuclear envelope interactions and the structural abnormalities commonly observed in cancer. Strikingly, cancer-associated single-nucleotide variants are enriched in genomic loci that locate at the nuclear periphery and decrease toward the center, suggesting that peripheral chromatin is more susceptible to somatic mutations^31^.

How does DNA damage lead to the detachment of LADs from the nuclear envelope? Our data indicate that DNA damage reduces the levels of LBR, which, together with lamin A, is known to anchor LADs to the nuclear envelope^32^. This reduction may contribute to the release of chromatin from the nuclear periphery. Notably, LBR contains a Tudor domain that binds modified histones, supporting a model in which LBR mediates chromatin tethering through direct histone interactions^33–35^. One possible mechanism is that, following DNA damage, 53BP1 competes with LBR for binding to shared histone marks through its Tudor domain, thereby weakening LBR-mediated LAD anchoring. However, the enhanced LAD detachment observed in LBR-depleted cells after DNA damage indicates that this mechanism alone is insufficient to account for the full extent of LAD release. Instead, these findings suggest that LBR functions as part of a broader network of chromatin–lamina tethering mechanisms, with additional anchoring pathways also being disrupted during the DNA damage response.

We envision two, not mutually exclusive, mechanisms by which ATM could couple damaged LADs to the cytoskeleton. ATM may alter chromatin mechanics, with these changes sensed and transmitted to the cytoskeleton through the LINC complex. Alternatively, ATM may directly coordinate chromatin and cytoskeletal remodeling by phosphorylating key components on both sides of the nuclear envelope. In agreement with the first, DNA damage in yeast leads to increased chromatin stiffness in an H2AX-dependent manner^36^ and global DNA damage leads to changes in nuclear stiffness with increased softening in an ATM-dependent manner^37,38^. These findings raise the possibility that the LINC complex may respond to alterations in chromatin stiffness, as it has been reported to sense changes in nuclear and chromatin mechanics, thereby providing a potential mechanism linking DNA damage-induced chromatin remodeling to LAD repositioning. This aligns with the view that chromatin–lamina interactions are dynamic and influenced by local chromatin context. Notably, subtelomeric domains enriched in γH2AX exhibit a reduced propensity to associate with the nuclear envelope, suggesting that intrinsic chromatin features modulate tethering efficiency^39^. In the second scenario, ATM might alter the abundance of nuclear envelope components^40^ and cytoskeleton as reported before.

The differential response of SUN1 and SUN2 to DSBs (Fig. 4) occurring at LADs is intriguing. Although previous studies often describe SUN1 and SUN2 as functionally complementary^21^ or even redundant in conditions of DNA damage, our findings suggest they may play distinct roles in this context. SUN1 depletion leads to the detachment of LADs both in the absence and presence of DNA damage, suggesting that SUN1 plays a constitutive and active role in tethering LAD chromatin to the nuclear envelope. This is consistent with previous reports showing that SUN1 anchors DSBs induced in active genes to the nuclear periphery^8^. Upon SUN2 depletion, DSBs at LADs remain localized at the nuclear periphery. This may indicate that SUN2 is actively involved in the detachment of these lesions from the lamina, possibly through a conformational change triggered by DNA damage. Alternatively, SUN1 may mediate their retention at the periphery in the absence of SUN2.

Interestingly, SUN2 has been shown to be targeted for proteasomal degradation via a mechanism involving Casein Kinase 2–dependent phosphorylation and recognition by the ubiquitin ligase Skp/Cullin1/F-BoxβTrCP (SCFβTrCP). This process is counteracted by the phosphatase CTDNEP1, which limits SUN2 ubiquitination^41^. Once ubiquitinated, SUN2 is extracted from the inner nuclear membrane by the AAA+ ATPase p97 and delivered to the proteasome. Notably, the accumulation of non-degradable SUN2 leads to aberrant nuclear morphology, increased sensitivity to DNA damage, and chromosomal instability during mitosis^41^. These findings are in line with the moderate decrease we observe after DNA damage and suggest that SUN2 turnover may represent a critical mechanism for modulating chromatin–lamina interactions in response to DNA damage.

Notably, reorganization of the LINC complex, decreased levels of lamin B1 and LBR, and detachment of LADs closely resemble phenotypes observed in premature aging conditions such as Hutchinson-Gilford Progeria Syndrome^42,43^, suggesting a potential link between DNA damage responses at LADs and aging-related nuclear defects^44–46^. Moreover, senescence is accompanied by reorganization of peripheral heterochromatin, indicative of LAD detachment, and an altered nuclear lamina, with loss of lamin B1^47^. Given that persistent DNA damage is a hallmark of senescence^48^, it is tempting to speculate that the progressive loss of LAD localization during senescence arises from the accumulation of DNA damage within these domains. Finally, endogenous DNA breaks at promoters and other gene regulatory elements driven by torsional stress and topoisomerase activity^49^ may similarly trigger local ATM activation, potentially initiating LAD detachment and chromatin reorganization during cellular differentiation, thereby highlighting the broader significance of our findings.

## Acknowledgements

We thank Yan Gu for help with the image analysis and microscopy and Alex Herbert for developing a plugin in imageJ to analyse the LADs signal from the NUP153 to the center of the cell. We are grateful to B. van Steensel and D. Durocher for the gifts of Shield1-inducible Dam-Lamin B1 clone11 (HT1080) and hTERT RPE-1 Cas9 cell lines, to Chris Chan for the plasmid pMito-mCherry-FRB_IRES_FKBP(III)-GBPen (Addgene #128268) and the 53BP1-deficient cells, and to Steve Jackson for H2AX KO/KI cell lines.

This work in the E.S lab was supported by the Medical Research Council (MR/X000818/1) and European Research Council (ERC) Advanced grant (ChromoSENSOR)/UK guarantee funding from EPSRC (EP/Y027124/1). The microscopy facility at the Genome Center is supported by the Wolfson Foundation. The Spinning Disk Confocal System (VisiScope 5-Elements, IMB Microscopy Core Facility) was supported by the Deutsche Forschungsgemeinschaft (INST 247/912-1FUGG). Work in the S.J.B. lab is supported by the Francis Crick Institute (CC2098), European Research Council Advanced Investigator grants (TelMetab, ChrEndProt), a Wellcome Trust Senior Investigator Award, and CRUK RadNet City of London. Work in the S.P. lab is supported by the European Research council (ERC) Starting grant (MitErrA) and the Rising Star Award from EACR, the Mark and Pezcoller foundations. R.A.L. is supported by a Swiss National Science Foundation fellowship.

## Extended Data

### Extended Data figure legends

**Extended Data Figure 1.**
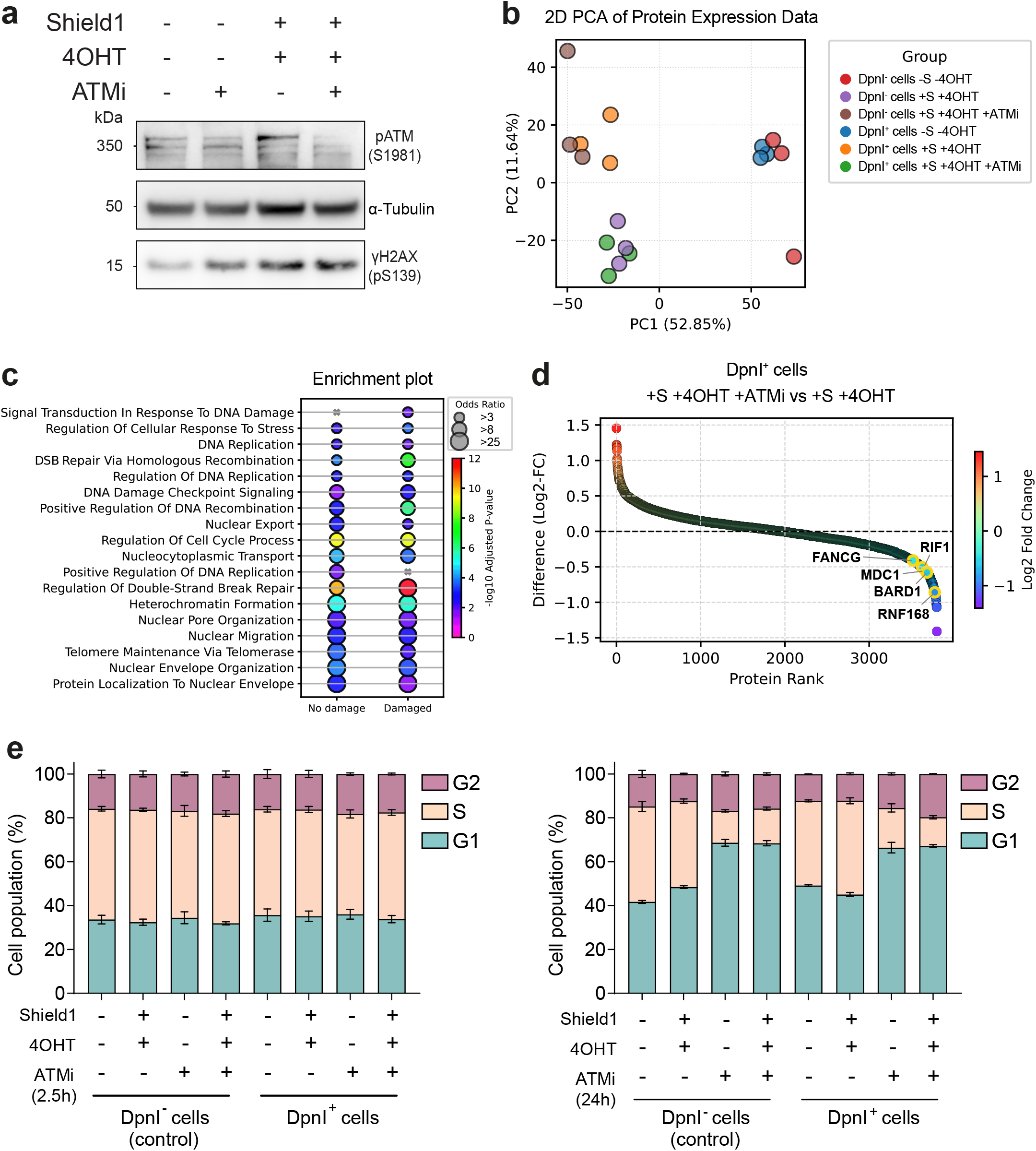
a) Immunoblot of phosphorylation of ATM (pATM) and phospho-S139 (pS139) of H2AX (γH2AX) in DpnI expressing cells in the absence and presence of Shield1 and 4OHT. Cells were treated with the ATM inhibitor (ATMi; 10µM) or mock (DMSO). alpha-tubulin was used as loading control. b) Principal component analysis of cells expressing (DpnI^+^) or not (DpnI^-^) DpnI, with or without Shield1 (S) and 4OHT and treated with ATMi (10µM) or mock (DMSO). Conditions show tight clustering of biological replicates and clear separation between treatment groups, reflecting high reproducibility and distinct proteomic responses. c) Functional enrichment analysis of lamina-associated proteins under basal and DNA damage conditions. Dot plot showing representative GO terms enriched in cells expressing or not DpnI and treated with Shield1 and 4OHT. (DpnI^-^ cells +S +4OHT = No damage) and (DpnI^+^ cells +S +4OHT = Damaged). Enrichment was performed using Fisher’s exact test; selected terms illustrate functional differences between basal and DNA damage-associated chromatin proteomes. d) ATM inhibition abrogates DpnI-induced recruitment of DNA repair factors. Rank-ordered plot of protein abundance differences in DpnI^+^ cells +S +4OHT treated with either DMSO or ATMi (10µM). Proteins are color-coded by log₂ fold change, with selected DNA damage response factors highlighted and annotated. Loss of enrichment under ATMi treatment indicates that recruitment of these factors to chromatin is ATM-dependent. e) Flow cytometry analysis of DpnI cells (Dpn1^−^ and Dpn1^+^) treated with Shield1 and 4OHT. ATMi was added 30 min prior to 4OHT treatment, with total ATM inhibition lasting for either 2.5 h (*left*) or 24 h (*right*). DNA synthesis and total DNA content were assessed by BrdU incorporation and propidium iodide staining. Data are presented as the mean percentage of cells in each phase of the cell cycle ± SEM from 3 biological replicates on the left panel and two on the right panel.

**Extended Data Figure 2.**
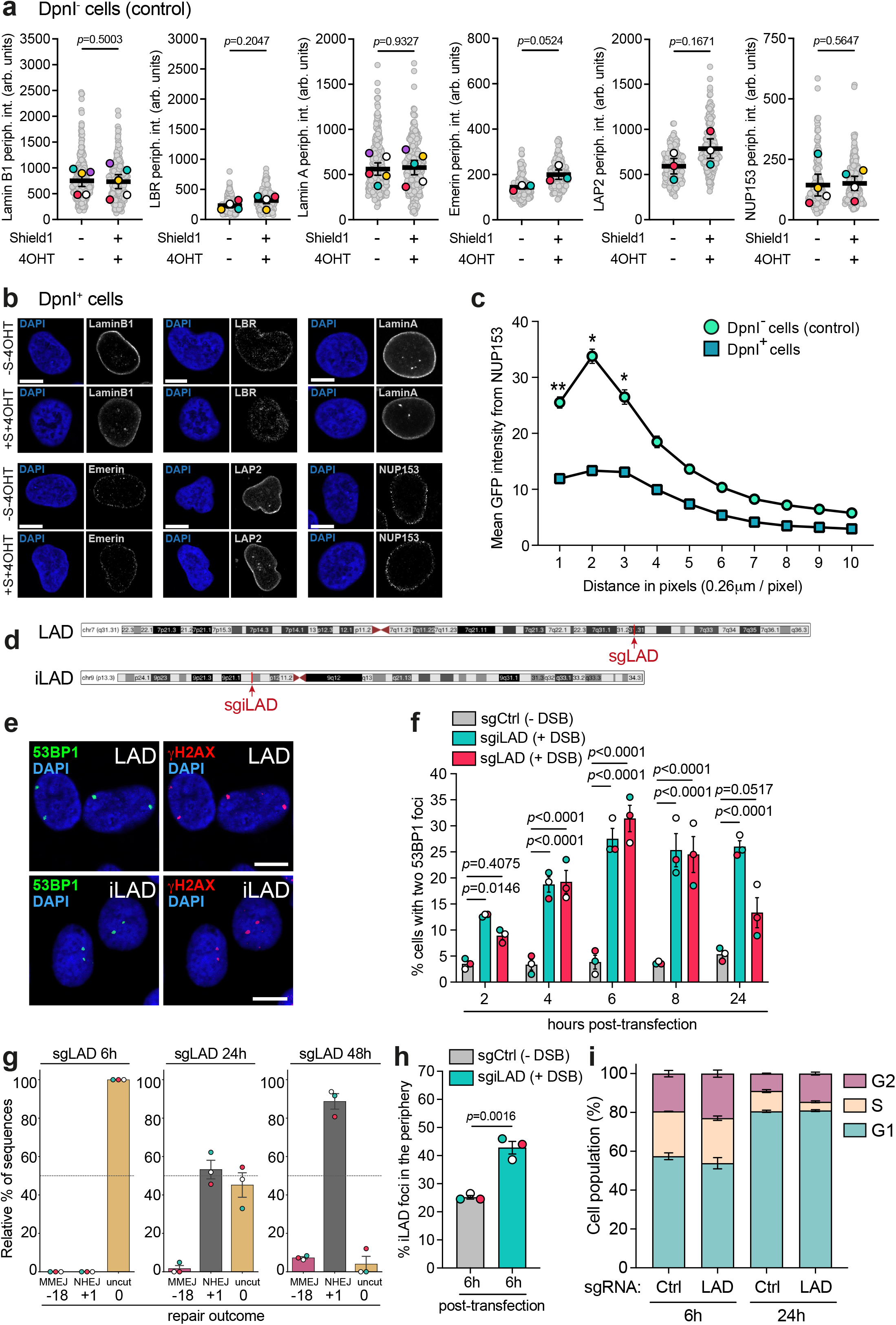
a) Quantification in arbitrary units of lamin B1, LBR, lamin A, emerin, LAP2 and NUP153 at the nuclear periphery in cells which do not express DpnI treated or not with Shield1 and 4OHT. Data represent the mean of at least 3 independent biological replicates. Statistical comparisons were performed using a ratio paired *t-*test. b) Representative confocal images of lamin B1, LBR, lamin A in DpnI expressing cells treated or not with Shield1 and 4OHT. Scale bar 10µm. c) Quantification of mean GFP intensity along a 10 × 3-pixel conical spatial gradient extending from each NUP153-marked nuclear pore focus toward the nuclear interior. Cells expressing or not DpnI were analyzed. Data represent mean ± SEM from 3 biological replicates. Statistical comparisons were performed using a two-way ANOVA followed by Šídák’s multiple comparisons test between DpnI⁺ and DpnI⁻ conditions at each corresponding spatial distance. Significant statistics, position at pixel 1 ** *p*=0.0027, position at pixel 2 * *p*=0.0256 and position at pixel 3 * *p*=0.0461. d) Indication of the genomic location (red arrow) in chromosome 7 (sgLAD) and 9 (sgiLAD) where the single DSB is induced. e) Representative confocal images of RPE-1 cells expressing Cas9 6 h post-transfection with a single guide RNA which targets a LAD (sgLAD, *top*) and an iLAD (sgiLAD, *bottom*). Induction of DSBs is shown by the 53BP1 (green) and γH2AX (red) foci. The nucleus is stained with DAPI (blue). Scale bar 10µm. f) Percentage of RPE-1 cells expressing Cas9 and transfected with non-targeting control (sgCtrl), sgiLAD, or sgLAD containing two 53BP1 foci 2, 4, 6, 8 and 24 hours post transfection. Data are presented as mean ± SEM from 3 biological replicates. Statistical comparisons were performed using two-way ANOVA followed by Šídák’s multiple comparisons test. g) TIDE analysis showing the relative percentage of the types of significant indels (*p* < 0.001) accumulated and the kinetics of indel formation in RPE-1 cells expressing Cas9 and transfected sgLAD. Samples were collected and analyzed at 6-, 24-, and 48-hours post-transfection. Data represent the mean ± SEM percentage of repair outcomes of 3 biological replicates. h) Percentage of iLAD loci which colocalise with lamin B1 in RPE-1 cells expressing Cas9 and transfected with non-targeting control (sgCtrl) or sgiLAD for 6 h. Data are presented as mean ± SEM from 3 biological replicates. Statistical comparisons were performed using unpaired *t*-test. i) RPE-1 cells transfected with non-targeting control (sgCtrl) or sgLAD for 6 h and 24 h were stained for incorporated BrdU against total DNA content using propidium iodide. Data are presented as the mean percentage of cells in each phase of the cell cycle ± SEM from 3 biological replicates.

**Extended Data Figure 3.**
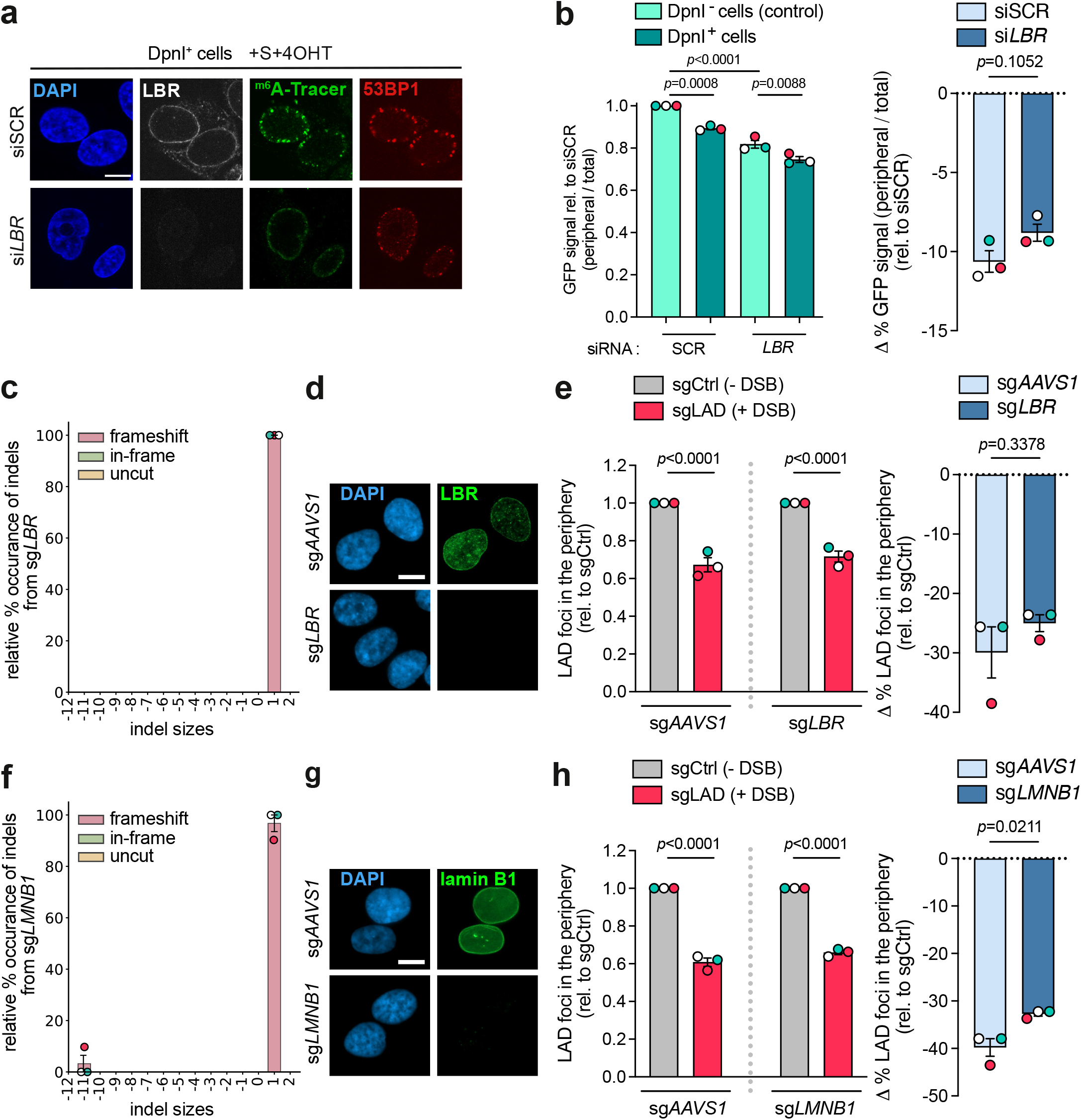
a) Representative immunofluorescence images showing 53BP1 foci (red), m6A-Tracer-GFP (green), and lamin B receptor (LBR) expression (grey) in mock depleted (siSCR) or *LBR* depleted (si*LBR*) cells expressing DpnI treated with Shield1 and 4OHT. DNA is counterstained with DAPI (blue). Scale bar 10 µm. b) *Left*, quantification of the relative abundance of m^6^A-Tracer at the nuclear rim in cells expressing or not DpnI, depleted using siRNA for *LBR* or control (SCR), treated with Shield1 and 4OHT. Data are presented as mean ± SEM from 3 biological replicates. Statistical comparisons were performed using two-way ANOVA followed by Šídák’s multiple comparisons test. *Right*, percentage change in abundance of m^6^A-Tracer in the nuclear periphery (DpnI+ - DpnI-) relative to the corresponding DpnI- condition. Data are presented as mean ± SEM from 3 biological replicates. Statistical comparisons were performed using unpaired *t*-test. c) TIDE analysis showing the relative percentage occurrence of significant indels (*p* < 0.001) accumulated 5 days post transfection with the sgRNA targeting *LBR* for depletion. Indels resulting in a frameshift mutation are coloured in pink, indels retaining the frame (in-frame) in green, and “uncut” sequences in yellow. Data are presented as mean ± SEM from 3 biological replicates. d) Representative images depicting LBR expression (green) in mock-depleted (sg*AAVS1*) or LBR-depleted (sg*LBR*) RPE-1 cells expressing Cas9. DNA is stained with DAPI (blue). Scale bar 10µm. e) *Left*, quantification of LAD foci which colocalise with lamin B1 in mock-depleted (sg*AAVS1*) or LBR-depleted (sg*LBR*) cells transfected with either non-targeting control guide (sgCtrl) or sgLAD for 6 h. Data are presented as mean ± SEM from 3 biological replicates. Statistical comparisons were performed using two-way ANOVA followed by Šídák’s multiple comparisons test. *Right*, percentage change in LAD foci in the nuclear periphery (sgLAD - sgCtrl) relative to the corresponding sgCtrl condition. Data are presented as mean ± SEM from 3 biological replicates. Statistical comparisons were performed using unpaired *t*-test. f) TIDE analysis showing the relative percentage occurrence of significant indels (*p* < 0.001) accumulated 5 days post transfection with the sgRNA targeting *LMNB1* for depletion. Indels resulting in a frameshift mutation are coloured in pink, indels retaining the frame (in-frame) in green, and “uncut” sequences in yellow. Data are presented as mean ± SEM from 3 biological replicates. g) Representative images depicting lamin B1 expression (green) in mock-depleted (sg*AAVS1*) or LMNB1-depleted (sg*LMNB1*) RPE-1 cells expressing Cas9. DNA is stained with DAPI (blue). Scale bar 10µm. h) *Left*, quantification of LAD foci which colocalise with lamin A in mock-depleted (sg*AAVS1*) or LMNB1-depleted (sg*LMNB1*) cells transfected with either non-targeting control guide (sgCtrl) or sgLAD for 6 h. Data are presented as mean ± SEM from 3 biological replicates. Statistical comparisons were performed using two-way ANOVA followed by Šídák’s multiple comparisons test. *Right*, percentage change in LAD foci in the nuclear periphery (sgLAD - sgCtrl) relative to the corresponding sgCtrl condition. Data are presented as mean ± SEM from 3 biological replicates. Statistical comparisons were performed using unpaired *t*-test.

**Extended Data Figure 4.**
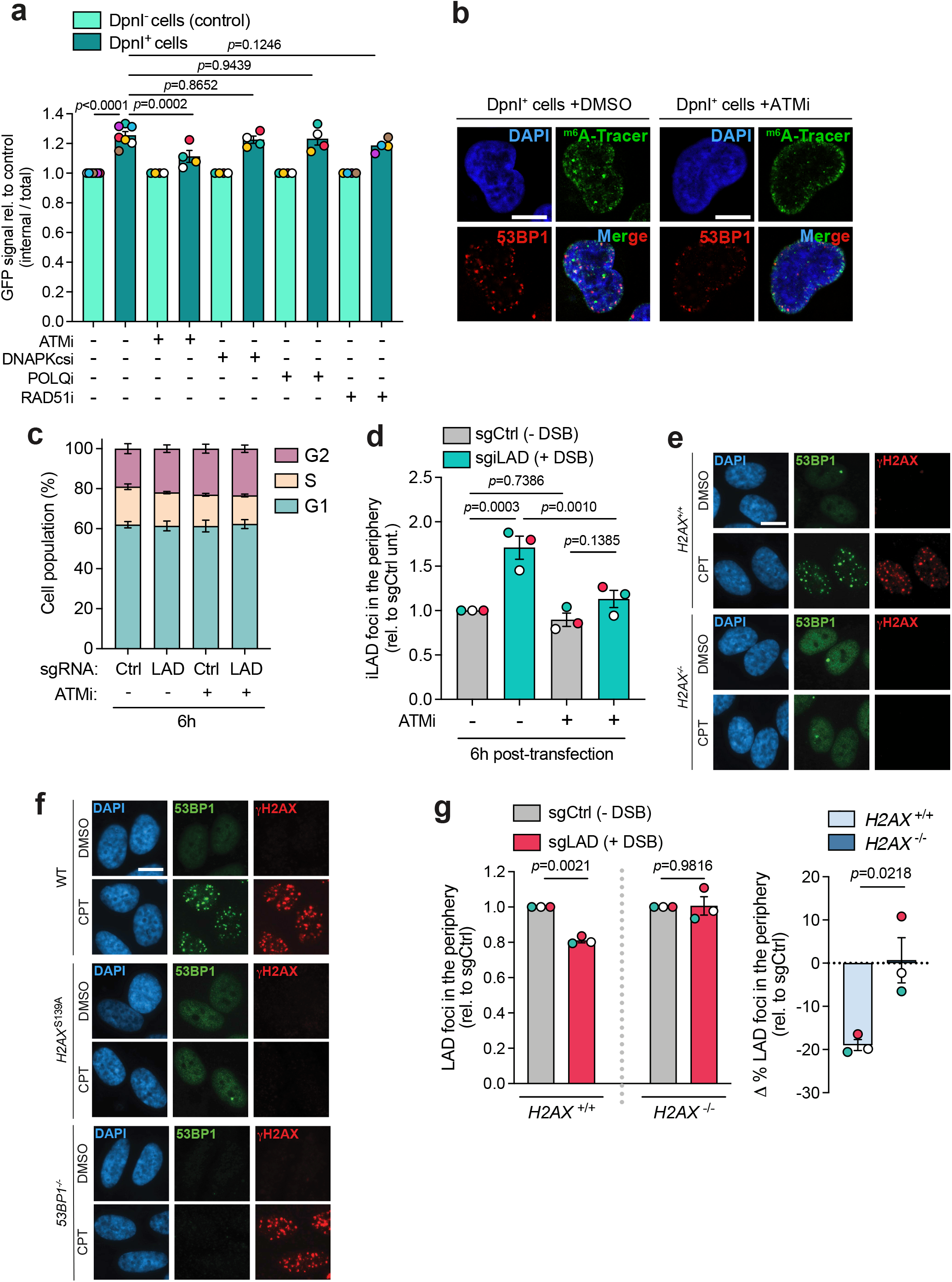
a) Quantification of the relative abundance of m^6^A-Tracer at the nuclear interior in cells expressing or not DpnI, treated with Shield1 and 4OHT in control conditions (DMSO) or treated with inhibitors for ATM (ATMi, 10µM), DNAPKcs (DNAPKi, 10µM) and Pol theta (PolQi, 10µM) and RAD51i (20µM). Data are presented as mean ± SEM from 3 biological replicates. Statistical comparisons were performed using two-way ANOVA followed by Šídák’s multiple comparisons test. b) Representative confocal images of m^6^A-Tracer (green) and 53BP1 (red) in DpnI expressing cells treated with Shield1 and 4OHT in control conditions (DMSO) or treated with ATMi (10µM). Scale bar 10µm. c) RPE-1 cells transfected with non-targeting control guide (sgCtrl) or sgLAD for 6 h treated with ATMi (10µM) or DMSO (-) were stained for incorporated BrdU against total DNA content using propidium iodide. Data are presented as the mean percentage of cells in each phase of the cell cycle ± SEM from 3 biological replicates. d) Quantification of iLAD loci which colocalise with lamin B1 in RPE-1 cells expressing Cas9 and transfected with non-targeting control (sgCtrl) or sgiLAD for 6 h treated with ATMi (10µM) or DMSO (-). Data are presented as mean ± SEM from 3 biological replicates. Statistical comparisons were performed using two-way ANOVA followed by Šídák’s multiple comparisons test. e) Representative images depicting 53BP1 (green) and γH2AX (red) in WT (*H2AX*^+/+^) and H2AX-deficient (*H2AX*^-/-^) RPE-1 cells treated with 1h CPT (25µM) or DMSO. DNA is stained with DAPI (blue). Scale bar 10µm. f) Representative images depicting 53BP1 (green) and γH2AX (red) in WT, *H2AX*^S139^ mutant and 53BP1-depleted (*53BP1*^-/-^) RPE-1 cells treated with 1h CPT (25µM) or DMSO. DNA is stained with DAPI (blue). Scale bar 10µm. g) *Left*, quantification of LAD foci which colocalise with lamin B1 in WT (*H2AX*^+/+^) and H2AX-depleted (*H2AX*^-/-^) RPE-1 cells transfected with Cas9 and either non-targeting control guide (sgCtrl) or sgLAD for 7 h. Data are presented as mean ± SEM from 3 biological replicates. Statistical comparisons were performed using two-way ANOVA followed by Šídák’s multiple comparisons test. *Right*, percentage change in LAD foci in the nuclear periphery (sgLAD - sgCtrl) relative to the corresponding sgCtrl condition. Data are presented as mean ± SEM from 3 biological replicates. Statistical comparisons were performed using unpaired *t*-test.

**Extended Data Figure 5.**
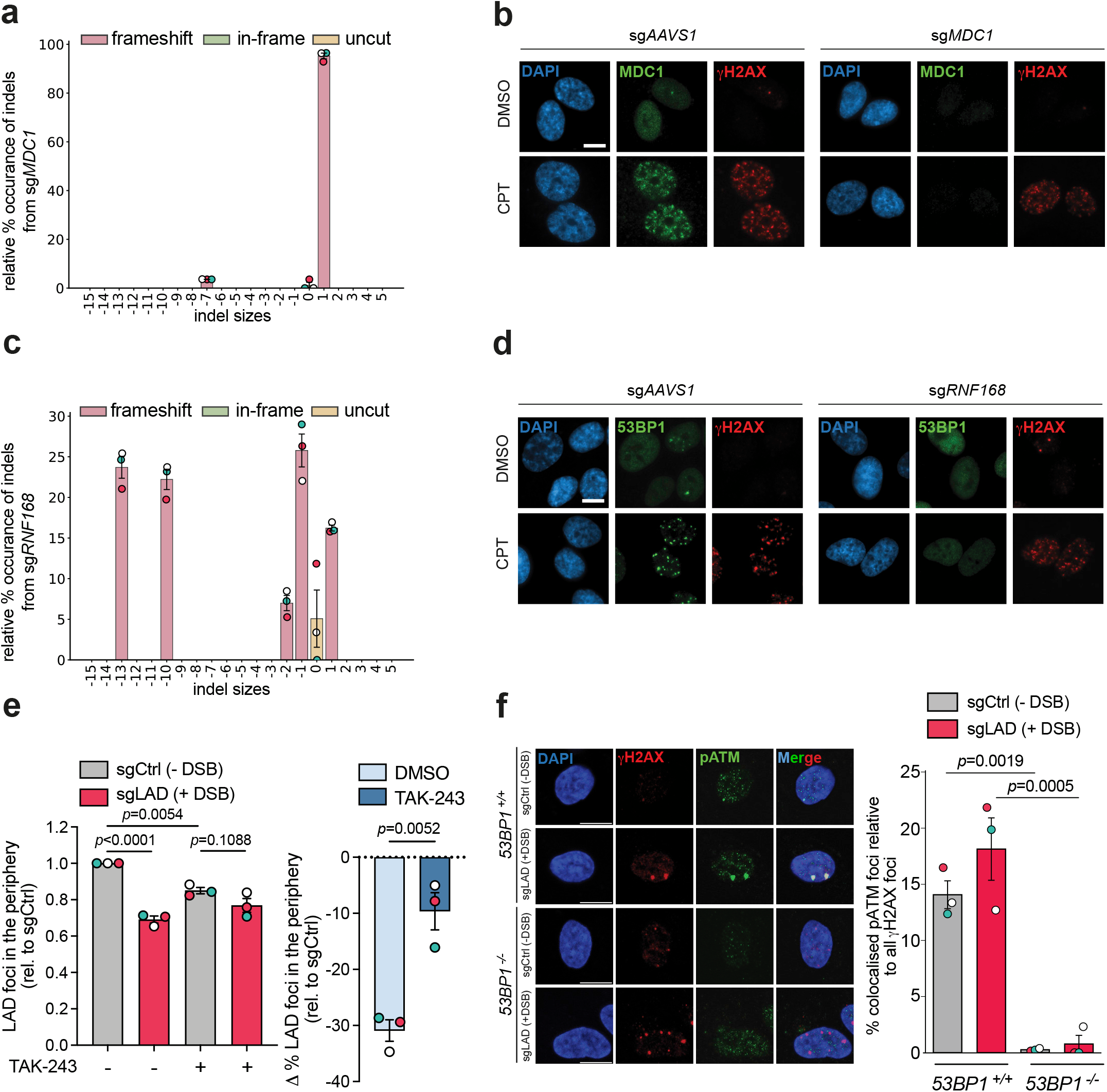
a) TIDE analysis showing the relative percentage occurrence of significant indels (*p* < 0.001) accumulated 5 days post transfection with the sgRNA targeting *MDC1* for depletion. Indels resulting in a frameshift mutation are coloured in pink, indels retaining the frame (in-frame) in green, and “uncut” sequences in yellow. Data are presented as mean ± SEM from 3 biological replicates. b) Representative images depicting 53BP1 (green) and γH2AX (red) in mock-depleted (sg*AAVS1*) or MDC1-depleted (sg*MDC1*) RPE-1 cells expressing Cas9 treated with 1h CPT (25µM) or DMSO. DNA is stained with DAPI (blue). Scale bar 10µm. c) TIDE analysis showing the relative percentage occurrence of significant indels (*p* < 0.001) accumulated 5 days post transfection with the sgRNA targeting *RNF168* for depletion. Indels resulting in a frameshift mutation are coloured in pink, indels retaining the frame (in-frame) in green, and “uncut” sequences in yellow. Data are presented as mean ± SEM from 3 biological replicates. d) Representative images depicting 53BP1 (green) and γH2AX (red) in mock-depleted (sg*AAVS1*) or RNF168-depleted (sg*RNF168*) RPE-1 cells expressing Cas9 treated with 1h CPT (25µM) or DMSO. DNA is stained with DAPI (blue). Scale bar 10µm. e) *Left*, quantification of LAD foci which colocalise with lamin B1 in cells transfected with either non-targeting control guide (sgCtrl) or sgLAD for 6 h treated with the ubiquitin inhibitor TAK-243 (5µM) or DMSO (-). Data are presented as mean ± SEM from 3 biological replicates. Statistical comparisons were performed using two-way ANOVA followed by Šídák’s multiple comparisons test. *Right*, percentage change in LAD foci in the nuclear periphery (sgLAD - sgCtrl) relative to the corresponding sgCtrl condition. Data are presented as mean ± SEM from 3 biological replicates. Statistical comparisons were performed using unpaired *t*-test. f) *Left*, representative images depicting γH2AX (red) and pATM (green) in WT (*53BP1*^+/+^) and 53BP1-deficient (*53BP1*^-/-^) RPE-1 cells transfected with either non-targeting control guide (sgCtrl) or sgLAD for 7 h. DNA is stained with DAPI (blue). Scale bars are 20µm. *Right*, quantification of the percentage of pATM foci which colocalised with γH2AX relative to all quantified γH2AX in 100 cells for each condition. Data are presented as mean ± SEM from 3 biological replicates. Statistical comparison was performed using two-way ANOVA followed by Šídák’s multiple comparisons tests.

**Extended Data Figure 6.**
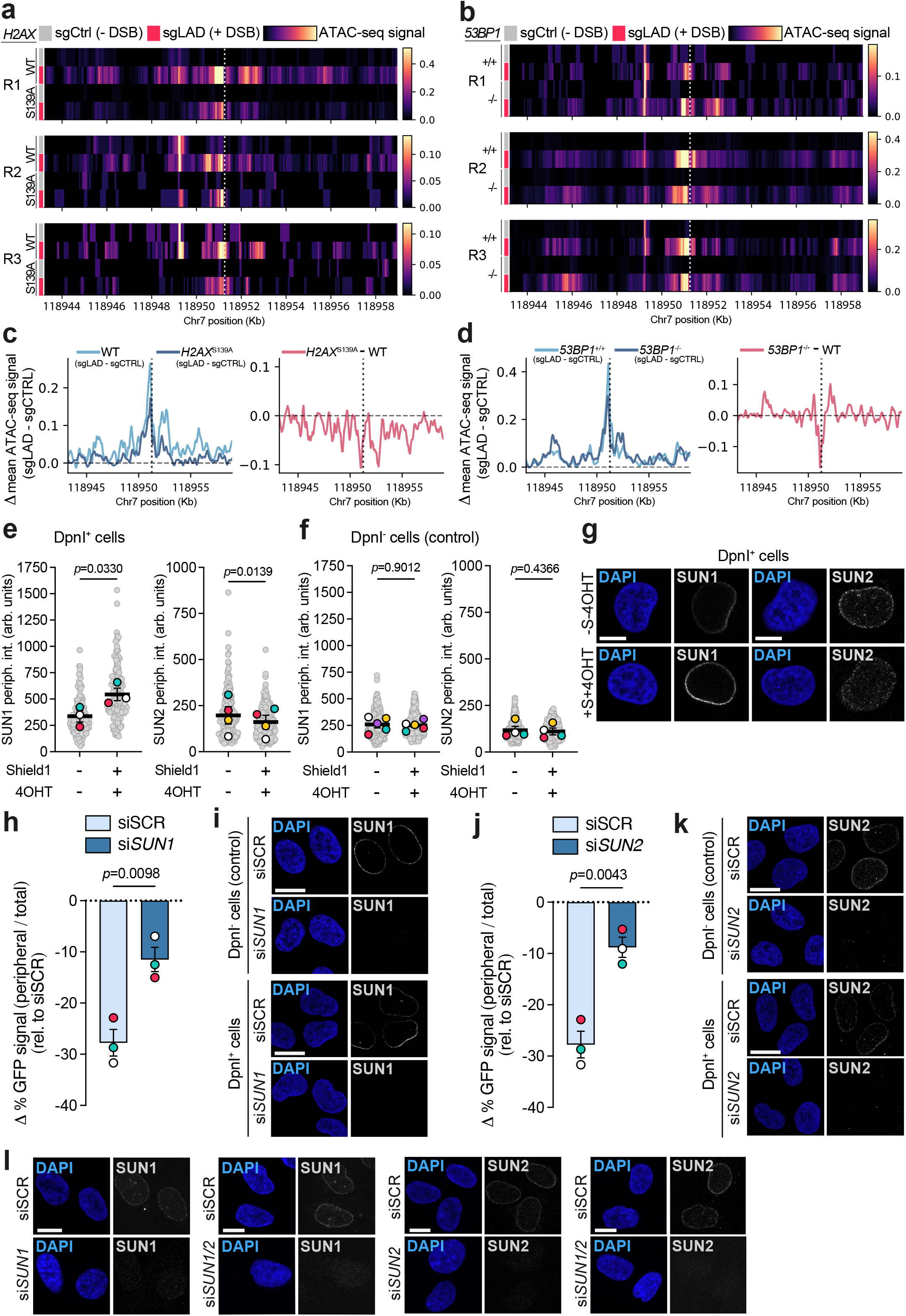
a) Heatmaps of ATAC-seq signal across a 30 Kb region on chromosome 7 in WT (*H2AX*^WT^) and *H2AX*^S139^ mutant RPE-1 cells transfected with either control guide (sgCtrl) or sgLAD for 7 h. The site of DSB induction is denoted by the dotted vertical lines. Three independent biological replicates for each condition are represented with corresponding scale bars for each replicate. b) Heatmaps of ATAC-seq signal across a 30 Kb region on chromosome 7 in WT (*53BP1*^+/+^) and 53BP1-deficient (*53BP1*^-/-^) RPE-1 cells transfected with either control guide (sgCtrl) or sgLAD for 7 h. The site of DSB induction is denoted by the dotted vertical lines. Three independent biological replicates for each condition are represented with corresponding scale bars for each replicate. c) WT (*H2AX*^WT^) and *H2AX*^S139^ mutant RPE-1 cells. *Right,* the mean difference in ATAC-seq signal between sgLAD and sgCtrl transfected sample tracks (mean Δ ATAC-seq signal) across a 15 Kb region on chromosome 7 with 200bp smoothing for each represented cell line. *Left,* The difference between the mean Δ ATAC-seq signal of WT and mutant cells is represented in pink, termed as the mean ΔΔ ATAC-seq signal. Vertical dotted lines denote the site of DSB induction. d) WT (*53BP1*^+/+^) and 53BP1-deficient (*53BP1*^-/-^) RPE-1 cells. *Right,* the mean difference in ATAC-seq signal between sgLAD and sgCtrl transfected sample tracks (mean Δ ATAC-seq signal) across a 15 Kb region on chromosome 7 with 200bp smoothing for each represented cell line. *Left,* the difference between the mean Δ ATAC-seq signal of WT and mutant cells is represented in pink, termed as the mean ΔΔ ATAC-seq signal. Vertical dotted lines denote the site of DSB induction. e) Quantification in arbitrary units (linear scale) of SUN1 and SUN2 at the nuclear periphery in DpnI expressing cells treated or not with Shield1 and 4OHT. Data are presented as mean ± SEM from 3 biological replicates. Statistical comparisons were performed using a ratio paired *t*-test. f) Quantification in arbitrary units of SUN1 and SUN2 at the nuclear periphery in DpnI negative cells treated or not with Shield1 and 4OHT. Data are presented as mean ± SEM from 3 biological replicates. Statistical comparisons were performed using a ratio paired *t*-test. g) Representative confocal images of SUN1 and SUN2 (white) in DpnI expressing cells treated or not with Shield1 and 4OHT. DNA is stained with DAPI (blue). Scale bar 10µm. h) Percentage change in abundance of m^6^A-Tracer in the nuclear periphery (DpnI^+^ - DpnI^-^) relative to the corresponding DpnI^-^ condition in cells expressing or not DpnI, depleted using siRNA for *SUN1* or control (SCR), treated with Shield1 and 4OHT. Data are presented as mean ± SEM from 3 biological replicates. Statistical comparisons were performed using unpaired *t*-test. i) Representative confocal images of SUN1 (white) in cells expressing or not DpnI, depleted using siRNA for *SUN1* or control (SCR). DNA is stained with DAPI (blue). Scale bar 10µm. j) Percentage change in abundance of m^6^A-Tracer in the nuclear periphery (DpnI+ - DpnI-) relative to the corresponding DpnI^-^ condition in cells expressing or not DpnI, depleted using siRNA for *SUN2* or control (SCR), treated with Shield1 and 4OHT. Data are presented as mean ± SEM from 3 biological replicates. Statistical comparisons were performed using unpaired *t*-test. k) Representative confocal images of SUN1 (white) in cells expressing or not DpnI, depleted using siRNA for *SUN2* or control (SCR). DNA is stained with DAPI (blue). Scale bar 10µm. l) Representative confocal images depicting SUN1 and SUN2 expression (white) in RPE-1 cells expressing Cas9 and transfected with siSCR, si*SUN1*, si*SUN2* or si*SUN1*/si*SUN2*. DNA is stained with DAPI (blue). Scale bar 10µm. Note: All six conditions (siSRC, si*SUN1*, and si*SUN2*, ±DSB) were processed in the same experiment. The siSRC control lanes shown in panels (h) and (j) derive from the same samples and are presented separately for clarity of comparison with each depletion.

**Extended Data Figure 7.**
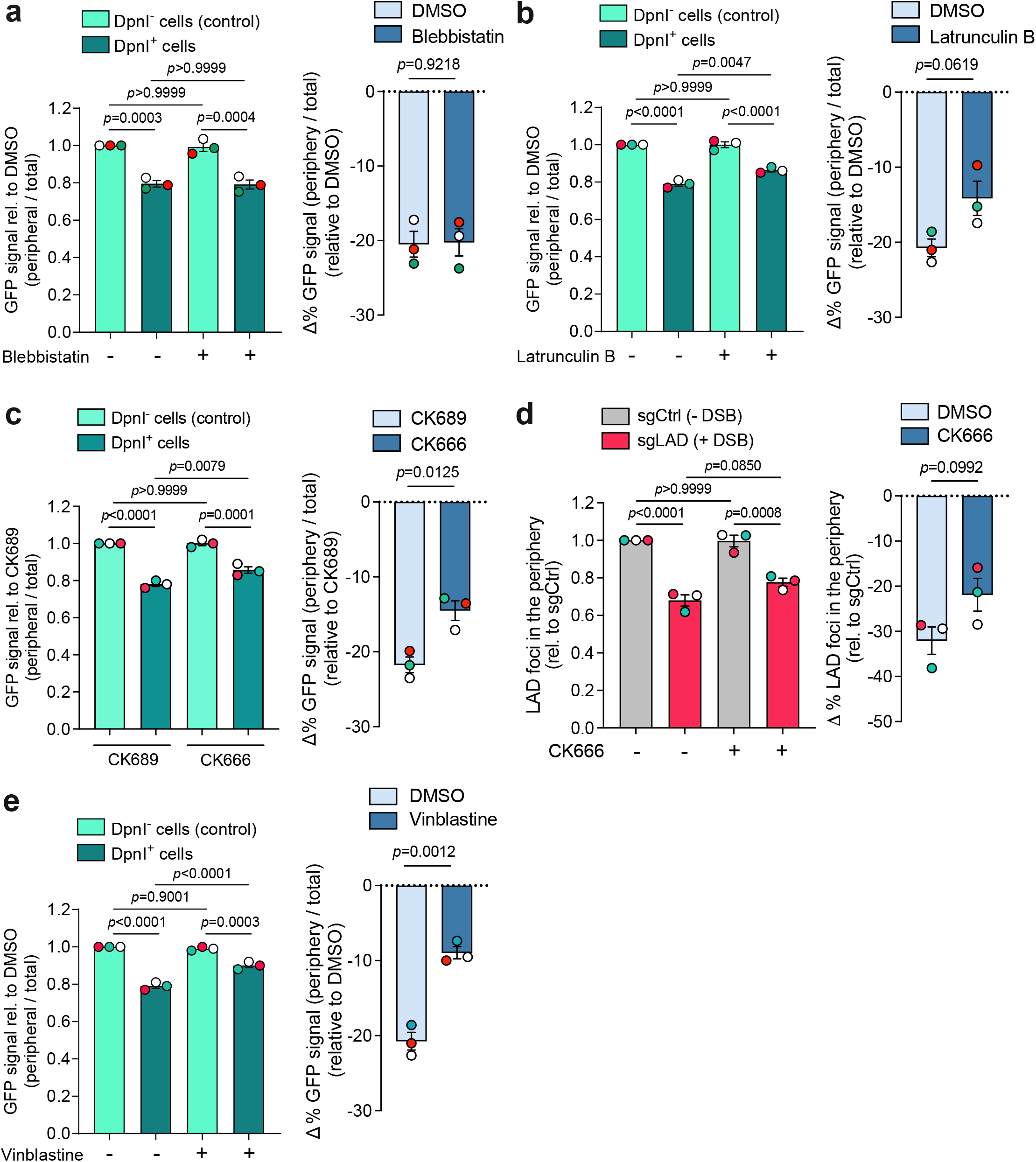
a) *Left*, quantification of the relative abundance of m^6^A-Tracer at the nuclear rim in cells expressing or not DpnI, treated with Shield1, 4OHT and DMSO or Blebbistatin (10μM) for 20 h. Data are presented as mean ± SEM from 3 biological replicates. Statistical comparisons were performed using two-way ANOVA followed by Šídák’s multiple comparisons test. *Right*, percentage change in abundance of m^6^A-Tracer in the nuclear periphery (DpnI^+^ - DpnI^-^) relative to the corresponding DpnI^-^ condition. Data are presented as mean ± SEM from 3 biological replicates. Statistical comparisons were performed using unpaired *t*-test. b) *Left*, quantification of the relative abundance of m^6^A-Tracer at the nuclear rim in cells expressing or not DpnI, treated with Shield1, 4OHT and DMSO or latrunculin B (100 nM) for 3 h. Data are presented as mean ± SEM from 3 biological replicates. Statistical comparisons were performed using two-way ANOVA followed by Šídák’s multiple comparisons test. *Right*, percentage change in abundance of m^6^A-Tracer in the nuclear periphery (DpnI^+^ - DpnI^-^) relative to the corresponding DpnI^-^ condition. Data are presented as mean ± SEM from 3 biological replicates. Statistical comparisons were performed using unpaired *t*-test. c) *Left*, quantification of the relative abundance of m^6^A-Tracer at the nuclear rim in cells expressing or not DpnI, treated with Shield1, 4OHT and mock control (CK689, 100 μM) or Arp2/3 (CK666, 100 μM) for 3 h. Data are presented as mean ± SEM from 3 biological replicates. Statistical comparisons were performed using two-way ANOVA followed by Šídák’s multiple comparisons test. *Right*, percentage change in abundance of m^6^A-Tracer in the nuclear periphery (DpnI^+^ - DpnI^-^) relative to the corresponding DpnI^-^ condition. Data are presented as mean ± SEM from 3 biological replicates. Statistical comparisons were performed using unpaired *t*-test. d) *Left*, quantification of LAD foci which colocalise with lamin B1 in cells transfected with either non-targeting control guide (sgCtrl) or sgLAD for 6 h treated with Arp2/3 (CK666, 100 μM) or DMSO (-). Data are presented as mean ± SEM from 3 biological replicates. Statistical comparisons were performed using two-way ANOVA followed by Šídák’s multiple comparisons test. *Right*, percentage change in LAD foci in the nuclear periphery (sgLAD - sgCtrl) relative to the corresponding sgCtrl condition. Data are presented as mean ± SEM from 3 biological replicates. Statistical comparisons were performed using unpaired *t*-test. e) *Left*, quantification of the relative abundance of m^6^A-Tracer at the nuclear rim in cells expressing or not DpnI, treated with Shield1, 4OHT and DMSO or Vinblastine (100nM) for 3 h. Data are presented as mean ± SEM from 3 biological replicates. Statistical comparisons were performed using two-way ANOVA followed by Šídák’s multiple comparisons test. *Right*, percentage change in abundance of m^6^A-Tracer in the nuclear periphery (DpnI^+^ - DpnI^-^) relative to the corresponding DpnI^-^ condition. Data are presented as mean ± SEM from 3 biological replicates. Statistical comparisons were performed using unpaired *t*-test.

**Extended Data Figure 8.**
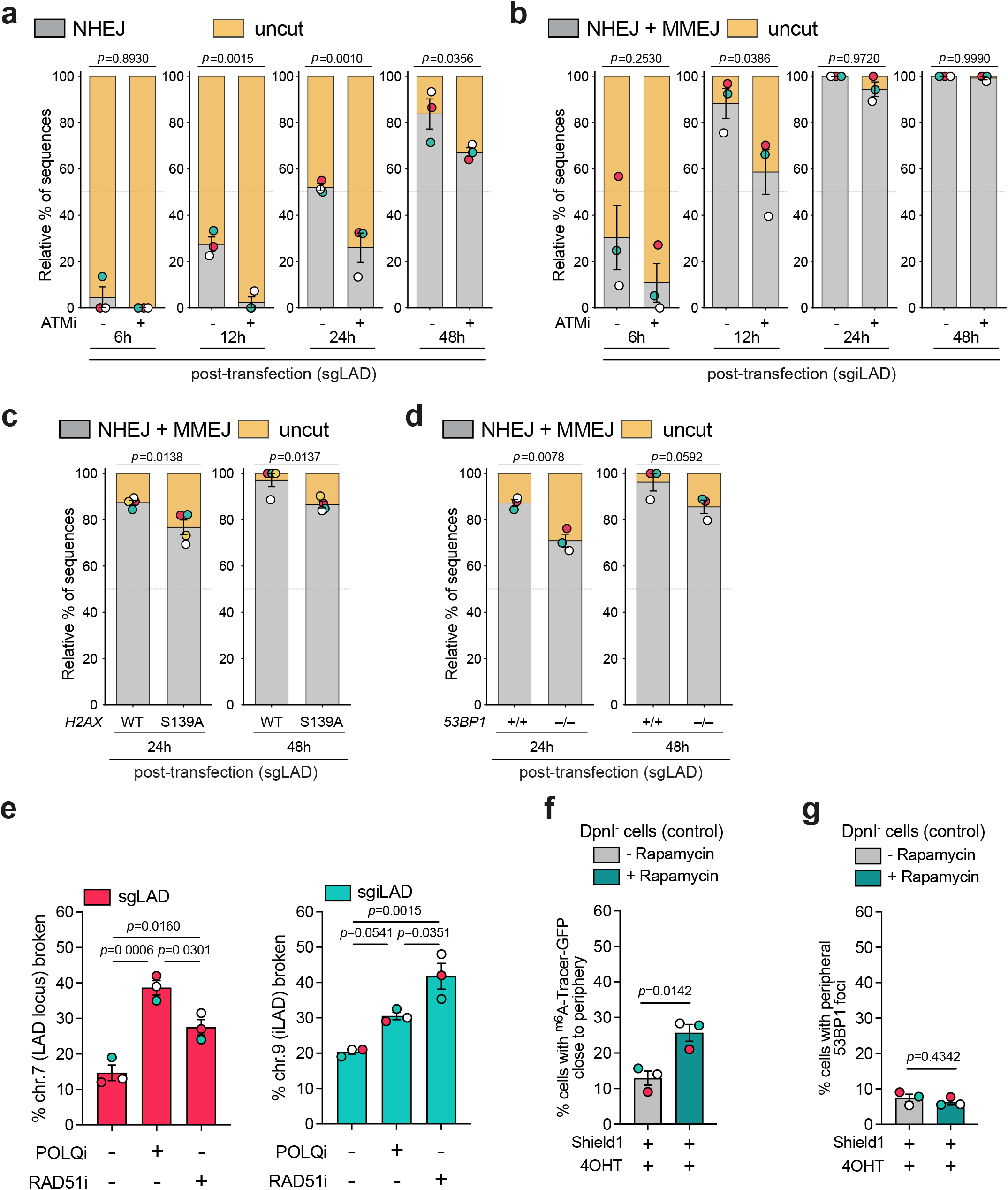
a) Relative percentage of significant indels quantified by TIDE, in RPE-1 cells expressing Cas9 6, 12, 24 and 48 hours post transfection with sgLAD and treated with ATM inhibitor (ATMi; 10µM) or mock (DMSO). The data represents the percentage of significant (*p* < 0.001) sequences with a +1 insertion indicative of NHEJ, and no insertions or deletions (‘0”) indicative of an uncut sequence. Data are presented as mean ± SEM from 3 biological replicates. Statistical comparisons were performed using one-way ANOVA with planned pairwise comparisons of DMSO and ATMi within each timepoint, with Šídák correction for multiple comparisons. b) Relative percentage of significant indels quantified by TIDE, in RPE-1 cells expressing Cas9 6, 12, 24 and 48 hours post transfection with sgiLAD and treated with ATM inhibitor (ATMi; 10µM) or mock (DMSO). The data represents the percentage of significant (*p* < 0.001) sequences with indels derived from NHEJ and MMEJ, and no insertions or deletions (‘0”) indicative of an uncut sequence. Data are presented as mean ± SEM from 3 biological replicates. Statistical comparisons were performed using one-way ANOVA with planned pairwise comparisons of DMSO and ATMi within each timepoint, with Šídák correction for multiple comparisons. c) Relative percentage of significant indels quantified by TIDE, in WT (*H2AX*^WT^) and *H2AX*^S139^ mutant RPE-1 cells 24 and 48 hours post transfection with sgLAD. The data represents the percentage of significant (*p* < 0.001) sequences with indels derived from NHEJ and MMEJ, and no insertions or deletions (‘0”) indicative of an uncut sequence. Data are presented as mean ± SEM from 4 biological replicates. Statistical comparisons were performed using one-way ANOVA with planned pairwise comparisons of *H2AX*^WT^ and *H2AX*^S139^ mutant cells within each timepoint, with Šídák correction for multiple comparisons. d) Relative percentage of significant indels quantified by TIDE, in WT (*53BP1*^+/+^) and 53BP1-deficient (*53BP1*^-/-^) RPE-1 cells 24 and 48 hours post transfection with sgLAD. The data represents the percentage of significant (*p* < 0.001) sequences with indels derived from NHEJ and MMEJ, and no insertions or deletions (‘0”) indicative of an uncut sequence. Data are presented as mean ± SEM from 3 biological replicates. Statistical comparisons were performed using one-way ANOVA with planned pairwise comparisons of *53BP1*^+/+^ and *53BP1*^-/-^ cells within each timepoint, with Šídák correction for multiple comparisons. e) *Left*, quantification of broken chromosome 7 (chr.7) per 100 metaphases in RPE-1 expressing Cas9 transfected with either non-targeting control (sgCtrl) or sgLAD. Cells were treated with control conditions (DMSO) or the inhibitors Pol theta (PolQi, 10µM) or RAD51i (20µM). Data are presented as mean ± SEM from 3 biological replicates. Statistical comparisons were performed using one-way ANOVA with Bonferroni correction for multiple comparisons. *Right*, quantification of broken chromosome 9 (chr.9) per 100 metaphases in RPE-1 expressing Cas9 transfected with either non-targeting control (sgCtrl) or sgiLAD. Cells were treated with control conditions (DMSO) or the inhibitors Pol theta (PolQi, 10µM) or RAD51i (20µM). Data are presented as mean ± SEM from 3 biological replicates. Statistical comparisons were performed using one-way ANOVA with Bonferroni correction for multiple comparisons. f) Quantification of DpnI^-^ cells treated with Shield1 and 4OHT with m^6^A-Tracer-GFP localised close to the nuclear periphery in the absence or presence of Rapamycin, as indicated. The graphs represent the average of 3 biological replicates. Statistical comparisons were performed using unpaired *t*-test. g) Percentage of cells showing 53BP1 foci at the nuclear periphery in DpnI^-^ cells expressing FRB fused to PRR14 and GFP-nanobody fused to FKBP, treated with Shield1 and 4OHT and in the absence or presence of Rapamycin, as indicated. Data is represented as a percentage of total PRR14 positive nuclei and is the mean of 3 independent experiments. Statistical comparisons were performed using unpaired *t*-test.

**Extended Figure 9.**
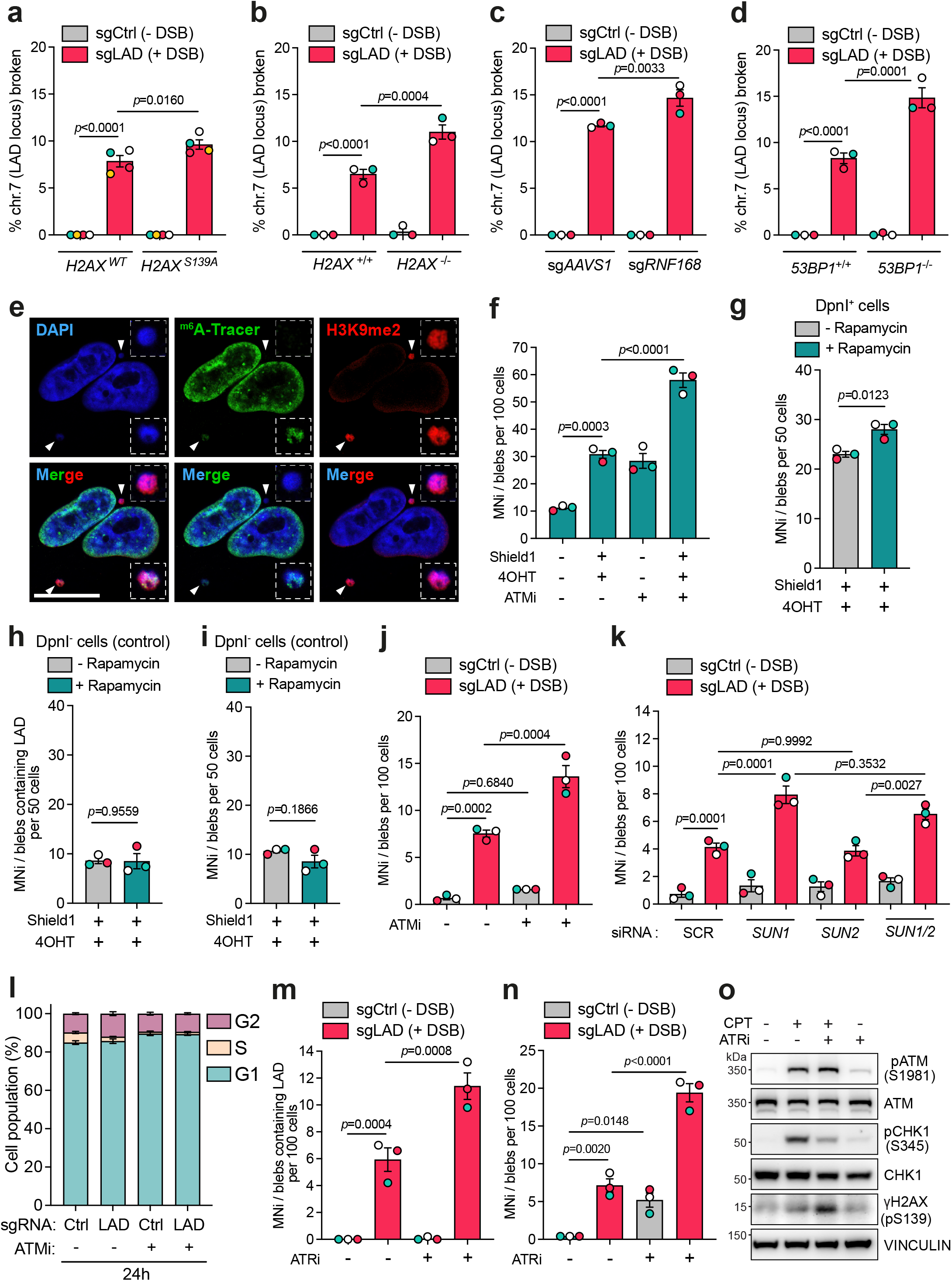
a) Quantification of broken chromosome 7 (chr.7) per 100 metaphases containing LAD in WT (*H2AX*^WT^) and *H2AX*^S139^ mutant RPE-1 cells transfected with Cas9 and non-targeting control guide (sgCtrl) or sgLAD for 7 h. Data are presented as mean ± SEM from 3 biological replicates. Statistical comparisons were performed using two-way ANOVA followed by Šídák’s multiple comparisons test. b) Quantification of broken chr.7 per 100 metaphases containing LAD in WT (*H2AX*^+/+^) and H2AX-deficient (*H2AX*^-/-^) RPE-1 cells transfected with Cas9 and non-targeting control guide (sgCtrl) or sgLAD for 7 h. Data are presented as mean ± SEM from 3 biological replicates. Statistical comparisons were performed using two-way ANOVA followed by Šídák’s multiple comparisons test. c) Quantification of broken chr.7 per 100 metaphases containing LAD in mock-depleted (sg*AAVS1*) or RNF168-depleted (sg*RNF168*) RPE-1 cells expressing Cas9 transfected either control guide (sgCtrl) or sgLAD for 7 h. Data are presented as mean ± SEM from 3 biological replicates. Statistical comparisons were performed using two-way ANOVA followed by Šídák’s multiple comparisons test. d) Quantification of broken chr.7 per 100 metaphases containing LAD in WT (*53BP1*^+/+^) and 53BP1-deficient (*53BP1*^-/-^) RPE-1 cells transfected with Cas9 and non-targeting control guide (sgCtrl) or sgLAD for 7 h. Data are presented as mean ± SEM from 3 biological replicates. Statistical comparisons were performed using two-way ANOVA followed by Šídák’s multiple comparisons test. e) Representative confocal microscopy images of micronuclei containing LADs (m^6^A-Tracer; green) or H3K9me2 (red) in DpnI expressing cells treated with Shield1 and 4OHT. DNA is stained with DAPI (blue). Scale bar 20µm. f) Quantification of MNi/blebs (based on DAPI) in DpnI expressing cells in the absence (-Shield1 -4OHT) or presence (+Shield1 +4OHT) of DNA damage, treated with either DMSO or ATM inhibitor (ATMi, 10µM). Data are presented as mean ± SEM from 3 biological replicates. Statistical comparisons were performed using two-way ANOVA followed by Šídák’s multiple comparisons test. g) Quantification of cells with MNi/blebs (based on DAPI) in DpnI expressing cells transfected with PRR14-FRB/FKBP-GFP nanobodies following DSB induction with Shield1 and 4OHT and in the absence or presence of Rapamycin, as indicated. Data are presented as mean ± SEM from 3 biological replicates. Statistical comparisons were performed using unpaired *t*-test. h) Quantification of cells with MNi/blebs containing LADs (m^6^A-Tracer) in DpnI^-^ cells transfected with PRR14-FRB/FKBP-GFP nanobodies following DSB induction with Shield1 and 4OHT and in the absence or presence of Rapamycin, as indicated. Data are presented as mean ± SEM from 3 biological replicates. Statistical comparisons were performed using unpaired *t*-test. i) Quantification of cells with MNi/blebs (based on DAPI) in DpnI^-^ cells transfected with PRR14-FRB/FKBP-GFP nanobodies following DSB induction with Shield1 and 4OHT and in the absence or presence of Rapamycin, as indicated. Data are presented as mean ± SEM from 3 biological replicates. Statistical comparisons were performed using unpaired *t*-test. j) Quantification of MNi/blebs (based on DAPI) in RPE-1 cells expressing Cas9 and transfected with non-targeted control (sgCtrl) and sgLAD in control conditions (DMSO) or treated with ATMi (10µM). Data are presented as mean ± SEM from 3 biological replicates. Statistical comparisons were performed using two-way ANOVA followed by Šídák’s multiple comparisons test. k) Quantification of MNi/blebs (based on DAPI) in RPE-1 cells expressing Cas9 in control conditions (SCR) or depleted for *SUN1* (si*SUN1*), *SUN2* (si*SUN2*) or both (si*SUN1*/*SUN2*) and then transfected with non-targeted control (sgCtrl) or sgLAD. Data are presented as mean ± SEM from 3 biological replicates. Statistical comparisons were performed using two-way ANOVA followed by Šídák’s multiple comparisons test. l) RPE-1 cells transfected with non-targeting control (sgCtrl) or sgLAD, treated with either DMSO or ATMi (10µM) for 24h were stained for incorporated BrdU against total DNA content using propidium iodide. Data are presented as the mean percentage of cells in each phase of the cell cycle ± SEM from 3 biological replicates. m) Quantification of MNi/blebs containing the LAD locus in RPE-1 cells expressing Cas9 and transfected with non-targeted control (sgCtrl) and sgLAD in control conditions (DMSO) or treated with ATRi (5µM). Data are presented as mean ± SEM from 3 biological replicates. Statistical comparisons were performed using two-way ANOVA followed by Šídák’s multiple comparisons test. n) Quantification of MNi/blebs (based on DAPI) in RPE-1 cells expressing Cas9 and transfected with non-targeted control (sgCtrl) and sgLAD in control conditions (DMSO) or treated with ATRi (5µM). Data are presented as mean ± SEM from 3 biological replicates. Statistical comparisons were performed using two-way ANOVA followed by Šídák’s multiple comparisons test. o) Immunoblot of phosphorylation of ATM (pATM), CHK1 (pCHK1) and phospho-S139 (pS139) of H2AX (γH2AX) in RPE-1 cells expressing Cas9, in the absence and presence of 1 h CPT (5µM). Cells were treated with the ATR inhibitor (ATRi; 5µM) or mock (DMSO) for 24 h. VINCULIN was used as loading control.

**Extended Data Figure 10.**
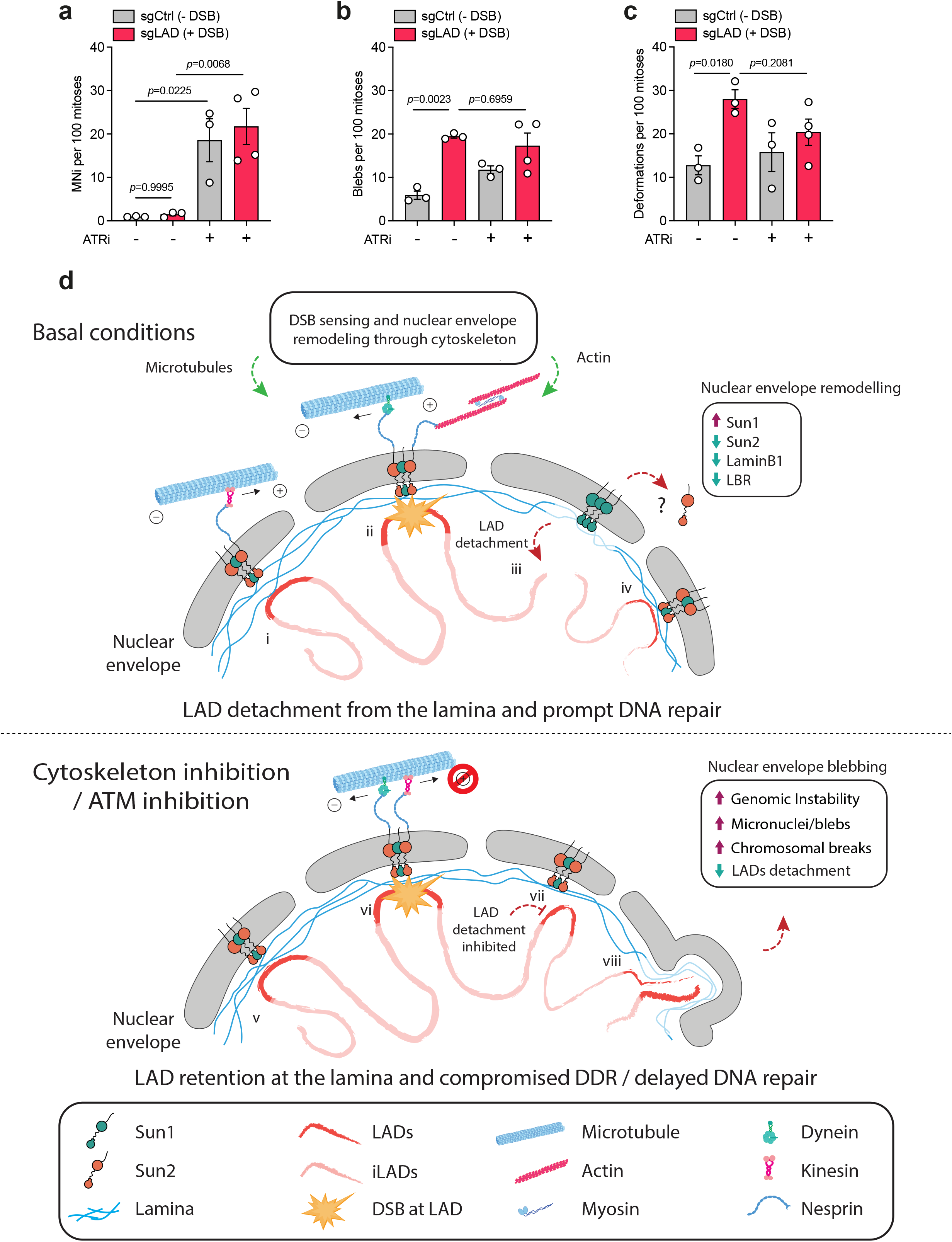
a-c) Quantification of micronuclei (MNi) (a), blebs (b) and deformations (c) in one or both daughter cells per 100 mitoses in RPE-1 cells expressing GFP-H2B, NLS-RFP, and SNAP-MDC1 transfected with Cas9 and non-targeting control (sgCtrl) or sgLAD in control conditions (DMSO) or treated with ATRi (5µM). Data are presented as mean ± SEM from 3 or 4 biological replicates. Statistical comparisons were performed using two-way ANOVA followed by Šídák’s multiple comparisons test. d) Model of LAD detachment in response to DNA damage and its functional implications. LADs are genomic regions that interact with the nuclear lamina at the periphery of the nucleus (i). Upon DNA damage at LADs (ii), structural remodeling of the nuclear lamina is triggered (iii), along with ATM-dependent reorganization of the LINC complex (iii). This reorganization mechanically couples the damaged chromatin to the microtubule and actin cytoskeletal networks, which exert active mechanical forces to drive the relocalization of damaged LADs from the nuclear periphery to the nuclear interior, enabling prompt DNA repair (iii). Once repair is complete, LADs return to the nuclear periphery (iv). Inhibition of ATM or microtubules polymerisation (v, vi) disrupts this dynamic detachment and relocalization process (vii), leading to impaired DNA damage response, compromised nuclear integrity, and increased genomic instability (vii).

### Supplementary video 1

Representative video from live-cell imaging of RPE-1 cells expressing GFP-H2B, NLS-RFP, and SNAP-MDC1 transfected with Cas9 and sgLAD and treated with ATM inhibitor (10µM), highlighting MN formation (green arrowheads). White arrowheads indicate the mother cell of interest. Video associated with Fig 6A. Scale bar 10 µm.

### Supplementary video 2

Representative video from live-cell imaging of RPE-1 cells expressing GFP-H2B, NLS-RFP, and SNAP-MDC1 transfected with Cas9 and sgLAD and treated with ATM inhibitor (10µM), highlighting MNi formation (green arrowheads). White arrowheads indicate the mother cell of interest. Scale bar 10 µm.

### Supplementary video 3

Representative video from live-cell imaging of RPE-1 cells expressing GFP-H2B, NLS-RFP, and SNAP-MDC1 transfected with Cas9 and sgLAD and treated with ATM inhibitor (10µM), highlighting blebbing (purple arrowheads) and nuclear deformations (orange arrowheads). White arrowheads indicate the mother cell of interest. Video associated with Fig 6A. Scale bar 10 µm.

### Supplementary video 4

Representative video from live-cell imaging of RPE-1 cells expressing GFP-H2B, NLS-RFP, and SNAP-MDC1 transfected with Cas9 and sgLAD, highlighting blebbing (purple arrowheads) and nuclear deformations (orange arrowheads). White arrowheads indicate the mother cell of interest. Video associated with Fig 6A. Scale bar 10 µm.

### Supplementary video 5

Representative video from live-cell imaging of RPE-1 cells expressing GFP-H2B, NLS-RFP, and SNAP-MDC1 transfected with Cas9 and sgLAD and treated with ATM inhibitor (10µM), highlighting blebbing (purple arrowheads). White arrowheads indicate the mother cell of interest. Video associated with Fig 6A. Scale bar 10 µm.

### Supplementary video 6

Representative video from live-cell imaging of RPE-1 cells expressing GFP-H2B, NLS-RFP, and SNAP-MDC1 transfected with Cas9 and sgLAD and treated with ATM inhibitor (10µM), highlighting nuclear deformations (orange arrowheads). White arrowheads indicate the mother cell of interest. Video associated with Fig 6A. Scale bar 10 µm.

### Supplementary video 7

Representative video from live-cell imaging of RPE-1 cells expressing GFP-H2B, NLS-RFP, and SNAP-MDC1 transfected with Cas9 and sgLAD and treated with ATM inhibitor (10µM), highlighting various instances of post-mitotic MNi formation (green arrowheads), blebbing (purple arrowheads) and nuclear deformations (orange arrowheads). White arrowheads indicate the mother cell of interest. Scale bars 10 µm.

## Supplementary information

### Materials and Methods

#### Cell lines and culture conditions

Shield1-inducible Dam-Lamin B1 HT1080 clone11 human cells^1^ were grown in low glucose Dulbecco’s Modified Eagle’s Medium (DMEM; Gibco) supplemented with 10% fetal bovine serum (FBS; Gibco), 1% penicillin-streptomycin (Sigma-Aldrich), 1% non-essential amino acids (NEAA; Sigma), G418 (25 µg/ml, Sigma Aldrich, A1720-5G) and puromycin (0.5 µg/ml, Sigma Aldrich, P9620). hTERT-immortalized human RPE-1 cells stably expressing Cas9^2^, RPE-1 cells (ATCC), RPE-1 *H2AX*^S139A^ cells^3^, RPE-1 *53BP1*^-/-^ cells^4^, RPE-1 FRT-derived cells^5^, RPE-1 FRT *H2AX*^-/-^ cells^5^ and hTERT-immortalized human RPE-1 cells with stable expression of GFP-H2B RFP-NLS SNAP-MDC1^6^ were grown in DMEM/F12 medium (Sigma Aldrich) supplemented with 10% FBS and 1% penicillin-streptomycin. Cell lines were cultured at 37°C in a 5% CO_2_ atmosphere. All cell lines regularly tested negative for mycoplasma contamination. Shield1-inducible Dam-Lamin B1 HT1080 clone11 and hTERT RPE-1 Cas9 cells were generous gifts from B. van Steensel and D. Durocher, respectively. RPE-1 *H2AX*^S139A^ knock in, RPE-1 FRT, RPE-1 FRT *H2AX*^-/-^ and RPE-1 *53BP1*^-/-^ cells were a generous gift from S. P. Jackson’s lab. All cell lines can be found in Supplementary Table 1.

#### Plasmid generation

PSA05-AID-ER*-DPN1 and PSA11-GFP-m^6^A-Tracer-APEX2-NES plasmids were assembled using golden gate cloning^7^. To generate PSA13-AID-ER*-DPN1, cassette AID-ER*-DPN1/TIR1 was PCR amplified (Primer 1 and Primer 2; IDT) from PSA05-AID-ER*-DPN1 and ligated into PLKU06-Hygro digested with ApaI and BamHI. Plasmids for PRR14 tethering were adapted from a previous study^8^. The pMito-mCherry-FRB_IRES_FKBP(III)-GBPen plasmid was a kind gift from Professor Chris Chan (Addgene #128268). pMito-mCherry was removed using restriction sites AgeI and BsrGI. mCherry was amplified using primers (Primer 3, and Primer 4) and fused by PCR with a custom insert of codon-optimised PRR14 gBlock (IDT) using primers (Primer 5 and Primer 6). NEBuilder HiFi DNA assembly cloning kit (New England Biolabs, E5520S) was used to assemble PRR14-mCherry-FRB_IRES_FKBP(III)-GBPen. All oligonucleotide sequences can be found in Supplementary Table 2.

#### Cell lines generation

To generate a DpnI⁻ cell line, Shield1-inducible Dam-Lamin B1 clonal line clone11 HT1080 cells were transfected with PSA11 carrying NES-m^6^A-GFP-APEX2 containing a puromycin resistance gene using Lipofectamine 2000 (Invitrogen, 11668-019), according to manufacturer’s instructions. Cells were treated with 10 µg/ml Puromycin (Sigma Aldrich, P9620-10ML) for 5 days. Single colonies were picked and expanded, and cells were treated with 0.5 µM Shield1 (Takara, 632189; Supplementary Table 3) for 18 h. Clones were seeded for GFP immunofluorescence, as described below. The clone with best GFP ring induction was identified and used for subsequent experiments. To generate a DpnI+ cell line, DpnI⁻ cell line was transduced with the lentiviral vector PSA13-AID-ER*-DPN1 containing a BFP expression cassette. The lentivirus was prepared by transfection of HEK293T cells with third generation lentiviral plasmids, packaging plasmid 17.5 µg psPAX2 (packaging plasmid), 7.5 µg pMD2.G (envelope plasmid) and 25 µg of PSA13-AID-ER*-DPN1 using Lipofectamine 2000, according to the manufacturer’s instructions. Viral particles were collected after 48 h, filtered (0.45 µm), and then used to infect 1x10^6^ Dpn1^-^ cells per 1 ml of virus. 48 hours after seeding, cells were FACS sorted for BFP expression. Single clones were expanded and tested for induction of 53BP1 and γH2AX foci formation at the nuclear periphery following 0.5 µM Shield1 (18 h) and 300 nM (Z)-4-Hydroxytamoxifen (4OHT; Sigma Aldrich, H7904; Supplementary Table 3) (2 h) treatment. The clone with the highest induction of damage was selected for further use. In pool knock-outs for *MDC1*, *RNF168*, *LMNB1* and *LBR* genes were generated through tracrRNA:crRNA transfection (see below) and used to seed experiments 72 h post-transfection.

#### Cas9 RNP transfection

For live cell imaging experiments, 2x10^4^ RPE-1 GFP-H2B RFP-NLS SNAP-MDC1 cells were seeded onto a µ-Dish 35 mm, high (ibidi, 81156). 24 h later, cells were transfected with 250 ng TrueGuide sgLAD or sgCtrl (Supplementary Table 2) and 750 ng TrueCut™ Cas9 Protein v2 (Thermo Fisher Scientific, A36498) using Lipofectamine™ CRISPRMAX™ Cas9 Transfection Reagent (Thermo Fisher Scientific, CMAX00008) according to manufacturer’s protocol. At the time of transfection, cells were also treated with either 10 µM ATMi KU-55933 (Sigma-Aldrich), 5 µM ATRi VE-821 (Sigma-Aldrich) or DMSO vehicle.

#### TracrRNA:crRNA transfection

Alt-R™ CRISPR-Cas9 tracrRNA (cat. number 1073191) and crRNAs (Alt-R™ Cas9 Negative Control crRNA #1 (cat. number 1079138), and CATTGACGATCATCCCACTG (Chr7 sgLAD; gRNA #1, see Supplementary Table 2 for details of all gRNAs)^9^, AGGTCAGCCCACCCCCAGTT (Chr9 sgiLAD; gRNA #2,)^10^ TGAATCCTTGAGGTGTAACG (sgMDC1 gRNA #3), GAGTCCACGACGATACCCGG (sgRNF168; gRNA #4), CGAGCTGGGCAAGTGCAAGG (sgLMNB1; gRNA #5) and CTTCACAGTGTAAAGCTGGG (sgLBR, gRNA #6) were purchased from Integrated DNA Technologies (IDT) and transfected into Cas9 expressing RPE-1 cells using Lipofectamine RNAiMAX (Invitrogen), following manufacturer’s instructions. Briefly, tracrRNA and crRNA were mixed in a 1:1 molar ratio and annealed at 95°C for 5 min, followed by gradual cooling to 20°C in a thermocycler. The resulting tracrRNA:crRNA duplex was diluted in Opti-MEM Medium (Gibco) and mixed with RNAiMAX (Invitrogen), also diluted in Opti-MEM. After a 15 min incubation at RT, the sgRNA-lipid transfection complexes were added dropwise to the culture plate. For RPE-1 cell lines that do not express Cas9, transfections with gRNA-duplex and Cas9 Protein v2 (Thermo Fisher Scientific) using Lipofectamine™ CRISPRMAX™ Cas9 Transfection Reagent (Thermo Fisher Scientific) were performed according to manufacturer’s protocol. At the time of transfection, cells were also treated with either 10 µM ATMi KU-55933 (Sigma-Aldrich), 5 µM ATRi VE-821 (Sigma-Aldrich), 100 µM Arp2/3i CK666 (Sigma-Aldrich), 5 µM UBA1i TAK-234 (MedChemExpress), 100 nM microtubule polymerization inhibitor Vinblastine (Sigma-Aldrich) or DMSO vehicle. All tracrRNA and crRNAs sequences can be found in Supplementary Table 2.

#### siRNA transfection

Dharmacon ON-TARGETplus Non-targeting pool (D-001810-10-20), SUN1 (L-025277-00-0005) and SUN2 (L-009959-01-0005) siRNA SMARTpools were transfected for 48 h using Lipofectamine RNAiMAX, according to manufacturer’s instructions. Briefly, siRNAs were diluted in Opti-MEM Medium and mixed with RNAiMAX, also diluted in Opti-MEM. After a 15 min incubation at RT, the siRNA-lipid transfection complexes were added dropwise to the culture plate.

#### Immunofluorescence (IF)

Cells were grown on glass coverslips and treated as indicated. Cells were pre-extracted (2 min on ice in 0.2% Triton X-100/PBS), and then fixed (10 min in 4% paraformaldehyde/PBS), permeabilized (5 min in 0.2% Triton X-100/PBS), blocked 1 h in 5% bovine serum albumin (BSA/PBS; Sigma Aldrich, A7030-100G), and incubated with the indicated primary antibodies (1 h at RT or overnight at 4°C in 1% BSA/PBS), followed by washing (3 × 5 min in 0.1% Tween20-PBS (PBS-T) and incubation with the corresponding AlexaFluor-conjugated secondary antibody (1 h at RT in 1% BSA/PBS) and washed again as described above. Finally, cells were counterstained with DAPI in antifade mounting medium for fluorescence (Vectashield, Vector Labs, H-1200). Primary antibodies: anti-γH2AX (Millipore, 05-636) 1:1000, anti-γH2AX (abcam, ab22551) 1:1000, anti-53BP1 (Novus Biologicals, NB100-904) 1:2500, anti-53BP1 (Millipore, MAB3802) 1:500, anti-Emerin (abcam, ab156871) 1:200, anti-GFP (abcam, ab290) 1:1000, anti-Lamin A (abcam, ab26300) 1:2000, anti-Lamin B1 (abcam, ab16048) 1:2000, anti-Lamin B1 (Sangon Biotech, D194916-0100) 1:200, anti-LAP2 (ThermoFisher Scientific, MA1-25323) 1:200, anti-LBR (abcam, ab32535) 1:200, anti-Nup153 (abcam, ab24700) 1:500, anti-pATM (abcam, 81292) 1:500, anti-MDC1 (abcam, ab11171) 1:400, anti-SUN1 (Novus Biologicals, NBP1-87396) 1:500, anti-SUN2 (abcam, ab124916) 1:500. Secondary antibodies: Alexa Fluor 488-donkey anti-rabbit IgG (H+L), Alexa Fluor 546-donkey anti-rabbit IgG (H+L), Alexa Fluor 546-goat anti-mouse IgG (H+L), Alexa Fluor 568-donkey anti-rabbit IgG (H+L), Alexa Fluor 568-donkey anti-mouse IgG (H+L), Alexa Fluor 647-donkey anti-goat IgG (H+L), Alexa Fluor 647-donkey anti-mouse IgG (H+L), Alexa Fluor 647-goat anti-rabbit IgG (H+L) (ThermoFisher Scientific A21206, A10040, A11003, A10042, A10037, A21447, A31571, A21244, respectively, at 1:500 dilution). Antibodies used in this study are summarized in Supplementary Table 4.

#### 53BP1 foci induced by Cas9

Cells with two 53BP1 foci were scored at various time points post-transfection. Values are shown as the percentage of cells with two 53BP1 foci. For each data point in each independent experiment, at least 200 cells were quantified. Scoring was performed manually under double-blind conditions.

#### γH2AX and pATM colocalisation

All γH2AX foci, and all pATM foci colocalising with a γH2AX focus were quantified in at least 100 cells per condition. The colocalisation percentage was calculated as the total number of pATM foci colocalised with a γH2AX focus relative to all γH2AX foci in conditions containing at least one γH2AX focus. Scoring was performed manually under double-blind conditions.

#### Metaphase spreads

Metaphase spreads were performed as previously described^11^. Cells were incubated with 0.2 µg/ml demecolcine (Sigma Aldrich, D1925) for 13 h and then harvested. Cells were collected using standard cytogenetic techniques, subject to hypotonic shock for 30 min at 37°C in 0.03 M sodium citrate and fixed in 3:1 methanol:acetic acid solution. Fixed cells were dropped onto acetic acid-humidified slides before dehydration and FISH.

#### Fluorescence *in situ* hybridization (FISH)

Whole chromosome FISH was performed according to the manufacturer’s protocol (MetaSystems probes, Whole Chromosome Paint, D-0307-050-FI & D-0309-050-OR). Briefly, sample and probe were simultaneously denatured (2 min at 75°C) and incubated for hybridization (overnight at 37°C). Post-hybridization washes included 0.4X SSC (2 min at 72°C), followed by 2X SSC, 0.05% Tween-20 (30 seconds at RT). Slides were then counterstained with DAPI in antifade mounting medium for fluorescence. When indicated, IF of anti-lamin B1 (abcam, ab16048; Supplementary Table 4) or anti-Lamin A (abcam, ab26300; Supplementary Table 4) was performed followed by FISH according to manufacturer’s protocol (Empire Genomics probes, RP11-460J13 BAC (LAD) & RP11-118N16 BAC (iLAD)).

#### Intensity measurements of proteins at the nuclear envelope

The intensity of nuclear envelope proteins under different experimental conditions was quantified using a custom macro plug-in designed for FIJI (ImageJ). Nuclei were first segmented based on a DAPI to generate masks for individual cells. To isolate the nuclear periphery, the masks were eroded (4 pixels total) to exclude the periphery of the nucleus. The intensity at the nuclear periphery was then calculated by subtracting the mean intensity of the internal (eroded) nuclear region from the total nuclear intensity for each cell. Intensities were quantified from at least 50 cells per replicate per condition.

#### LADs position analysis relative to NUP

Image analysis was performed using Fiji (ImageJ). Multi-channel images were split, and the DAPI channel was pre-processed using a Gaussian blur filter (radius = 1.50). Brightness and contrast were adjusted to fully saturate the signal, and binary nuclear masks were generated using Otsu thresholding with a dark background. Nuclear objects were segmented using the Analyze Particles tool (size filter: 800– pixel units, including holes). Masks were subsequently inverted, and a border mask was created with inner and outer widths set to 1 pixel.

Nuclear pore foci distributions were analyzed using the GDSC Plugins for ImageJ (version 2.0-SNAPSHOT, GDSC-Core version 2.3-SNAPSHOT; developed by Alex Herbert^12^. Within the FindFoci GUI, foci were detected relative to the generated ring mask on the NUP153 channel, with output coordinates saved to memory. A secondary nuclear mask was generated via a repeated particle analysis and LUT inversion workflow, and spot radial intensity profiles were quantified using the FindFoci Spot Radial Intensity module combined with the stored memory results.

#### Chromosomal breaks

Chromosomal breaks in metaphase spreads were quantified as the percentage of broken chromosome 7 (containing the LAD-targeting site). For each condition in each independent experiment, at least 50 metaphases were quantified. Scoring was performed manually under double-blind conditions.

#### Chromosomal translocations

Translocation frequencies in metaphase spreads were calculated as translocations per 100 metaphases in chromosomes 7 and 9, and plotted together. For each condition in each independent experiment, at least 150 metaphases were quantified. Scoring was performed manually under double-blind conditions.

#### Micronuclei/blebs

Micronuclei/blebs were analysed in HT1080 DpnI⁺ cells and RPE-1 cells previously seeded onto coverslips. For HT1080 DpnI⁺ cells, 10 µM ATM inhibitor (ATMi; KU-55933, Sigma Aldrich), or DMSO, was added for the last 30 min of the -Shield1/-4OHT or +Shield1 (0.5 µM)/+4OHT (300 nM) treatments. After treatment, the media was changed and Auxin (500 µg/ml) was added with 10 µM ATMi or DMSO, and Shield1 (0.5 µM). A summary of all chemical treatments can be found in Supplementary Table 3. 24 h post-treatment, cells were fixed and subject to IF using anti-GFP (abcam, ab290) and anti-H3K9me2 (abcam, ab1220) antibodies, as described above (Supplementary Table 4). For RPE-1 cells, 10 µM ATMi, 5 µM ATRi or DMSO were added at the moment of transfection. 24 h post-transfection, cells were fixed and subject to immuno-FISH as described above. For each condition, in each independent experiment, at least 500 cells were quantified. Scoring was performed manually under double-blind conditions.

#### PRR14 LAD tethering induction

Cells were seeded and transfected using Lipofectamine 2000 following manufacturer’s instructions. 24 h post-transfection cells were re-seeded onto glass coverslips. 6 h post reseeding, cells were treated with 0.5 µM Shield1 for 18 h. Following this, cells were treated with 200nM Ramapycin (553210, Sigma Aldrich; Supplementary Table 3) or DMSO control for 30 min, before addition of 300 nM 4OHT for 2 h. Cells were collected for 0, 4 and 8 h timepoints. For the 4 and 8 h samples, the cells were washed three times in PBS at 0h, and supplemented with media containing 500 µg/ml Auxin (I5148, Sigma Aldrich) +/- Rapamycin or DMSO control. For IF, cells were pre-extracted (30s on ice in 0.5% Triton X-100/PBS) and then fixed (10 min in 4% paraformaldehyde/PBS, on ice). An additional fixation with 4% paraformaldehyde/PBS at RT was then performed for 10 min. Coverslips were washed twice in PBS and then stored in PBS for future immunostaining alongside later timepoints. Cells were blocked for 1 h in 5% BSA/PBS and immunostaining conducted using primary antibodies against mCherry, 53BP1 and GFP (1 h at RT in 1% BSA/PBS). Antibody details are provided in Supplementary Table 4. Coverslips were then washed three times, for 5 min each, in PBS-T and incubated in the appropriate secondary antibody (1 h at RT in 1% BSA/PBS; Supplementary Table 4). Coverslips were washed again for three 5 min washed in PBS-T and counterstained with DAPI in antifade mounting medium for fluorescence (Vectashield, Vector Labs (H-1200)).

#### Microscopy

Confocal images were taken using a LSM880 AxioObserver confocal microscope (Zeiss) with a C Plan-Apochromat 63x.1.4 Oil DIC UV-VIS_IR M27 lens. Microscopy images for Cas9 expressing cells were taken using Apotome Axio Observer Z1/7 microscope (Zeiss) using a 100x/1.40 Oil DIC M27 lens. For immunoFISH, to analyze if the LAD or iLAD foci were attached to the lamina, Z-stacks were taken at 0.2µm intervals. These images were deconvolved using Huygens Professional 25.4 (Scientific Imaging B.V). Images were analyzed using Fiji imageJ.

Prior to live-cell microscopy, 250 nM JF646-SNAP-tag ligand (Janelia Materials) was added to the Cas9 RNP-transfected RPE-1 GFP-H2B RFP-NLS SNAP-MDC1. Live cell imaging was then performed with the Visitron Spinning Disc Confocal System VisiScope 5-Elements in an environmental enclosure to maintain cell culture conditions (37 °C and humidified 5% CO_2_). Images were collected for 24 h with a 20x/0.75 Air objective, at intervals of 20 min. Live cell imaging was analyzed with Fiji imageJ. Approximately 100 random mitoses were considered for each replicate and scored based on whether one or both daughter cells present nuclear deformations, blebs, or MNi formation.

#### Cell cycle analysis

For RPE-1 cells, cells were incubated with 10 µM BrdU (Sigma, B5002) for the last 15 min of the indicated treatments. Cells were then washed with PBS, trypsinised and fixed with 70% ethanol. DNA was denatured with 2 N HCl/0.5% Triton X-100 for 30 min, and then neutralized in 0.1 M sodium tetraborate for 2 min. Cells were incubated with anti-BrdU-FITC (Becton Dickinson, 347583) for 1 h at RT. Finally, before flow cytometry, cells were incubated with 20 µg/ml PI and 100 µg/ml RNAse A for 30 min at RT. Data was collected in a BC Cytoflex flow cytometer and analyzed in CytExpert Software (CytoFLEX Platform.v. 2.4.0.28).

For HT1080 DpnI cells, cells were seeded in a 96 well plate at 0.4 × 10^4^ per well (Revvity, Cat# 6055300) and 24 h later cells were treated with Shield1 for 18h. Then cells were incubated with ATMi or DMSO 30min prior incubation of 2 h with 4OHT. A 10 µM EdU pulse was added (ThermoFisher, Cat# C10640) for the last 30 min. Cells were then washed twice with 1× PBS and pre-extracted (1XPBS, 0.5% Triton X-100) for 5 min on ice, and then fixed with 4% formaldehyde for 10 min. Cells were permeabilized using 1XPBS, 0.5% Triton for 10 min and blocked for 1h in BSA. EdU click reaction was carried out after the blocking step using Click-iT EdU Alexa Fluor 647 Imaging Kit (ThermoFisher, Cat# C10640) according to the manufacturer’s instructions. DNA was counterstained with DAPI. Immunofluorescence images were acquired using an Olympus IX81 microscope equipped with scanR screening system using ×40 objective at a single autofocus-directed z-position under non saturating settings. Olympus scanR image analysis software was used to analyze and quantify the fluorescence intensity of individual cells (thousands per sample).

#### PCR and Sanger Sequencing for Tracking of Indels by Decomposition (TIDE)

Briefly, genomic DNA from cells transfected with gRNA was extracted with DNeasy Blood and Tissue Kit (QIAGEN, 69504), following manufacturer’s instructions. PCR of the target region was carried out using 100 ng of gDNA and PrimeSTAR Max DNA Polymerase (Takara, R045A) according to manufacturer’s instructions (see Supplementary Table 2 for primers used). PCR conditions were: 1 min at 98°C once, then 10 seconds at 98°C, 10 seconds at 60°C, and 10 seconds at 72°C for 34 cycles, then 2 min at 72°C once. PCR products were inspected for quality and yield by 1.2% agarose gel electrophoresis in 1X TBE, and were purified by 0.9X ratio of AMPure XP Beads (Beckman Coulter, A63881), according to the manufacturer’s protocol. Samples were sequenced by Eurofins Genomics TubeSeq Supreme, (see Supplementary Table 2 for the primers used).

#### TIDE

Analysis of sequenced products to identify insertions and deletions (indels) produced after repair of Cas9-induced DSBs were performed as previously described^13^, with TIDE version 5.0.0. Briefly, raw ab1 files were inspected visually for quality control. Samples transfected with non-targeting gRNA (Alt-R™ Cas9 Negative Control crRNA #1; cat. number 1079138) for the appropriate cell line and condition were used as a control unmutated sequence in comparisons against samples with induced Cas9 DSBs. Using the specific guide sequence (Supplementary Table 2) and ab1 files from the uncut control and cut samples, samples were deconvoluted with TIDE. The setting used varied by guide. For sgLAD alignment window left boundary was set to 300 bp from the start of the DNA sequence, and a 150 bp (500-650 bp from the start of the DNA sequence decomposition window; for sgiLAD the alignment window left boundary was set to 100bp and the decomposition window to 240-500 bp; for sgMDC1: alignment window left boundary was 140bp, with the decomposition window 300-550bp; for sgRNF168: the alignment window left boundary was 200bp, with the decomposition window 300-550bp; for sg*LMNB1* the alignment window left boundary was 400 bp, with the decomposition window 580-700bp; and finally for sg*LBR*: alignment window left boundary 300bp, decomposition window 500-650bp. For all sgRNA samples, a 20 bp indel size range was used. To calculate the proportion of repair events, indels detected with non-significant *p*-values (*p* > 0.001) were disregarded from analysis and all significant events were first summed. The proportion of NHEJ events was the summed incidence of indels between -4 and +2. Similarly, the proportion of MMEJ events was calculated as the summed incidence of deletions greater than 5bp. The incidence of ‘0’ indels were ascribed as uncut events.

#### APEX2 and Tandem mass tag (TMT) labelling

15-20 million cells per condition were treated with Shield1 (0.5 µM, Takara 632189) for 20 h prior to labelling. APEX labelling was performed similar to a previous study^14^. Briefly, cells were incubated with Biotin-Phenol (500 µM, Iris Biotech, LS-3500) for 30 min before addition of 1 mM H_2_O_2_ for 1 min. Cells were washed three times with quencher solution (10 mM Sodium Ascorbate, 5 mM TROLOX, 10 mM Sodium Azide), harvested by scraping on ice and pelleted. To isolate nuclei, pellets were washed in PBS, resuspended in Buffer A (10 mM HEPES/KOH pH7.9, 1.5 mM MgCl_2_, 10 mM KCl) and incubated for 10 min on ice, before centrifugation. Cell pellets were resuspended in Buffer A + 0.2% NP-40, 1 mM DTT, 1 mM PMSF, EDTA-free protease inhibitors (Roche, 11836170001). Cells were homogenised (Dounce homogeniser) and the suspension centrifuged. Nuclei pellets were resuspended in Buffer A + 150 mM NaCl, 1 mM DTT, centrifuged and incubated on ice with regular agitation by vortex. The suspension was centrifuged and the pellet retained. Nuclei were resuspended at RT in 4% SDS, 100 mM Tris pH 7.5, for 3 min. 350 µl SDS-free RIPA buffer (1 mM DTT, 1% NP40, 50 mM Tris pH7.5, 150 mM NaCl, 0.5% Na-Deoxycholate, 10 mM Sodium Ascorbate, 5 mM Sodium Butyrate, EDTA-free Protease Inhibitor Cocktail, Phosphatase Inhibitor (PhosStop, Roche, 4906845001)) was added for lysis, reducing SDS concentration to <1%. The suspension was centrifuged to extract chromatin and the supernatant (SN1) was retained. Chromatin pellets were resuspended in 300 µl SDS-free RIPA and sonicated (Qsonica, 70% Amp, 10 s on 20 s off, 6.5 min). Suspensions were centrifuged at maximum speed (12,000 RPM) for 20 min. Supernatants were collected (SN2) and any pellets resuspended and incubated for 20 min at RT in 20 µl of 8 M Urea. The urea concentration was gradually diluted to 0.4M using nuclease-free H_2_O, and the suspension centrifuged, retaining supernatant (SN3). SN1, SN2 and SN3 were combined, and quantified (Bradford Plus protein assay reagent, Thermo Scientific, PI23238) to equalize protein loading for immunoprecipitation (IP). Dynabeads™ MyOne™ Streptavidin T1 (Thermo Fisher Scientific, 65601) were equilibrated in SDS-free RIPA, and equalized protein extract loaded for overnight incubation (at 4°C, rotating), retaining 0.1% total extract for input immunoblot. Following incubation, beads were washed twice with SDS-free RIPA, once with 1 M KCl, once with Na_2_CO_3_, once with 2 M Urea 10 mM Tris-HCl pH 7.5, and three times with 100 mM triethylammonium bicarbonate (TEAB, Thermo Fisher Scientific, 90114). Beads were then transferred to a new low protein binding vial (Thermo Fisher Scientific, 90410) and washed twice with 100 mM TEAB. All washes were conducted for 5 min at 4°C, with rotation. Beads were centrifuged and resuspended in 100 µl TEAB + 5 mM TCEP (Sigma Aldrich, C4706-2G) + 10 mM Auxin (IAA; Merck, 8.04744) for 30 min at RT. 10 mM DTT was added to quench IAA, and beads incubated with 5 µg Trypsin (Promega, V5280) for 18 h at RT. Peptides were transferred to a new low protein binding vial and dried (SpeedVac). Peptides were resuspended in 15 µl TEAB and pH checked to be pH8. Peptides were incubated with 5 µl TMT (Thermo Fisher Scientific, A52047) at RT for 2 h. 0.5 µl of each sample was added to 100 µl 0.1% TFA (Thermo Fisher Scientific, 400003). A preliminary Mass spectrometry (see experimental details in next section) was conducted to ensure labelling efficiency, which was confirmed to be 99.70%. Once confirmed, samples were combined and dried (SpeedVac). Pellets were resuspended in 200 µl of 0.1% TFA. Peptides were fractionated using Pierce™ High pH Reversed-Phase Peptide Fractionation Kit (Thermo Fisher Scientific, 84868), according to manufacturer’s instructions for TMT labelled peptides. Samples were resuspended in 45 µl of 0.1% formic acid and 20 µl was injected into the mass spectrometer.

#### (TMT)-based mass spectrometry (MS)

LC-MS/MS analysis was performed using a Vanquish Neo UHPLC system coupled to an Orbitrap Exploris 480 mass spectrometer equipped with a FAIMS Pro Duo interface (Thermo Fisher Scientific), operated in data-dependent acquisition (DDA) mode.

In summary, samples were loaded onto a trap column (PepMap NEO C18 5µ, 300µm x 5mm, 100 A°) at a flow rate of 20 µl/min using 0.1% formic acid in H_2_O and separated on an analytical column (Easy-Spray PepMap Neo 2 μm C18 75 μm X 500 mm, 100 A°, Thermo Scientific) with a flow rate of 350 nl/min with a linear gradient of 1 to 35% solvent B (80% AcN, 20% H_2_O, 0.1% formic acid) over a 90 min gradient. Ion source conditions were maintained at 1.75kV positive voltage, 275°C ion transfer tube temperature and standard resolution with 3.8 L/min FAIMS carrier gas flow throughout the acquisition. Two compensation voltages (CV), -45 and – 65 V with 2 and 1 sec cycle time respectively for each CV were used with RF lens of 50% in both cases. Mass spectrometric data was acquired using a DDA MS2 fragmentation, with RunStart internal EasyIC, i.e. fluoranthene (*m/z* 202.0777), as lock mass. Monoisotopic Precursor Selection MIPS (peptide mode), intensity (1.0E4), precursor fit (50% and 0.7 amu fit window), charge state (2-7) and dynamic exclusion (30 sec) mass filters were used. Details are as follows: MS^1^ Orbitrap resolution was set at 120,000 (*m/z* 200) over a mass range of *m/z* 375 to 1550 using a normalized automatic gain control (AGC) target of 300% and maximum injection time (maxIT) of 20 ms; MS^2^ Orbitrap resolution was set at 60,000 (*m/z* 200) with the first mass set at *m/z* 110, an isolation window of 0.7 amu was used, with AGC was set at 200% with 105 msec IT, fragmentation was carried out using higher-energy collisional dissociation (HCD) with a normalized collision energy (NCE) of 34.

Data was analysed using Proteome Discoverer 3.0 (Thermo Fisher Scientific) with Sequest HT and Comet as the search engines. A modified *Homo Sapiens* (TaxID=9606) FASTA file (with Streptavidin and GFP-m^6^a-Tracer-APEX2 peptide sequence), was used for database searching. For the Sequest HT engine precursor and fragment mass limits were fixed at 20 ppm and 0.02 Da accordingly and Trypsin protease was used to generate peptides. For database search, the following dynamic modifications were used: oxidation +15.995 Da (M), Deamidated / +0.984 Da (N, Q), TMTpro, +304.207 Da (K), Carbamidomethyl, +57.021 Da (C) with a Dynamic Modifications (peptide terminus) N-Terminal Modification: TMTpro, +304.207 Da (N-Terminus. The protein FDR and the peptide spectrum match were both set to 0.01%. For the Comet engine the following variable (oxidation / +15.995 Da (M), Deamidated / +0.984 Da (N, Q)) and static (TMTpro / +304.207 Da (Any N-terminus), TMTpro / +304.207 Da (K) and Carbamidomethyl / +57.021 Da (C)) modifications were selected. FDR was set to the same value (0.01) for both search engines.

#### Proteomics Data Processing and Statistical Analysis

Raw protein abundances were first median-normalized within each sample (column-wise) to account for variation in protein loading. To reduce potential batch effects, log₂ transformation and centring were performed within each biological group. Statistical analysis was conducted in *Perseus* (version 2.0.11.0^15^, where two-sample *t*-tests were applied to assess differential protein abundance between conditions. Proteins with *p* < 0.05 and absolute log₂ fold change > 0.5 were considered significantly regulated and used as input for functional enrichment analysis using Fisher’s exact test. Principal component analysis was performed in Python 3 using pandas and scikit-learn libraries. Expression values were standardized using StandardScaler from scikit-learn before dimensionality reduction with PCA. The data was visualised using matplotlib and seaborn libraries.

The mass spectrometry proteomics data have been deposited to the ProteomeXchange Consortium via the PRIDE^16^ partner repository with the dataset identifier PXD066954 and 10.6019/PXD066954

#### Western blot analysis

Cells were cultured and treated as indicated. Cells were lysed in 30 µl RIPA buffer supplemented with Protease and Phosphatase inhibitors. Protein concentration was determined using the BCA assay (Thermo Fisher Scientific, 23225). Equal volumes of protein were denatured using 4x Laemmli buffer (Thermo Scientific, J60015.AD) and heated at 95°C for 10 min. Proteins were separated by gel electrophoresis on SDS-Page gels (Invitrogen, 4-12% NuPAGE Bis-Tris) in Tris/Glycine/SDS buffer at 80-140V, and then transferred onto Nitrocellulose membranes (0.45 µm, GE Healthcare, 10600062) for 90 min at 400 mA, using ice cold Tris/Glycine buffer + 10% methanol. Membranes were blocked for >30 min in 5% milk/PBS-T at RT. Membranes were incubated with primary antibodies against pATM (Abcam, ab81292) 1:1000, ATM (Abcam, ab32420) 1:1000, CHK1 (Santa Cruz, sc8408) 1:1000, pCHK1 (Cell Siganling, 2348S) 1:500, γH2AX (Abcam, ab22551) 1:1000, VINCULIN (Sigma Aldrich, V4505) 1:1000, GAPDH (Millipore, MAB374) 1:30000 (Supplementary Table 4), overnight at 4°C with agitation. Three PBS-T washes were conducted prior to incubation for 1 h at RT with the appropriate secondary antibody: Goat Anti-Mouse IgG (H+L), HRP (Thermo Fisher Scientific, 31430) 1:5000 or Goat and-Rabbit IgG (H+L), HRP (Thermo Fisher Scientific, 31460) 1:5000 (Supplementary Table 4), diluted in 5% milk/PBS-T. Membranes were then washed three times, for 5 min each, in PBS-T. Western blots were developed using chemiluminescent reagent (Pierce ECL western blotting substrate, Thermo Fisher Scientific, 32109) and imaged using ImageQuant LAS 4000.

#### CUT&RUN

CUT&RUN was carried out using reagents from the CUT&RUN Assay Kit (Cell signalling, 86652) unless specified otherwise. 1x10^5^ RPE-1 cells per input or reaction were collected 6 hr post-transfection with guide RNA and treatment with DMSO or 10 µM ATM inhibitor. Inputs were processed following the protocol of the CUT&RUN Assay Kit and sonication was carried out using a Qsonica Q800R2 instrument at 70% amplitude for 6 min with cycles of 10 seconds on and 20 seconds off, at 4°C. The remaining cells were processed following the protocol of the CUT&RUN Assay Kit. Briefly, cells were washed and bound to Concanavalin A magnetic beads. Then, incubation with primary antibodies (Supplementary Table 4) was then carried out at 4°C overnight, using rabbit IgG isotype control (1:20, Cell signalling 66362), mouse IgG1k isotype control (1:100, Invitrogen 14-4714-82), Lamin B1 (1:50, Santa Cruz sc56144), 53BP1 (1:50, Novis Bio NB100-904) and H3K9me3 (1:100, Abcam ab8898). Beads were then washed and pAG-MNase binding was carried out for 1 hr at 4°C. Beads were then washed twice more and MNase digestion was carried out by adding calcium chloride and incubating samples for 30 min at 4°C. To release digested DNA fragments into solution, beads were incubated with 1X Stop Buffer containing 50pg spike-in DNA per reaction for 10 min at 37°C. The solution containing DNA fragments was then collected and purified using QIAquick PCR Purification Kit (Qiagen), following the manufacturer’s protocol. For analysis by qPCR, Luna qPCR master mix (NEB, M3003E) was used per the manufacturer’s instructions, and the reactions were carried out in triplicate using an AriaMx Real-time system (Agilent). Primers used for qPCR are listed in Supplementary Table 2. The data was then analysed firstly by subtracting the average Ct of each antibody condition from the average Ct of the corresponding input to calculate the ΔCt. The % enrichment compared to input was then calculated with 100 x 2^ΔCt^. To account for non-specific signals, the % of input values for the corresponding isotype controls were subtracted.

#### CUT&RUN sequencing library preparation

Libraries were generated with the NEBNext Ultra II DNA Library Prep Kit for Illumina (NEB #E7645S) according to manufacturer’s specifications and a previously published protocol^17,18^. Briefly, samples were quantified by Tapestation (Agilent, #5067-5584 and #5067-5603) and sample volumes adjusted for a minimum of 5ng input for the End Repair reaction. 25µl of sample was added to 3.5µl of NEBNext Ultra II End Prep Reaction Buffer, and 1.5µl of NEBNext Ultra II End Prep Enzyme Mix. This was mixed well and incubated at 20µl for 30 minutes and at 50°C for 60 minutes in a thermocycler. Next, 1.25µl of appropriately diluted NEBNext Adaptor for Illumina (according to input quantification and NEBNext Ultra II suggested ratios), 15µl of NEBNext Ultra II Ligation Master Mix, and 0.5µl of NEBNext Ligation Enhancer was added per sample. This was incubated at 20°C for 30 minutes before addition of 1.5µl of USER Enzyme and further incubation at 37°C for 15 minutes in a thermocycler. Samples were then purified by AMPure XP magnetic beads (Beckman Coulter, #A63882) at 1.2X volume to sample according to manufacturer’s specifications. Samples were then eluted into 19µl of 10mM Tris-HCl. 15µl was transferred to a fresh PCR tube. 2µl was used to verify the number of barcoding PCR cycles by qPCR (Ct of each sample +2) with 0.25µl 10mM universal p5 primer, 0.25µl 10mM p7 primer, 5µl Luna qPCR master mix (NEB, M3003E). 15µl of sample was then mixed with 10µl of Dual Index Primer Mix (NEB #E6440S) and 25µl of NEBNext Ultra II Q5 Master Mix. Samples were then incubated with the following program: 30 seconds at 98°C, Ct+2 cycles of 10 seconds at 98°C and 75 seconds at 65°C, then 5 minutes at 65°C, followed by a hold at 4°C. Samples were then purified by AMPure XP magnetic beads (Beckman Coulter, #A63882) at 0.9X volume according to manufacturer’s protocol. Libraries were then quantified by Tapestation (Agilent, #5067-5584 and #5067-5603).

#### ATAC-sequencing libraries

ATAC-seq was performed with the ATAC-seq kit (Active Motif, #53150) according to manufacturer’s specifications with few alterations. 2x10^5^ Cas9 expressing RPE-1 cells were collected 6 h post-transfection with sgRNA and treatment with DMSO or 10 µM ATM inhibitor. For RPE-1 cells not expressing Cas9, 2x10^5^ cells were collected 7 h post-transfection as described previously. 20µl of ATAC Spike nuclei (Active Motif, #53154) were added to each sample. Cells were washed with 100µl of ice-cold PBS and then with 100µl ice-cold ATAC Lysis Buffer. Cells were pelleted and resuspended in 50µl of the Tagmentation Master Mix and incubated at 37°C for 30 minutes. This was purified by Silica Binding Columns, and 33.5µl of eluate was used for PCR amplification and barcoding. Resulting ATAC-seq libraries were purified by SPRI beads (1.2X sample volume) and quantified by Tapestation (Agilent, #5067-5584 and #5067-5603).

#### Deep Sequencing and analysis workflow

Multiplexed library pools were sequenced on the Illumina NextSeq 1000 platform, with paired-end read lengths of 51bp using the P2 XLEAP-SBS™ Reagent Kit (100 Cycles) (Illumina, #20100987). Code optimization and scripting assistance was supported by ChatGPT (OpenAI; GPT-4, GPT-5, GPT-5.5, and GPT 5.6). All AI-assisted code was reviewed, modified where necessary, and validated by the authors before use in data analysis. Raw CUT&RUN sequencing reads were then processed with the nf-core/cutandrun pipeline^19,20^ with the following command: nextflow run nf-core/cutandrun -r 3.2.2 -profile docker --input ∼/samplesheet.csv --genome hg38 --spikein_genome R64-1-1 \ --outdir /path/to/ouput --work-dir ∼/nf_work --save_reference --save_trimmed --use_control true --max_memory ‘24.GB’ --max_cpus 8 --cores 30 -with-report report.html -with-trace trace.txt -with-timeline timeline.html -with-dag dag.png.

Raw ATAC-seq reads were processed with the nf-core ATAC-seq pipeline^19,21^ with the following command: nextflow run nf-core/atacseq -r 2.1.2 -profile docker --input ∼/atac_samplesheet.csv --genome hg38 --read_length 150 --outdir /path/to/output --work-dir ∼/nf_work --save_reference --save_trimmed --max_memory ‘24.GB’ --max_cpus 8 --cores 30 -with-report atacseq_report.html -with-trace atacseq_trace.txt -with-timeline atacseq_timeline.html -with-dag atacseq_dag.png. Where relevant, bigWig signal tracks for three biological replicates were averaged with wiggletools (version 1.2.11).

#### Domainograms

Domainogram calculations and representation of nuclear lamina detachment was adapted for CUT&RUN sequencing data from a previously described pipeline for DamID data^22^. BigWigs were generated by the nf-core/cutandrun pipeline^19,20^. Lamin B1 samples were normalised relative to their matched IgG control tracks. This was expressed as a log₂ normalised ratio calculated in 50-bp genomic bins using deepTools bigwigCompare^23^ with a pseudocount of 0.001. Log₂(Lamin B1/IgG) ratios of all samples were summarised in common 5-kb genomic bins with deepTools multiBigwigSummary^23^. The difference (delta) between sgLAD and sgCTRL transfected samples were calculated independently for each biological replicate as sgLAD - sgCTRL, using three biological replicates in DMSO treated conditions, and two biological replicates in ATMi treated conditions. These were then averaged, and a sliding mean across a range of increasing window sizes from 25 Kb to 1 Mb was calculated for sgLAD, sgCTRL, and delta tracks with 5000bp bins. For each window size, quantiles of log₂(Lamin B1/IgG) ratios were calculated for sgLAD and sgCTRL conditions independently, and windows where both conditions fell below or at the 0.3 quantile were excluded. This filtration was applied to match the DamID based pipeline^22^ to exclude low-signal genomic regions. The delta signal of retained windows was ranked independently for each window size, and the central 90% of data was masked. The lower and upper 5% were then tail-scaled from -1 to 0, and 0 to +1 respectively. Consequently, the lower and upper 5% of relative LaminB1 signal change genome-wide following DSB induction at the target LAD are displayed in blue and red respectively.

## Data Availability

Raw and processed sequencing data generated during this study is available at **<u>GSE343651</u>** (CUT&Run) and **<u>GSE343837</u>** (ATAC-seq). Open-source LAD annotations from LaminB1 DamID-seq data for RPE-hTERT cells were retrieved from the 4D Nucleome Data Portal^24^ (4DN accession 4DNEXUJUPQ4D). All custom Python and shell scripts employed for NGS, TIDE, and foci colocalisation data analysis and presentation of these data can be found on Zenodo^25^.

## Statistical analysis

Statistical analysis is included in figure legends. Unless otherwise stated, comparison tests were performed using GraphPad Prism version 11.1.0 for macOS. Analyses performed in Python (version 3.11.6) for all TIDE data and pATM/gH2AX colocalisation, were conducted using the statsmodels package (version 0.14.0^26^).

## Supplementary tables

**Supplementary Table 1:**
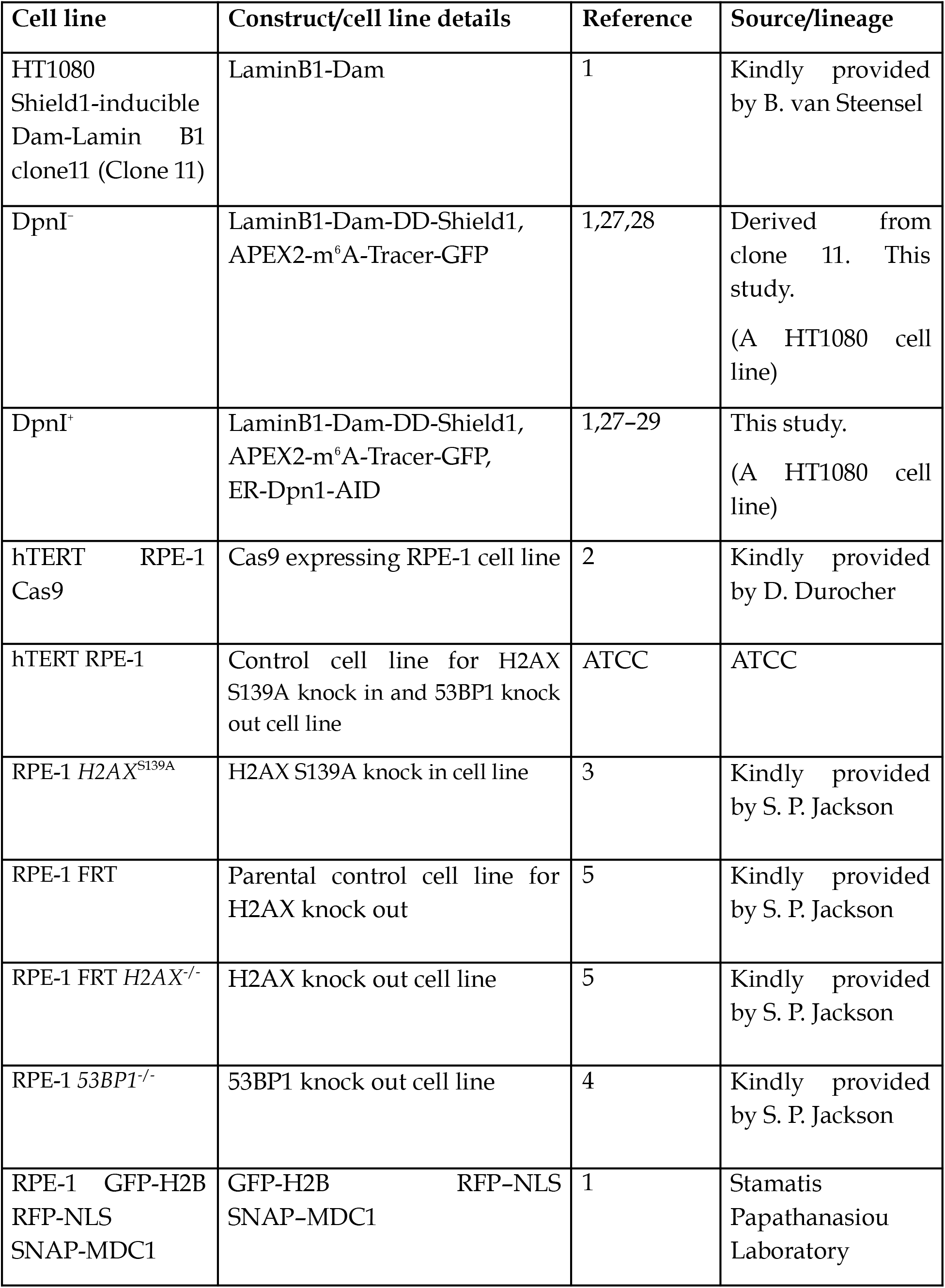
Experimental systems/cell lines used in this study.

**Supplementary Table 2:**
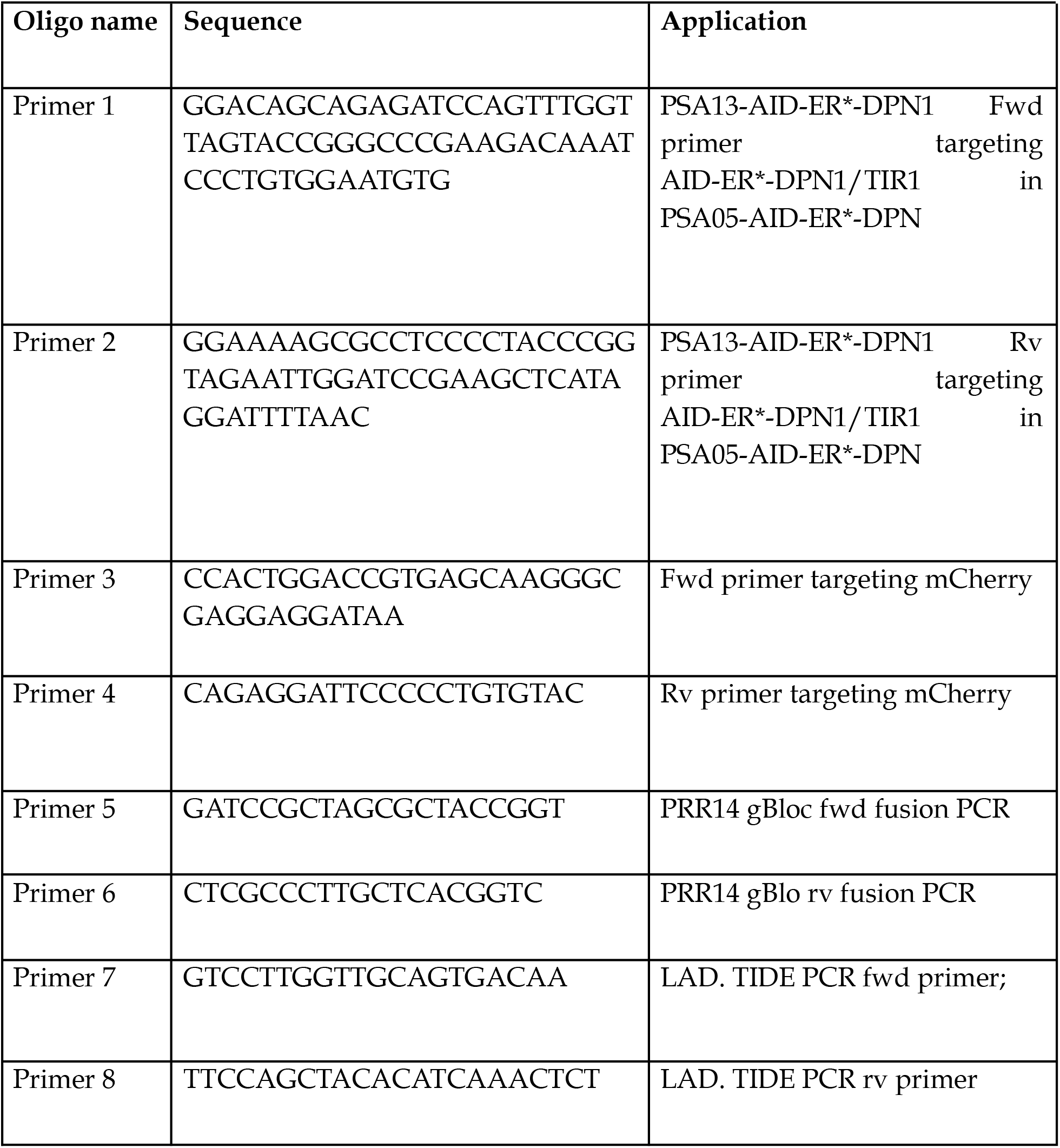
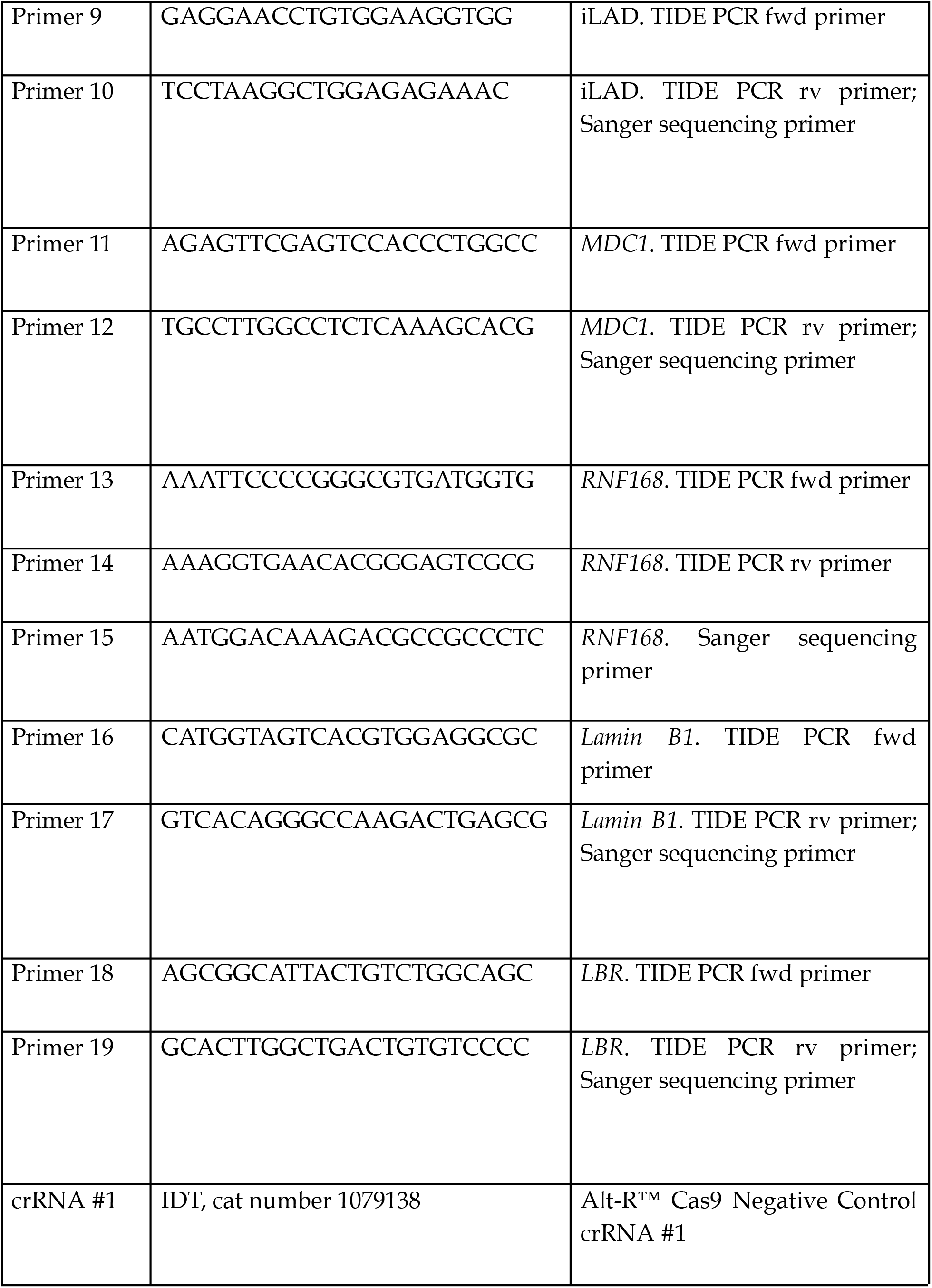
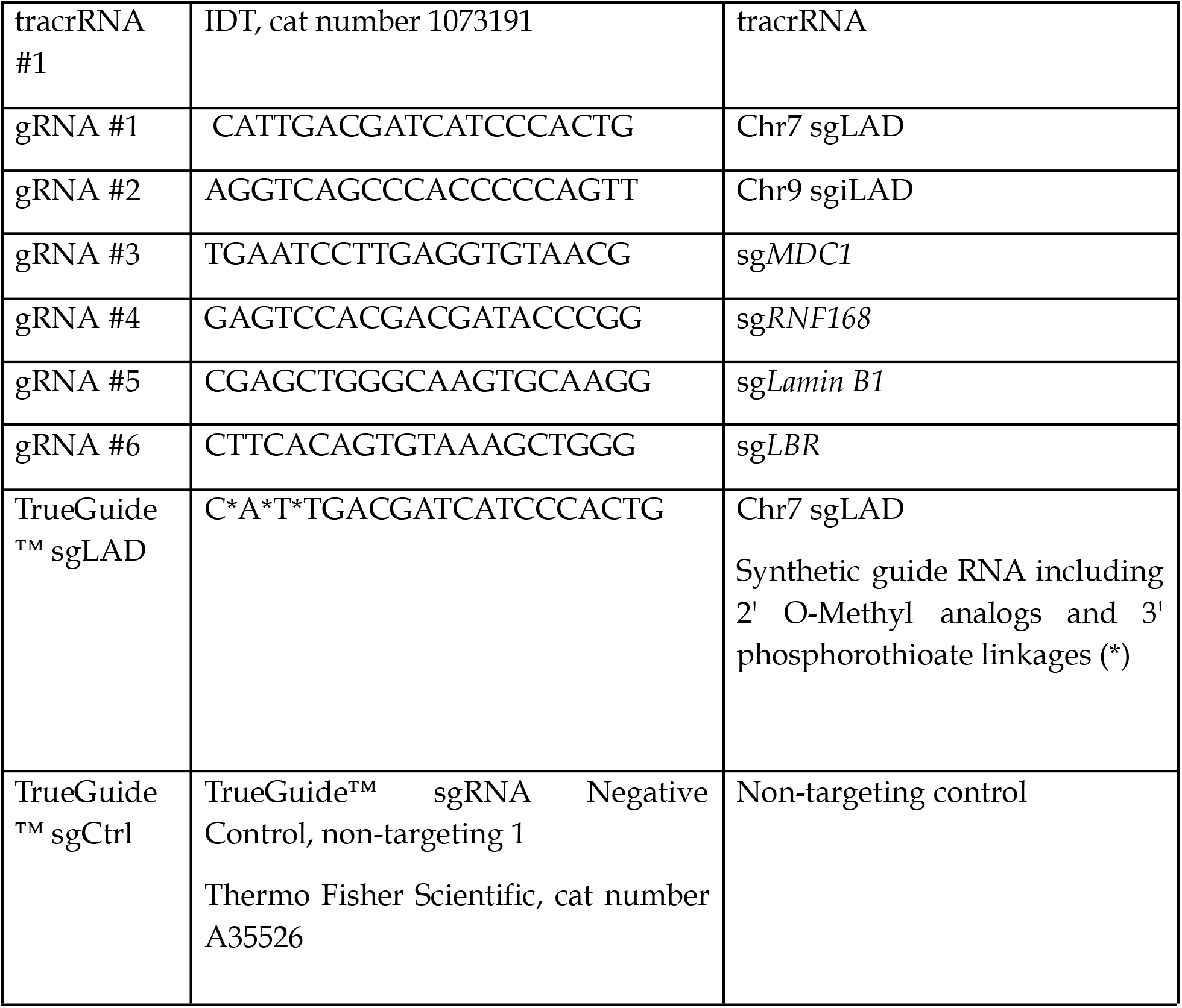
Oligonucleotides used in this study. All oligos were purchased from IDT, with the exception of the TrueGuide gRNA which was from Thermo Fisher Scientific. Fwd = forward primer. Rv = reverse primer

**Supplementary Table 3:** Chemicals used in this study.

| Chemicals | Concentration | Manufacturer | Cat no |
| --- | --- | --- | --- |
| Shield1 | 0.5 μM | Takara | 632189 |
| (Z)-4-Hydroxytamoxifen (4OHT) | 300 nM | Sigma Aldrich | H7904 |
| IAA | 10 mM | Merck | 8.04744 |
| KU-55933 (ATMi) | 10μM | Sigma Aldrich | Sml1109 |
| G418 | 25 µg/ml | Sigma Aldrich | A1720 |
| Puromycin | 0.5 µg/ml | Sigma Aldrich | P9620 |
| Ramapycin | 200 nM | Sigma Aldrich | 553210 |
| NU7441 (KU-5778) (DNAPKi) | 10 µM | Selleckchem | S2638 |
| ART558 (PolQi) | 10 µM | Artios | gift |
| B02 (RAD51i) | 20 µM | Sigma Aldrich | SML0364 |
| Demecolcine | 0.2 µg/ml | Sigma Aldrich | D1925 |
| VE-821 (ATRi) | 5 µM | Sigma Aldrich | SML1415 |
| (S)-(+)-Camptothecin (CPT) | 25 µM | Sigma Aldrich | C9911 |
| Vinblastine | 100 nM | Sigma Aldrich | V1377 |
| Latrunculin B | 100 nM | Abcam | ab144291 |
| Blebbistatine | 10 µM | Sigma Aldrich | 203390 |
| Nocodazole | 50 µM | Sigma Aldrich | SML1665 |
| BrdU | 10 µM | Sigma Aldrich | B5002 |
| TAK-243 (UBA1 inhibitor) | 5 µM | MedChemExpress | MNL7243 |
| CK666 (Arp2/3 complex inhibitor) | 100 µM | Sigma Aldrich | SML006 |
| CK689 | 100 µM | Sigma Aldrich | 182517 |

**Supplementary Table 4:**
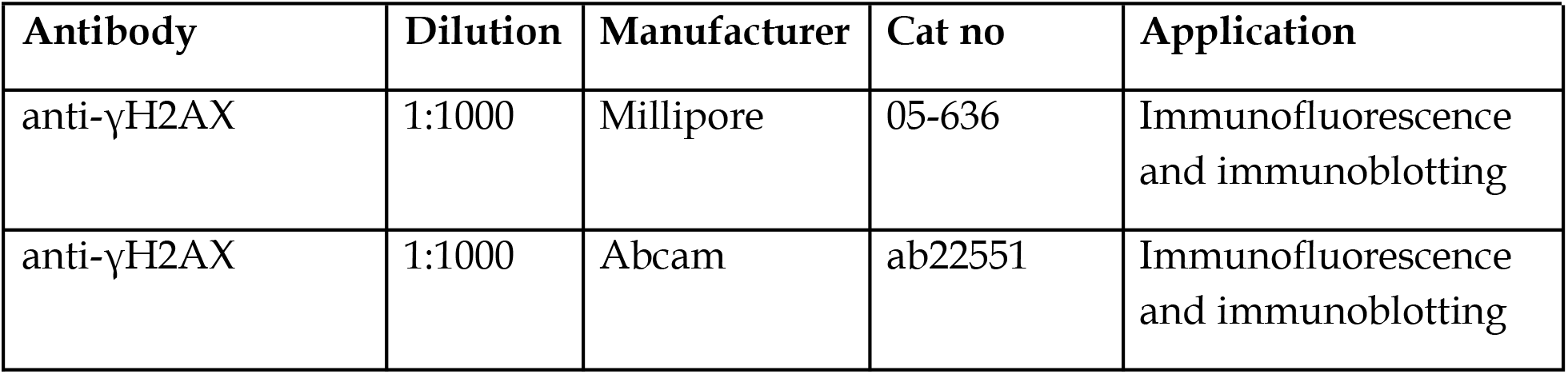
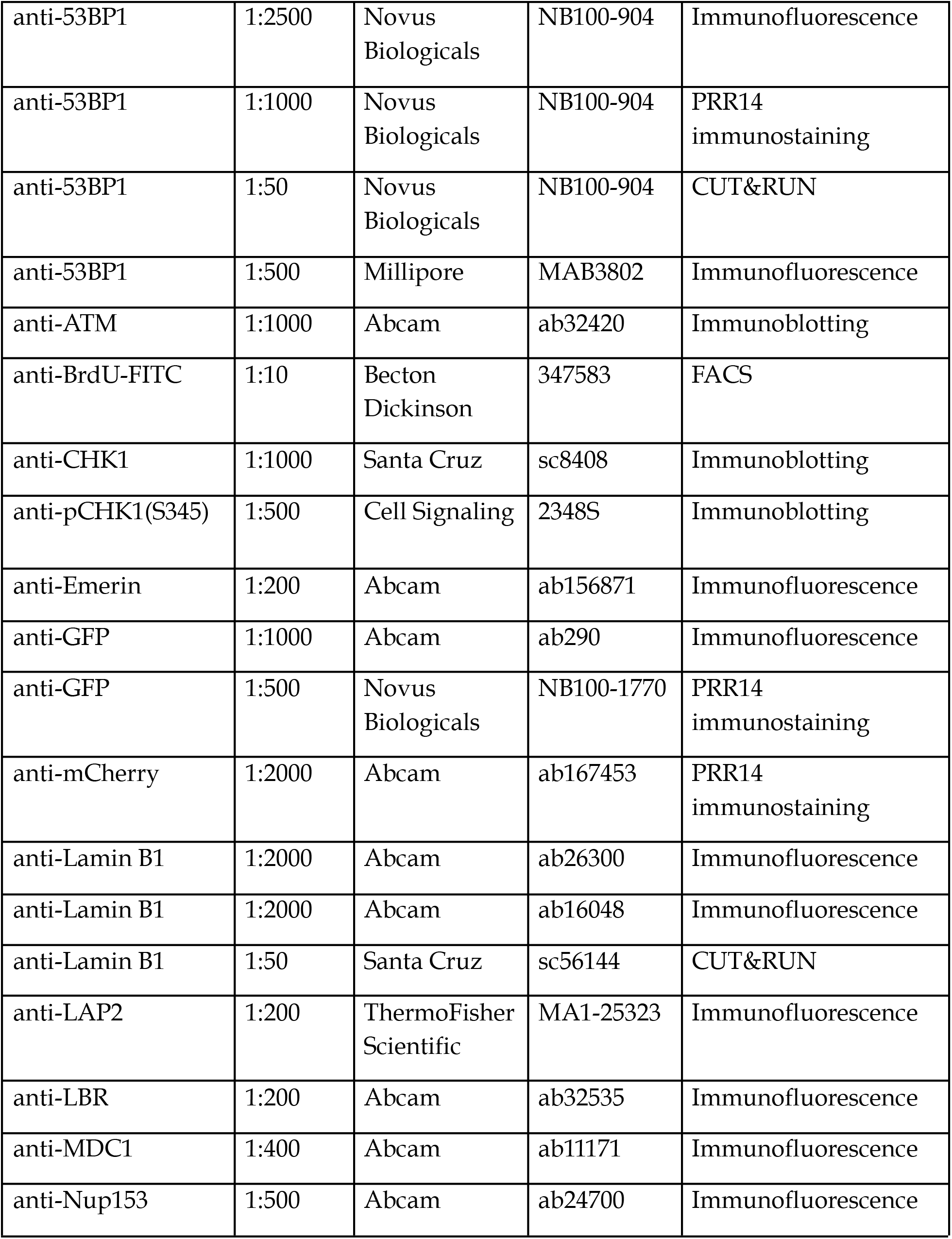
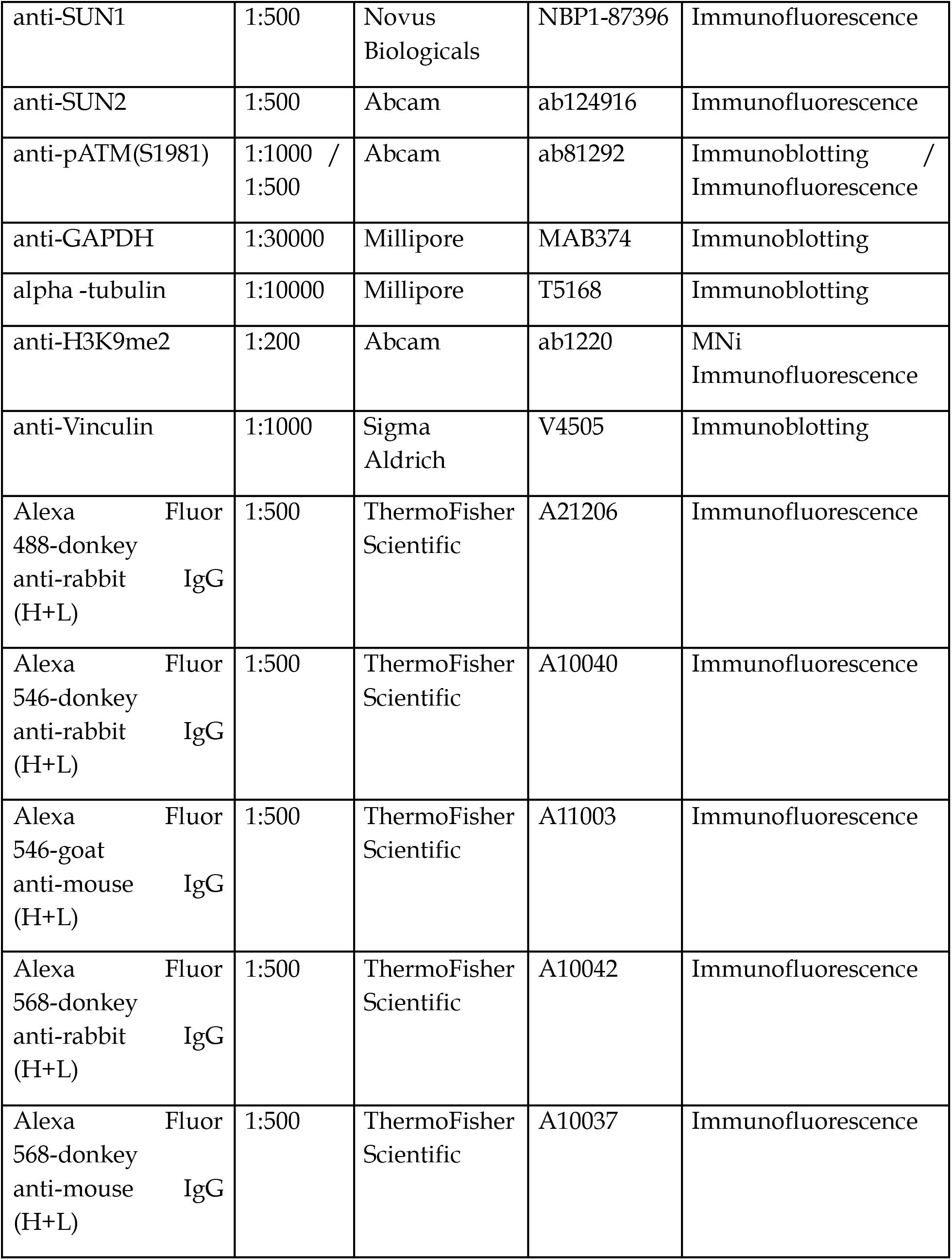
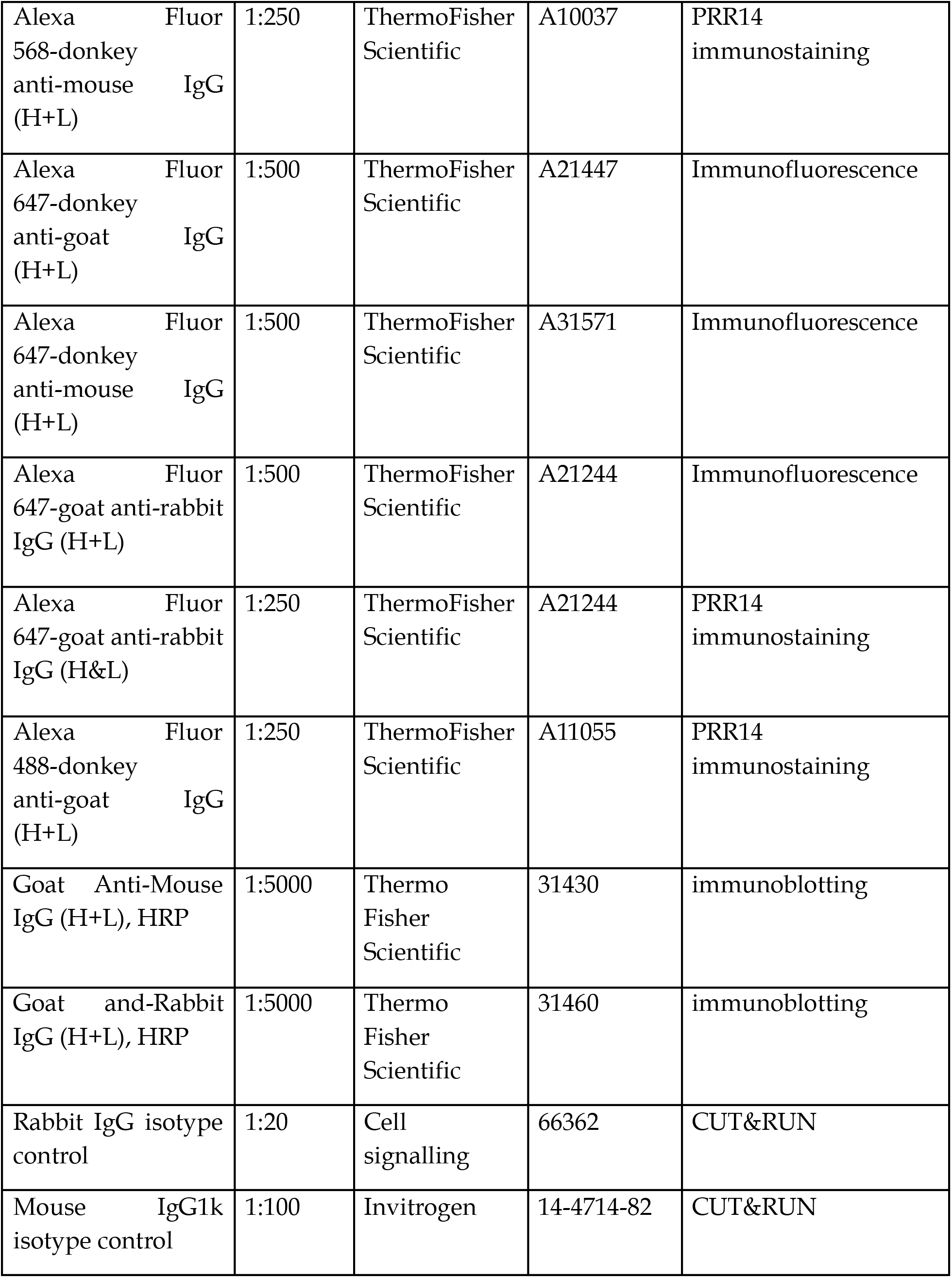
Antibodies used in this study with their respective applications and dilutions.

## Notes

https://www.ncbi.nlm.nih.gov/geo/query/acc.cgi?acc=GSE343651

https://www.ncbi.nlm.nih.gov/geo/query/acc.cgi?acc=GSE343837

### Competing Interest Statement

The authors have declared no competing interest.

